# Regulation of adaptive growth decisions via phosphorylation of the TRAPPII complex in Arabidopsis

**DOI:** 10.1101/2023.04.24.537966

**Authors:** Christian Wiese, Miriam Abele, Benjamin Al, Melina Altmann, Alexander Steiner, Nils Kalbfuß, Alexander Strohmayr, Raksha Ravikumar, Chan Ho Park, Barbara Brunschweiger, Chen Meng, Eva Facher, David W. Ehrhardt, Pascal Falter-Braun, Zhi-Yong Wang, Christina Ludwig, Farhah F. Assaad

## Abstract

Plants often adapt to adverse or stress conditions via differential growth. The trans-Golgi Network (TGN) has been implicated in stress responses, but it is not clear in what capacity it mediates adaptive growth decisions. In this study, we assess the role of the TGN in stress responses by exploring the interactome of the Transport Protein Particle II (TRAPPII) complex, required for TGN structure and function. We identified physical and genetic interactions between TRAPPII and shaggy-like kinases (GSK3/AtSKs). Kinase assays and pharmacological inhibition provided *in vitro* and *in vivo* evidence that AtSKs target the TRAPPII-specific subunit AtTRS120/TRAPPC9. GSK3/AtSK phosphorylation sites in AtTRS120/TRAPPC9 were mutated, and the resulting AtTRS120 phosphovariants subjected to a variety of single and multiple stress conditions *in planta*. The non-phosphorylatable TRS120 mutant exhibited enhanced adaptation to multiple stress conditions and to osmotic stress whereas the phosphomimetic version was less resilient. Higher order inducible *trappii atsk* mutants had a synthetically enhanced defect in root gravitropism. Our results suggest that the TRAPPII phosphostatus mediates adaptive responses to abiotic cues. AtSKs are multifunctional kinases that integrate a broad range of signals. Similarly, the TRAPPII interactome is vast and considerably enriched in signaling components. An AtSK-TRAPPII interaction would integrate all levels of cellular organization and instruct the TGN, a central and highly discriminate cellular hub, as to how to mobilize and allocate resources to optimize growth and survival under limiting or adverse conditions.

## INTRODUCTION

Plant responses to environmental stimuli involve diverse forms of growth or movement. The slow, cyclic movement of shoots and stems helps climbing plants such as vines find supportive structures, and stem growth or tendrils are then used to wrap around and cling to such structures (Darwin, 1880). Leaf movements linked to the circadian rhythm enable plants to maximize their exposure to sunlight (McClung, 2006). Tropisms are additional examples of movement in plants: the shoot bends towards directional light, whereas the root bends away from the light. In addition to phototropism, plants have tropic responses to a range of stimuli including moisture, fluctuations in temperature, gravity, touch and other mechanical cues (Garzón & Keijzer, 2011). Darwin argues that the movements of plants are driven by growth and the need to access resources, such as sunlight and water (Darwin, 1880). Gyrations, revolutions and tropisms require some form of differential growth or bending at the organ level. Bending is achieved when one side of an organ grows more rapidly than the opposing side. This differential growth is a result, at least in part, of the differential sorting of PIN-FORMED (PIN) auxin transporters, resulting in the unequal distribution of auxin, a morphogen, at opposing sides of a cell (Friml *et al*, 2002; Ding *et al*, 2011). It follows that differential growth responses such as bending require differential sorting decisions. How environmental stimuli are translated into sorting decisions remains largely unclear.

The sorting of PIN transporters has been shown to require trans-Golgi network (TGN) function. Indeed, disruption of TGN function by mutation results in the ectopic distribution of PIN or AUX proteins (Naramoto *et al*, 2014; Qi *et al*, 2011; Rybak *et al*, 2014; Ravikumar *et al*, 2018). The TGN plays a key role not only in the sorting of macromolecules, but also in exocytosis and endocytosis. In addition, the TGN performs specialized functions such as cytokinesis, cell differentiation, the establishment of cell polarity, and anisotropic growth (Gendre *et al*, 2015; Ravikumar *et al*, 2017). As an early endosome, the plant TGN is a central hub in the flow of information to and from the plant cell surface (Uemura, 2016). The plant TGN has been implicated in responses to abiotic stimuli such as drought, heat, salt stress and osmotic stress and to biotic stimuli such as fungal attack (Rosquete & Drakakaki, 2018). Studies on the role of the TGN in stress responses have been carried out predominantly with core trafficking components such as Rab GTPases, tethering factors, and Q-SNAREs, required for membrane tethering, docking and fusion (Rosquete & Drakakaki, 2018; Ravikumar *et al*, 2017). Trafficking mutants typically exhibit root growth defects and/or hypersensitivity to abiotic cues such as salt stress, osmotic stress, drought or heat (Asaoka *et al*, 2013; Kim & Bassham, 2011; Lee *et al*, 2006; Rosquete *et al*, 2019; Uemura *et al*, 2012; Wang *et al*, 2011; Zhu *et al*, 2002). However, whether trafficking mutants have primary defects in growth with secondary consequences in stress responses, or whether the primary defects lie in an impaired response to stress factors remains unclear. More broadly, in the context of stress responses the question pertains as to whether the TGN is involved in decision-making processes *per se*, or merely in the execution of adaptive growth decisions.

As regards decision-making processes, there is a growing body of evidence to suggest that plants have the ability to learn, to process information, to communicate, to reach decisions, and in general to exhibit behavior that could be considered cognitive (reviewed in Severino, 2021). Severino (2021) makes the case for experimental approaches to study the decisions plant reach in complex environments. A recent experimental approach for the study of decision-making processes in germinating seedlings has incorporated two tools used in decision theory: the use of a limited budget and of conflict-of interest scenarios (Kalbfuß *et al*, 2022). A limited budget was achieved by germination in the dark in the absence of a carbon source, such that the only available energy source is that available in the seed (Kalbfuß *et al*, 2022). A conflict-of-interest scenario comprises the simultaneous withdrawal of light, which promotes hypocotyl elongation, and water, which promotes root elongation (Kalbfuß *et al*, 2022). As the severity of water stress increased, root length increased while hypocotyl length decreased; importantly, the total seedling length remained constant (Kalbfuß *et al*, 2022). Thus, trade-offs in hypocotyl versus root growth were observed and these comprise a binary readout for responses to these additive stress conditions. Decision mutants were defined as mutants that were either incapable of adjusting their hypocotyl/root ratios in response to additive stress, or that consistently reached the wrong growth decisions as compared to the wild type (Kalbfuß *et al*, 2022). By emphasizing growth trade-offs, the experimental approach developed by Kalbfuß *et al* (2022) is aligned with the definition of decision-making as entailing an appraisal of the advantages and disadvantages of various courses of action (Karban & Orrock, 2018). While Kalbfuß *et al* (2022) address decision-making at a cellular level, the literature on plant decision-making has, to our knowledge, not included considerations about the possible role of the TGN.

To understand the role of the TGN in adaptive or stress responses, it would be important to deploy a battery of gene products not only broadly associated with or localized to the TGN, but also intrinsic to TGN structure and function. Two such proteins or complexes are ECHIDNA and the Transport Protein Particle II (TRAPPII) complex. ECHIDNA was identified as an upregulated transcript in elongating cells (Gendre *et al*, 2011). Yeast and metazoan TRAPPII is a hetero-oligomeric complex that acts as a Guanine nucleotide Exchange Factor (GEF) for Rab GTPases, converting GDP-bound inactive Rab GTPases to active GTP-bound forms (Cai *et al*, 2005; Morozova *et al*, 2006; Pinar *et al*, 2015; Thomas & Fromme, 2016; Riedel *et al*, 2018). TRAPPII has been shown to play a key role in the regulation of the TGN in all eukaryotes, but our understanding of its potential physiological roles is incomplete (Pinar & Peñalva, 2020). The Arabidopsis TRAPPII (AtTRAPPII) complex was identified by mutation in screens for seedlings with aberrant morphogenesis or cytokinesis defects (Söllner *et al*, 2002; Thellmann *et al*, 2010; Jaber *et al*, 2010). AtTRAPPII most resembles fungal and metazoan TRAPPII complexes, with the exception of one plant-specific subunit (Garcia *et al*, 2020; Kalde *et al*, 2019; Pinar *et al*, 2019). We have previously shown that ECHIDNA and TRAPPII have overlapping yet distinct functions at the TGN in Arabidopsis (Ravikumar *et al*, 2018). ECHIDNA is primarily required for the genesis of secretory vesicles and, as a consequence, for cell expansion (Boutté *et al*, 2013; Gendre *et al*, 2013; McFarlane *et al*, 2013). AtTRAPPII plays a role not only in basal TGN functions – exocytosis, endocytosis, protein sorting – but also in more specialized TGN functions such as cytokinesis and the establishment of cell polarity (Ravikumar *et al*, 2018). Whether or not AtTRAPPII plays a role in responses to abiotic cues such as osmotic or drought stress remains to be determined.

In this study, we focus on the TRAPPII complex as a starting point as it is required for all aspects of TGN function, including the sorting of proteins such as PINs to distinct membrane domains (Qi *et al*, 2011; Rybak *et al*, 2014; Ravikumar *et al*, 2018). We first explored the TRAPPII interactome and also surveyed dynamic or conditional interactions. Together with yeast two-hybrid screens, this identified shaggy-like kinases such as AtSK21/BIN2 as TRAPPII interactors. We corroborated this finding with *in vitro* kinases assays and pharmacological inhibition *in vivo*. Shaggy-like kinases are multi-taskers that integrate a vast number of biotic and abiotic cues (Lv & Li, 2020; Planas-Riverola *et al*, 2019; Youn & Kim, 2015; Li *et al*, 2021; Song *et al*, 2023). AtSK21/BIN2 has recently been implicated in decision-making in the Arabidopsis seedling (Kalbfuß *et al*, 2022). We explore the meaning of the AtSK-TRAPPII interaction using a variety of assays to monitor stress responses and differential growth decisions.

## RESULTS

### The TRAPPII interactome contains a large number of signaling components

To gain insight into TGN function, we used the TRAPPII interactome as a starting point. We have previously identified TRAPP and EXOCYST subunits, cytoskeletal proteins and Rab GTPases in the TRAPPII interactome (Rybak *et al*, 2014; Steiner *et al*, 2016; Kalde *et al*, 2019; Garcia *et al*, 2020). However, no global meta-analysis of the vast Arabidopsis TRAPPII interactome has been carried out to date. We, therefore, performed a gene ontology (GO) term enrichment analysis. AtTRAPPII consists of seven shared core subunits and three TRAPPII-specific subunits (AtTRS120/TRAPPC9, CLUB/AtTRS130/TRAPPC10 and TRIPP; Garcia *et al*, 2020; Kalde *et al*, 2019). We focused on the TRAPPII-specific CLUB:GFP interactome. This interactome was significantly enriched (fold-enrichment ≥ 4; adjusted *P*-value ≤ 0.003) in proteins involved in cell division and in trafficking or transport (Fig. 1A; Fig. S1A), which is consistent with known cytokinesis, trafficking, sorting and TGN defects in *trappii* mutants (Jaber *et al*, 2010; Qi *et al*, 2011; Rybak *et al*, 2014; Ravikumar *et al*, 2018). GO terms describing root hair elongation and microtubule organization were also scored (Fig. 1A), and this is equally unsurprising in light of *trappii* defects in root hair tip growth and microtuble organization (Jaber *et al*, 2010; Söllner *et al*, 2002; Thellmann *et al*, 2010; Steiner *et al*, 2016). Interestingly, a large number of significantly enriched GO categories describe responses to stimuli such as light, metal ions or sugar (grouped under “signaling”, “response to stimulus”, “light responses” or “photosynthesis” in Fig. 1A and Fig. S1A). The large number of enriched GO terms implicated in signaling suggests that AtTRAPPII may act as a cellular hub.

**Figure 1.**
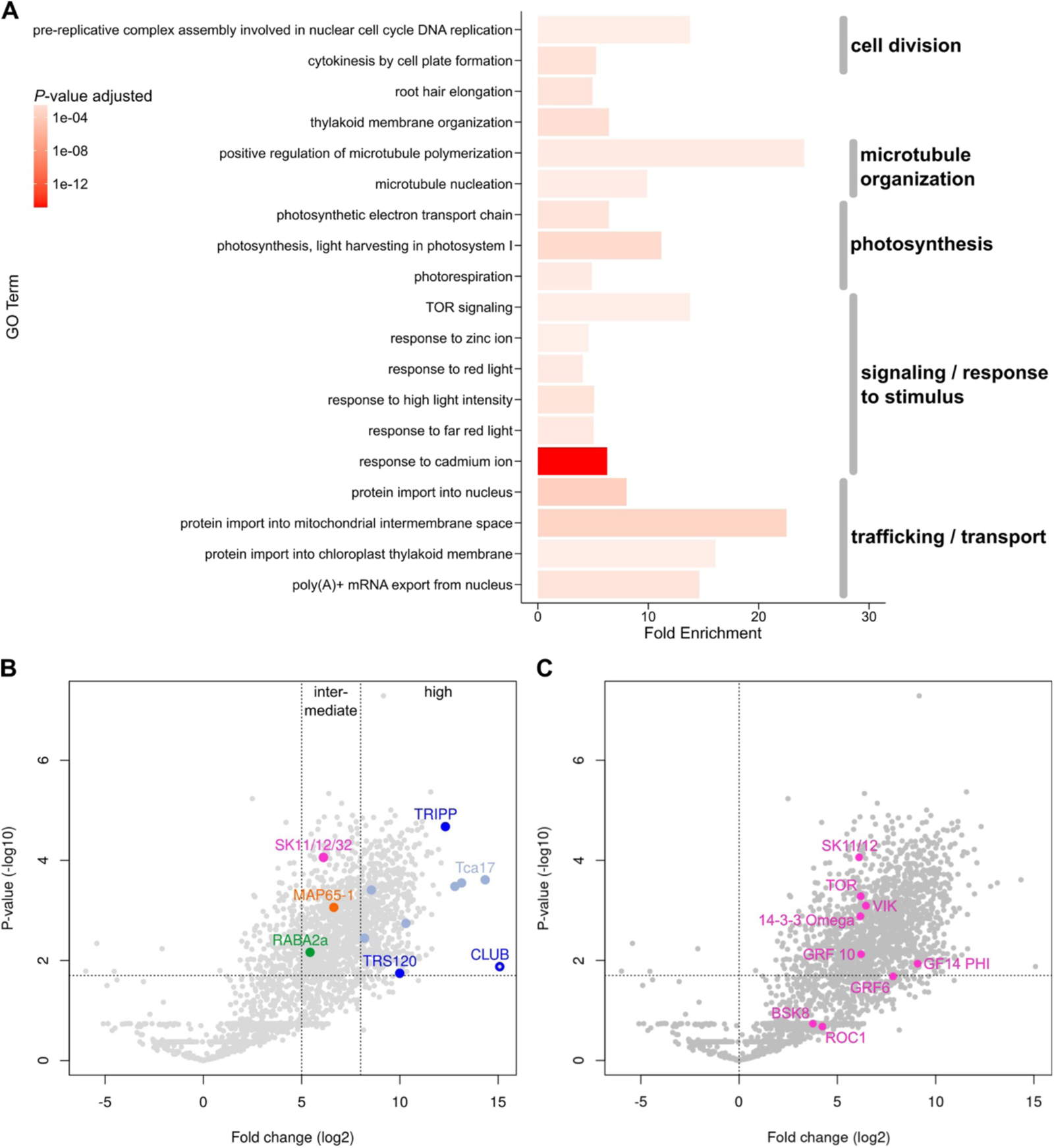
The TRAPPII interactome. The data are derived from an analysis of IP-MS from inflorescences with the TRAPPII-specific subunit CLUB:GFP as bait (see Fig. 1B; Kalde *et al*, 2019). Each protein was present in all three biological replicates. The soluble GFP empty vector was used as a negative control. **A.** Gene ontology (GO) term enrichment analysis of the TRAPPII interactome. Depicted are highly enriched (fold enrichment ≥ 4) and significant (FDR-adjusted *P*-value ≤ 0.003) GO term associations of biological processes of level 0 as bar plots. The length of each bar depicts the fold enrichment of GO terms associated with detected proteins, while the color intensity indicates the significance given as the *P*-value adjusted for the false discovery rate (FDR). Interactors of intermediate intensity (> 5 and < 8) were used (see Fig. S1A for an analysis of high confidence interactors). **B, C.** Volcano plots are presented. On the X axis: The ratio (or fold-change) was calculated for each protein as the average intensity of the signal in the experiment divided by its average intensity in the control. On the Y axis the *P*-values of the signal in the experiment versus the control, depicted along a negative log10 scale, are shown. Dotted grey lines represent cutoffs: *P*-value ≤ 0.02 and Ratio > 8 for high fold-change or > 5 for intermediate fold-change for panel B and > 1 for panel C. **B.** Note that TRAPPII subunits (light blue for core TRAPP; dark blue for TRAPPII-specific subunits; the CLUB/AtTRS130:GFP TRAPPII-specific bait is depicted as an open circle) are in the upper right field, indicating high abundance and good reproducibility. AtSKs (magenta), MAP65-1 (orange) and RAB-A2a GTPase (green) are all in the upper middle field; these may be transient interactors of TRAPPII. Note that AtSKs (AtSK11/12/32) are more significant than validated interactors such as MAP65 and RAB-A2a (Kalde *et al*, 2019; Steiner *et al*, 2016). **C.** Highlighted in magenta are members of the brassinosteroid signaling pathway that were differentially enriched over light-versus dark-grown seedlings in a different IP-MS experiment. Note that BSK8, and ROC1 did not meet the significance cutoffs. The most significant interactors were GSK3/AtSK shaggy-like kinases and TOR. Related to Fig. S1, Table S2.

To further explore a possible implication of the TRAPPII complex in signaling, we surveyed the dynamic interactome, by which we refer to interactors perceived under one environmental condition (for example in the light) but not in another (for example in the dark). By these criteria, brassinosteroid (BR) signaling components were significantly enriched (P = 0.016); this category encompasses 11 proteins, of which 9 were detected in CLUB:GFP immunoprecipitates from light-grown influorescences (Fig. 1B-C; Fig. S1B; Table S2). Among these were the TOR kinase and a family of shaggy-like kinases (AtSKs; Fig. 1C; Fig. S1B; Table S2). TOR signaling was also a significantly enriched GO term in the TRAPPII interactome (Fig. 1A). TOR and AtSKs are highly significant interactors in the TRAPPII-specific subunit CLUB:GFP interactome (Fig. 1B-C; Fig. S1B). They show a fold change that is lower than seen for components of the TRAPPII complex, but similar to that seen for validated interactors such as RAB-A2a and MAP65, expected to form more transient associations (Fig. 1B-C; Kalde *et al*, 2019; Steiner *et al*, 2016). We then used yeast two-hybrid (Y2H) to probe for binary interactions between TRAPPII and signaling components identified in the IP-MS. Y2H was carried out with TRAPPII subunits and truncations thereof (Fig. 2A; Kalde *et al*, 2019; Steiner *et al*, 2016; Garcia *et al*, 2020). In a large scale Y2H screen including 2400 pair-wise tests, an interaction was detected between a TRS120^499-1187^ truncation (TRS120-T2) and the shaggy-like kinase BIN2 (AtSK21; Fig. 2B). BIN2 interacted specifically with AtTRS120 and not with other tested TRAPPII subunits (Fig. 2B). Furthermore, we did not detect any other TRAPPII-kinase interactions in our pairwise Y2H assays. In conclusion, mass spectrometry and Y2H identify physical interactions between TRAPPII and AtSKs, *in planta* and in a heterologous system.

**Figure 2.**
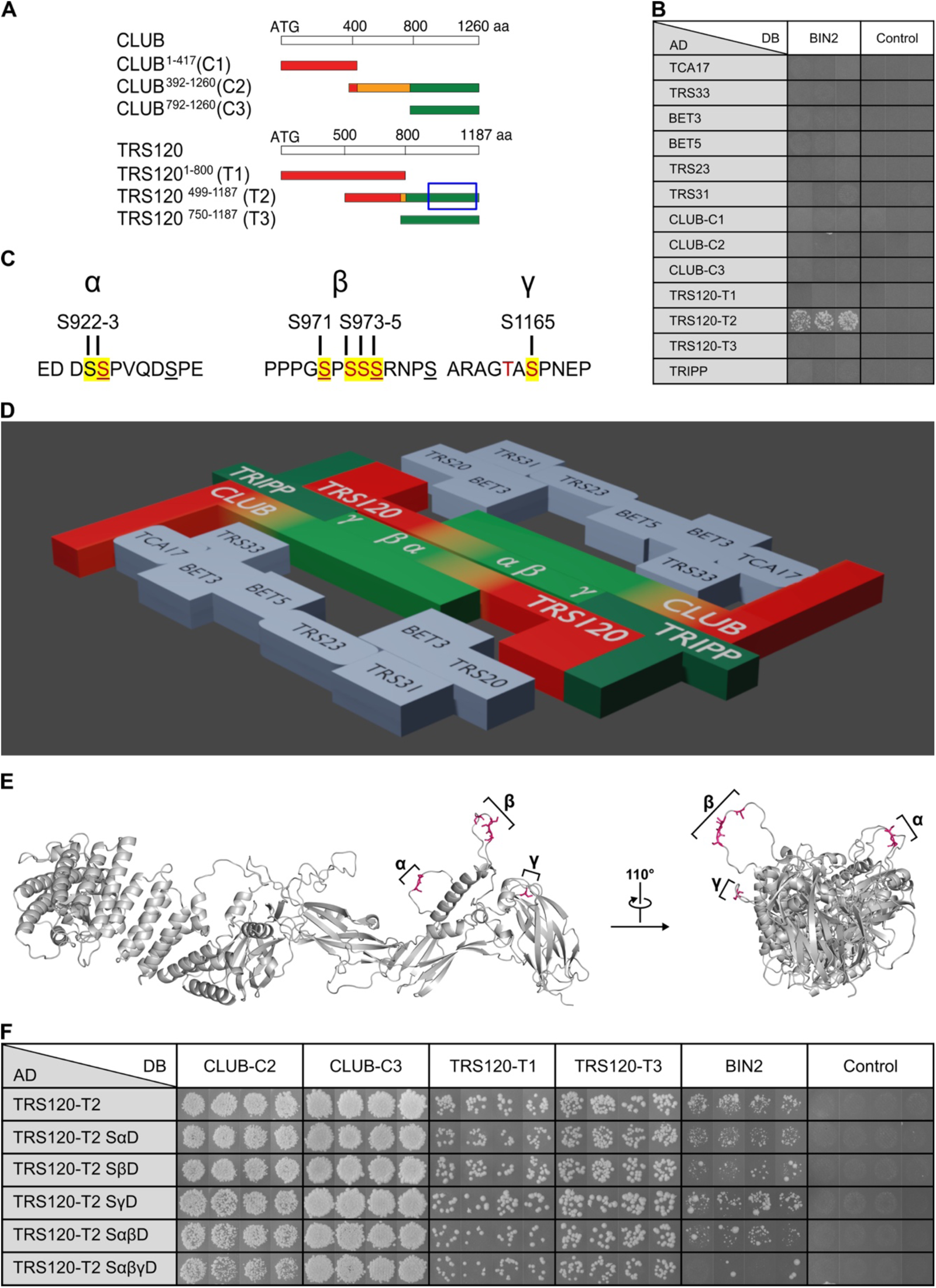
GSK3/AtSK sites in TRAPPII and binary interactions. Yeast two-hybrid assays were carried out by pairwise one-on-one mating in four independent replicate experiments, of which three (B) or four (F) are shown. As negative control, the respective AD constructs were tested with the empty DB vector. The panels are from different plates (in B, F). **A.** TRAPPII-specific truncations for yeast two-hybrid were based on phylogenetic analysis (Steiner *et al*, 2016). Conserved sequences are depicted in red, intermediate degrees of conservation in orange, and plant-specific sequences are in green. All three GSK3 sites found to be phosphorylated *in vivo* reside in the plant-specific moiety of TRS120-T2 (green; a blue rectangle delineates the region of interest). **B.** Yeast two-hybrid assays of interactions between BIN2 and TRAPPII subunits. BIN2 was fused to the GAL4 DNA-binding domain (DB) and TRAPPII subunits and truncations thereof (CLUB-C1, -C2, -C3 and TRS120-T1, -T2, -T3) to the GAL4 activation domain (AD). The results show interactions between BIN2 and AtTRS120-T2 wild type. BIN2 bound specifically to the TRAPPII subunit TRS120 and not with other TRAPPII subunits or truncations. In a total of > 2400 pairwise tests, BIN2 was the only kinase we found that interacted with a TRAPP subunit. The Y2H did not detect CLUB/AtTRS130-BIN2 interaction shown in Fig. 1B, which shows co-purified proteins that can include indirect -as opposed to binary – interactors. **C.** GSK3 sites in AtTRS120-T2. The α-phosphosite of AtTRS120 spans amino acid positions S922 and S923, the β-phosphosite amino acids S971-S975 and the γ-phosphosite amino acid residue S1165. The canonical GSK3 consensus sequence is: (<u>pS/pT</u>)*XXX*(<u>S/T</u>), and the β-phosphosite fits this definition (see underlined <u>S</u>XXX<u>S)</u>. Even though they are annotated as GSK3 sites in the PPSP (https://www.phosphosite.org) database, the α-phosphosite deviates somewhat with <u>S</u>XXXX<u>S</u>, and the γ-phosphosite deviates completely. Amino acids in red were mutated to A or D via site-directed mutagenesis. Amino acids highlighted in yellow were found to be phosphorylated *in vivo* (see Fig. S2) and these were all serines. **D.** The Arabidopsis TRAPPII complex consists of seven shared core subunits (TCA17, TRS33, BET3, BET5, TRS23, TRS31, TRS20; light blue) and three TRAPPII-specific subunits (CLUB/AtTRS130, AtTRS120 and the plant-specific subunit TRIPP), and forms a dimer with plant-specific domains (green) at the predicted dimer interface (Kalde *et al*, 2019; Garcia *et al*, 2020). This model is based on extensive pair-wise yeast two-hybrid analysis between TRAPPII subunits in *Arabidopsis* (Kalde *et al*, 2019; Garcia *et al*, 2020). **E.** Mapping of GSK3/AtSK sites on an AlphaFold structural prediction (Jumper *et al*, 2021; Varadi *et al*, 2022) of AtTRS120. A lateral view and a frontal perspective are shown. Note that all three GSK3/AtSK sites (magenta sticks) reside in unstructured, flexible and accessible regions of the protein. **F.** Interactions between BIN2 and TRAPPII complex subunits (CLUB-C2, CLUB-C3, TRS120-T1, TRS120-T3 truncations), as positive controls, fused to the GAL4 DNA-binding domain (DB) and TRS120-T2 truncation and its phosphomutants TRS120-T2 SαD, SβD, SγD, SαβD and SαβγD fused to the GAL4 activation domain (AD). Note the positive interactions between BIN2 and AtTRS120-T2 wild type and phosphovariants; there was, however, no reproducible interaction if all three target sites were phosphomimetic (TRS120-T2 SαβγD). Related to Fig. S2-S4.

### The TRAPPII complex is a target of shaggy-like kinases

To assess whether the TRAPPII complex is a target of AtSK/GSK3 kinases, we first looked for the presence of phosphopeptides in co-immunoprecipitates. To this end, we immunoprecipitated two TRAPPII-specific subunits, AtTRS120 and CLUB/AtTRS130, and looked for phosphopeptides via mass spectrometry. This provided ample *in vivo* evidence for TRAPPII AtTRS120 phosphorylation at AtSK/GSK3 sites (Fig. S2). Furthermore, shaggy-like kinases were detected in IP-MS not only with the TRAPPII-specific subunit CLUB/AtTRS130 (Fig. 1B-C; Fig. S1B) but also with the TRAPPII-specific subunit AtTRS120 (Fig. S3). The Arabidopsis genome encodes ten shaggy-like kinases (AtSKs), which are classified into four clades (Fig. S3A). Razor peptides covering all four clades were found in the AtTRS120:GFP interactome (Fig. S3B-D).

The TRS120-T2 truncation contains the GSK3 sites we had found to be phosphorylated *in vivo* (Fig. S2; see yellow highlights in Fig. 2C). Our nomenclature for these sites is α, β and γ, which is short for TRS120-S922:S923 (α), TRS120-S971:S973:S974:S975 (β) and TRS120-S1165 (γ; Fig. 2C). These reside in the plant-specific moiety of AtTRS120 and are embedded in plant-specific sequences (see green shading in Fig. 2A; Fig. 2D) at the dimer interface inferred from extensive pair-wise Y2H tests in Arabidopsis (Kalde *et al*, 2019; Garcia *et al*, 2020) and consistent with cryo-electron micrograph (cryo-EM) structures of yeast and metazoan TRAPPII (Galindo *et al*, 2021; Mi *et al*, 2022; Bagde & Fromme, 2022). AlphaFold structural predictions (Varadi *et al*, 2022; Jumper *et al*, 2021) show that the three sites reside in unstructured, flexible and accessible regions of the AtTRS120 protein, as is observed in the majority of modified amino acid residues (Fig. 2E; Jiménez *et al*, 2007). Furthermore, a cross-kingdom structural alignment of AtTRS120 and CLUB/AtTRS130 with cryo-EM-generated structures of yeast TRAPPII (Mi *et al*, 2022) showed that the β and γ phosphorylation sites face the active site chamber or Rab GTPase binding pocket predicted by Mi *et al* (2022) and Bagde & Fromme (2022) (Fig. S4). To study the three sites, we generated site directed mutations in TRS120-T2, mutating the serine (S) and threonine (T) residues (depicted in red in Fig. 2C) to non-phosphorylatable alanine (A) residues, or to aspartate (D) to mimic constitutive phosphorylation. In the case of the TRS120-Sβ site, for example, we designate these variants as TRS120-SβA or TRS120-SβD (Fig. 2F; Fig. 3A). The point mutations were introduced into cDNA sequences for expression in yeast and bacteria. In Y2H screens, the BIN2-TRS120 interaction, but not TRAPPII complex interactions, was almost abolished when all three sites were phosphomimetic (TRS120-T2 SαβγD; Fig. 2F). As kinases typically have kiss and run interactions with their unphosphorylated substrates, and as BIN2 interacts more strongly with an unphosphorylated than a phosphorylated substrate (Pusch *et al*, 2012; Tang *et al*, 2011), this is consistent with AtTRS120/TRAPPC9 being targeted by the BIN2 kinase.

**Figure 3.**
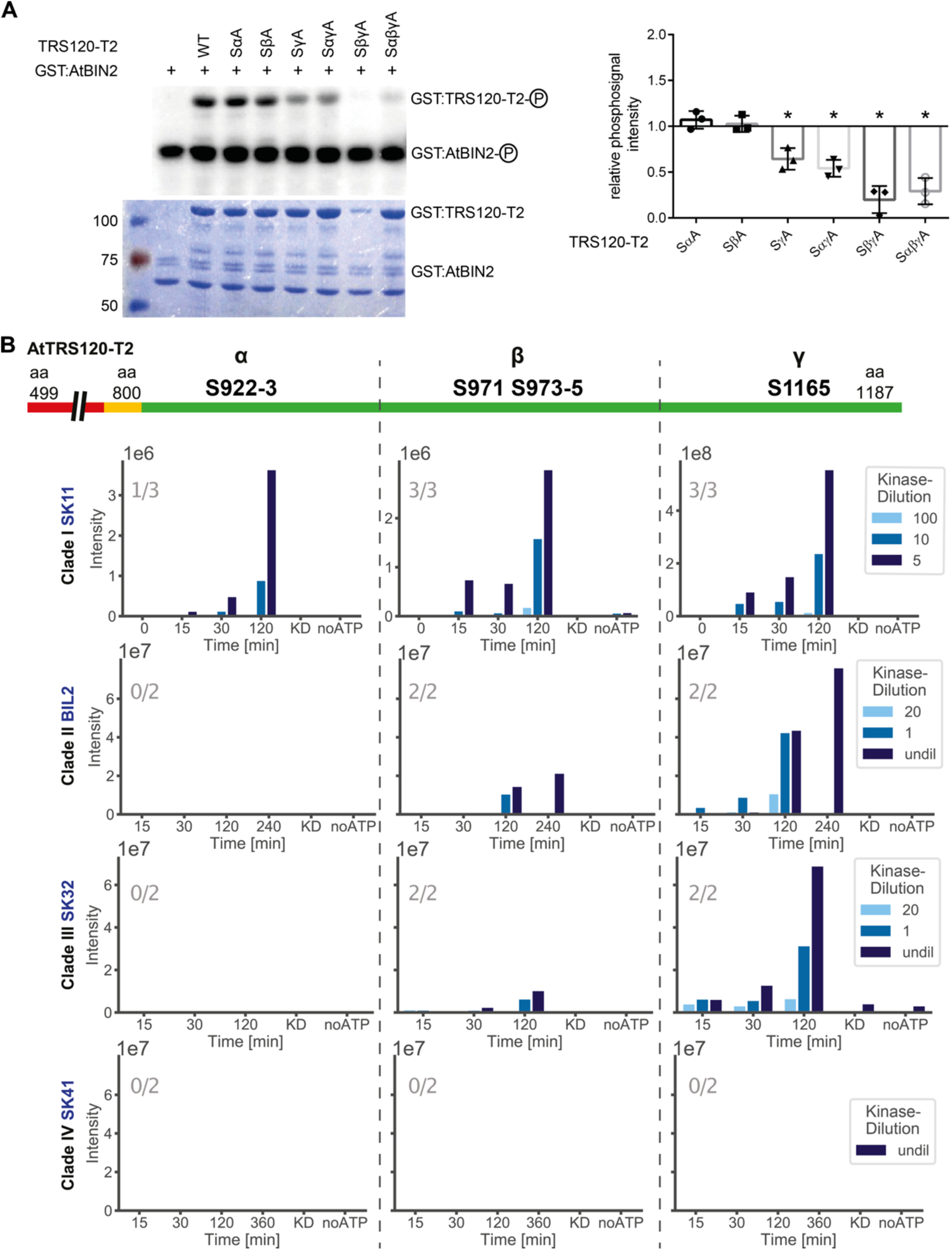
*In vitro* AtSK kinase assays with TRS120 as bait. **A.** *In vitro* kinase assays using GST:AtBIN2 (69 kDa) and GST:TRS120-T2 (100 kDa). The change of phosphosignal is shown in a representative autoradiograph (upper panel) and the loaded protein amount in the corresponding CBB (Coomassie stain, lower panel). Non-phosphorylatable S to A TRS120-T2 variants were used as negative controls. The means ± SD of phosphosignals were normalized to the protein amount and related to non-mutated TRS120-T2 wild-type control. Note that BIN2 phosphorylated AtTRS120-T2 *in vitro,* with a preference for wild-type (WT) sequences over non-phosphorylatable AtTRS120-SγA, AtTRS120-SαγA, AtTRS120-SβγA and AtTRS120-SαβγA substrates. n = 3 independent experiments; *: P<0.05 for significant differences to TRS120-T2 WT (set at 1.0 right panel) determined by using a one sample two-tailed t-test. **B.** Kinase assays were performed *in vitro* with mass-spectrometry readout. One member of each shaggy-like kinase clade (AtSKs; see Fig. S3A) was used with the TRS120-T2 truncation as substrate. Dilution series of the kinase are depicted in different shades of blue. AtTRS120-T2 has highly (red) and moderately (orange) conserved sequences, as well as plant-specific sequences (green). Three GSK3 sites (referred to as α, β, γ; see Fig. 2C) can be found in the plant-specific T2 domain. AtSKs in clades I-III differentially phosphorylated the substrate at three GSK3 consensus sites (with a preference for the γ site) in a time-dependent and concentration-dependent manner. A clade IV AtSK did not phosphorylate at all. Samples incubated for 120 min in a kinase buffer without ATP, or samples in which the kinase was heat inactivated (KD), served as negative controls. The numbers in grey in each plot denote the number of times the phosphorylation event was seen in the given number of independent replicates. Note the higher intensity of the TRS120-γ peptide, especially for Clade I/SK11 (1e8 for TRS120-γ versus 1e6 for TRS120-α and TRS120-β on the Y axis). Related to Fig. S2-S6.

The IP-MS and Y2H interactions were validated with *in vitro* kinase assays, performed with a phosphorus radioisotope (Fig. 3A; Fig. S5A). This showed that BIN2 and AtSK11 phosphorylated AtTRS120-T2 *in vitro,* with a preference for wild-type sequences over non-phosphorylatable substrates such as AtTRS120-SαβγA (Fig. 3A; Fig. S5A). Further, the phosphorylation was confirmed with mass-spectrometry using non-radioactive assays. The mass-spectrometry results showed that the AtTRS120 α, β, and γ sites phosphorylated *in vivo* (Fig. S2; IP-MS on seedlings using TRS120:GFP as bait) were phosphorylated by AtSKs *in vitro* (Fig. 3B; Fig. S5B). Kinase assays performed with varying kinase concentrations (but constant substrate concentrations) showed that the phosphorylation events were time and/or concentration dependent (Fig. 3B; Fig. S5B). AtTRS120 was a substrate of AtSKs in clades I-III; we did not detect phosphorylation of AtTRS120 with a clade IV AtSK *in vitro* (Fig. 3B). All AtSKs that targeted AtTRS120 had a marked and consistent preference for the TRS120-γ (S1165) site (Fig. 3B; Fig. S5B). *In vivo*, IP-MS performed on seedlings treated with the AtSK inhibitor bikinin showed a reduced extent of phosphorylation of the TRS120-γ peptide (Fig. S6). Conversely, seedlings treated with the BR biosynthesis inhibitor PPZ, which should relieve BR-mediated BIN2 inhibition, showed an increased extent of phosphorylation of the TRS120-γ peptide (Fig. S6). These *in vivo* observations are consistent with the *in vitro* kinase assays (Fig. 3; Fig. S5). In summary, several lines of *in vitro* (Y2H, kinase assays) and *in vivo* (IP-MS, pharmacological inhibition) evidence support the conclusion that the TRAPPII subunit AtTRS120 is a substrate of shaggy-like kinases.

### BIN2 and TRAPPII are required for differential growth decisions under additive stress

As TRAPPII is a BIN2 substrate, a question is whether *bin2* and *trappii* mutants have related phenotypes. We were not able to detect cytokinesis or protein sorting defects, characteristic of *trappii,* in semi-dominant *bin2-1* alleles (Fig. S7). We have recently shown that BIN2 is required for hypocotyl versus root trade-offs in the germinating seedling under additive stress conditions involving the simultaneous withdrawal of both light and water (Kalbfuß *et al*, 2022). Water stress in the dark is a “conflict of interest” scenario in which hypocotyl and root growth have competing interests (Kalbfuß *et al*, 2022). Kalbfuß *et al* (2022) defined decision mutants as ones that had either insignificant hypocotyl/root-ratio responses (P > 0.05), or consistently wrong growth responses as compared to the wild type (P < 0.05 but for an opposite growth phenotype, as depicted by red asterisks in Fig. 4B-D; Kalbfuß *et al*, 2022). Under additive stress, *trappii* null mutants failed to adjust their hypocotyl length along the same line as the wild type (red asterisks in Fig. 4C-D; Fig. 4F; see Fig. S8D-E for a direct comparison to the wild type and for the distribution of datapoints). In particular, *trappii trs120-4* mutants had non-significant hypocotyl/root ratio responses to water withdrawal in the dark (Fig. 4D; Fig. S8E). In addition to comparing organ lengths, each mutant line was normalized to its corresponding wild-type ecotype on the same (PEG) plate, which helped us to take the variability between PEG plates and experiments into account and enabled us to pool biological replicates (see methods for further detail). To this end, the response to water stress in the dark was represented as a normalized response quotient (RQ). The RQ is an indication of how well each mutant responds to a given combination of stress cues and indicates how much each line deviates from the wild type. A value of 1.0 means that the mutant line behaves exactly like its corresponding wild type. This rendition shows that *bin2* higher order and *trappii* null alleles considerably deviated from the wild type, with severely attenuated responses (Fig. 4G; Fig. S9). We reason that decision mutants unable to integrate environmental cues might have highly variable hypocotyl versus root lengths. This high variance would, in turn, translate into an insignificant (i.e. high) *P*-value, indicative of a low signal-to-noise ratio. We, therefore, plotted the median *P*-values against the normalized response quotients (referred to as volcano plots; mean RQ_ratio_ in Fig. 4H). Wild-type ecotypes had significant *P*-values < 10^-10^ (grey shading on the red line in Fig. 4H, green arrow). Mutants with insignificant *P*-values and response quotients considerably smaller than 1.0 would be considered “confused” decision mutants and these would map in the lower left quadrant of the RQ_ratio_ volcano plot (see peach shading in Fig. 4H). *bin2* higher order null alleles and *trs120-4* mutants clustered together in the lower left region of the volcano plot (see peach shading in Fig. 4H); both qualify as decision mutants (Fig. 4B, 4D, 4G-H; Kalbfuß *et al*, 2022). This was in contrast to the near-wild-type phenotype of higher order null mutants impaired in the perception of light or water stress (*phyAphyBcry1cry2* and *pyr1pyl1pyl2pyl4* in Fig. 4G-H). We conclude that *bin2* higher order and *trappii* null alleles are decision mutants.

**Figure 4.**
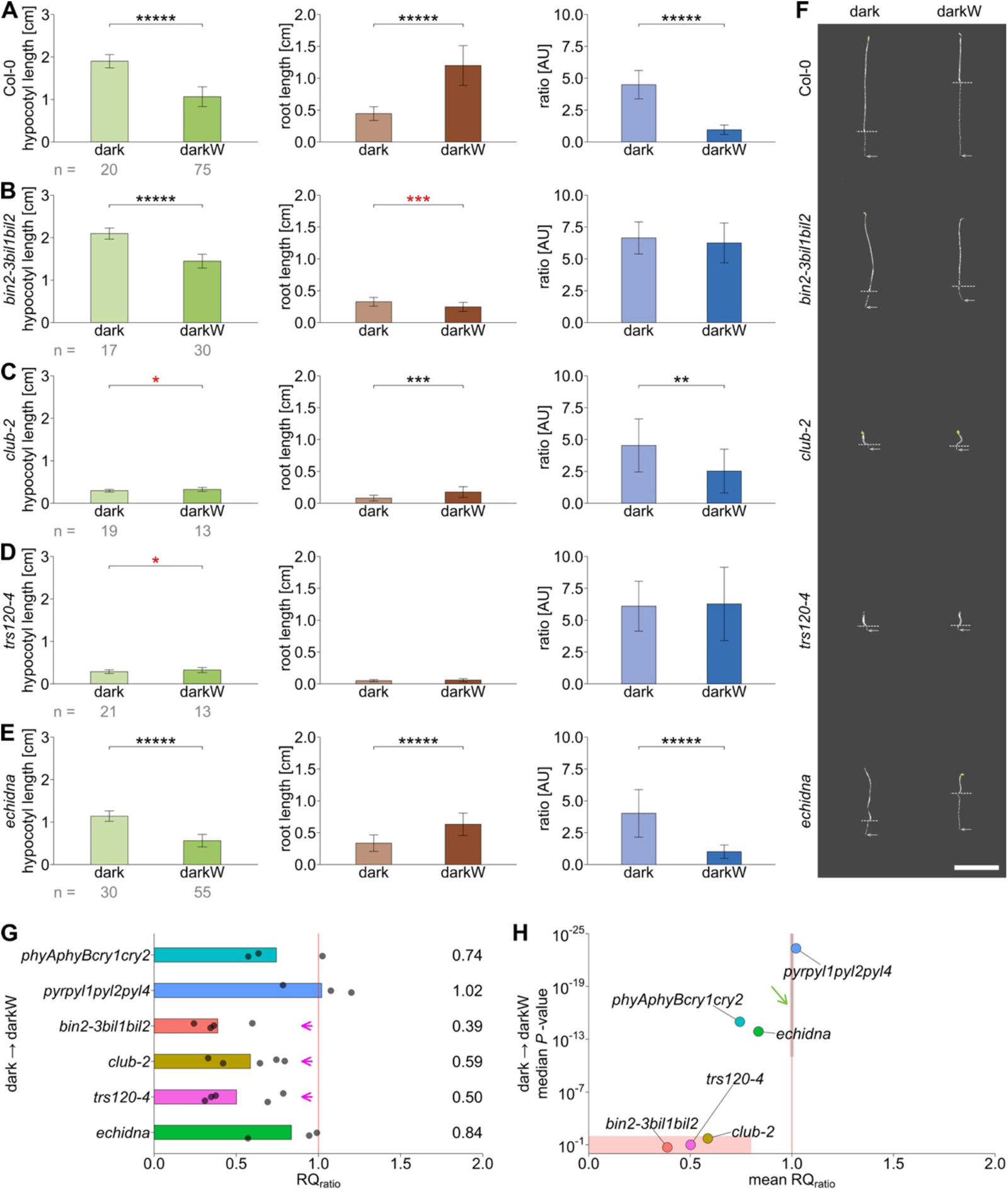
Role of the TRAPPII complex versus ECHIDNA in hypocotyl/root trade-offs. Seedlings were germinated on ½ MS in the dark (dark) or in dark with -0.4 MPa water stress (darkW). **A.** Col-0 (wild type). **B.** BR signaling mutant *bin2-3bil1bil2* triple knockout (from Kalbfuß *et al*, 2022). **C.** *club-2*, a null *trappii* mutant. **D.** *trs120-4*, a null *trappii* allele. **E.** *echidna* null allele, impaired in TGN structure and function. At least 3 experiments were performed for each line, and a representative one is shown here on the basis of RQ and *P*-values. **F.** Representative seedling images of Col-0 (wild type) and mutants shown in A-E. *bin2-3bil1bil2* and *trs120-4* mutants failed to correctly adjust their hypocotyl and root lengths from dark to darkW conditions. Dotted lines mark the hypocotyl-root junction, whereas arrows point to the end of the root. Scale bar is 1 cm. **G.** Normalized response quotient RQ_ratio_. Each replicate is represented by a dot. A value of 1 (vertical red line) corresponds to an identical adaptation to darkW conditions as the wild type. Note that the triple *bin2-3bil1bil2* knock out, *trs120-4* and *club-2* had attenuated responses (magenta arrows). **H.** Volcano plot with the mean RQ_ratio_ depicted on the X axis and the median *P*-value of the response on the Y axis (negative log scale; a median of all replicates was used). The area shaded in grey on the red line (green arrow) is where wild-type ecotypes would theoretically map onto the plot. Mutants in the lower left quadrant (peach shading) were considered to have a “confused decision phenotype” (see text). *trs120-4* mutants mapped to the lower left quadrant and qualified as decision mutants on two counts: (i) a consistently opposite hypocotyl response (red asterisk in D), (ii) failure to adjust the hypocotyl/root ratio to darkW (the ratio for darkW is the same as for dark in panel D), translating into a non-significant *P*-value for the ratio response (H). The number (n) of seedlings measured per condition is in grey below the mean ± SD bar graphs. *P*-values were computed with a two-tailed student’s t-test and are represented as follows: *: P<0.05; **: P<0.01; ***: P<0.001; ****: P<0.0001; *****: P<0.00001. Mutant alleles and the corresponding ecotypes are described in Table S1. Related to Fig. S8-S9.

A question that arises is whether *trappii* mutants are impaired in differential growth decisions as a secondary consequence of primary defects in morphogenesis or cytokinesis (Jaber *et al*, 2010; Rybak *et al*, 2014; Thellmann *et al*, 2010). The etiolation response was severely attenuated in *trappii* mutants, but nonetheless highly significant (P < 0.00001; Fig. S10C-D, S10F-I). Similarly, an attenuated but clear etiolation response has been shown for other cytokinesis-defective mutants including *keule* and *knolle* (Assaad *et al*, 2001). This shows that, despite a severe impairment in cell division and morphogenesis, cytokinesis-defective mutants are nonetheless capable of differential growth. In addition to their cytokinesis defect, *trappii* mutants are impaired in TGN function (Ravikumar *et al*, 2018). We, therefore, compared *trappii* mutants to *echidna* mutants, severely impaired in TGN structure and function (Boutté *et al*, 2013; Gendre *et al*, 2013; McFarlane *et al*, 2013). Both *echidna* and *trappii* mutants exhibited a severe impairment in root elongation in the light (Fig. S10C-F; Fig. S10H). In contrast to *trappii*, however, *echidna* mutants had highly significant responses to additive stress that resembled the wild type in all respects (Fig. 4E-H; Fig. S8F). Thus, *echidna* mutants do not qualify as decision mutants. In conclusion, a comparison to other cytokinesis-defective or TGN mutants suggests that neither cytokinesis defects nor TGN malfunction suffice to explain the *trappii trs120-4* decision phenotype.

Cellular growth parameters in *trappii* were assessed under single versus additive stress, in both the hypocotyl and root tip. The width, height and surface area of *trappii* hypocotyl cells grown in the light did not show any deviation from the wild type (Fig. 5A-B light; Fig. S11A-D light). In wild-type hypocotyls, both organ length and cell length decreased in response to water stress in the dark (Fig. 4A; Fig. 5A; Fig. S11C). In contrast, in *trappii* mutants, organ and cell length significantly increased (red asterisks or compact letter displays in Fig. 4C-D, Fig. 5B and Fig. S11A, C-D highlight a phenotype consistently opposite to the wild type). This was more pronounced in *trs120-4* (1.71-fold increase; Fig. 5B) than in *club-2* (1.46-fold increase; Fig. S11A) whereas the wild-type Col-0 showed a 0.55-fold decrease in cell length (Fig. 5A). In root tips, we monitored meristem properties and cell length along single cortical cell files as a function of distance from the quiescent center. In the wild type, meristem size was large in the light, intermediate in the dark and shortest under water stress in the dark (darkW; Fig. 5C). In contrast, meristem size in *trs120-4* remained constant under the three environmental conditions tested (Fig. 5D). We have recently shown that root growth in response to water stress in the dark is due to a combination of cell division and rapid exit from the meristem (Kalbfuß *et al*, 2022). An early exit from the meristem can be visualized as cell elongation in cells close to the quiescent center. This was observed under dark and darkW conditions in the wild type (green arrows in Fig. 5C) but not in the *trappii* mutant *trs120-4* (magenta arrows in Fig. 5D). While the curves differed under the different environmental conditions in the wild type (Fig. 5C), these were fairly similar regardless of the environmental cue in *trs120-4* (magenta arrows in Fig. 5D; note that the grey shading, which designates the 95% confidence interval, overlaps). We conclude that, at a cellular level, *trappii trs120-4* mutants are unable to differentially regulate their growth parameters in response to additive stress (Fig. 5; Fig. S11). This would suffice to explain the growth defects we observed at the organ level (Fig. 4). The *trappii* cellular phenotype in the decision screen is reminiscent of that reported for *bin2* (Kalbfuß *et al*, 2022). In summary, *bin2* and *trappii* alleles have related phenotypes with respect to an inability to differentially regulate cell growth in both the hypocotyl and root tip in response to additive stress (Figs. 4-5; cf. Kalbfuß *et al*, 2022).

**Figure 5.**
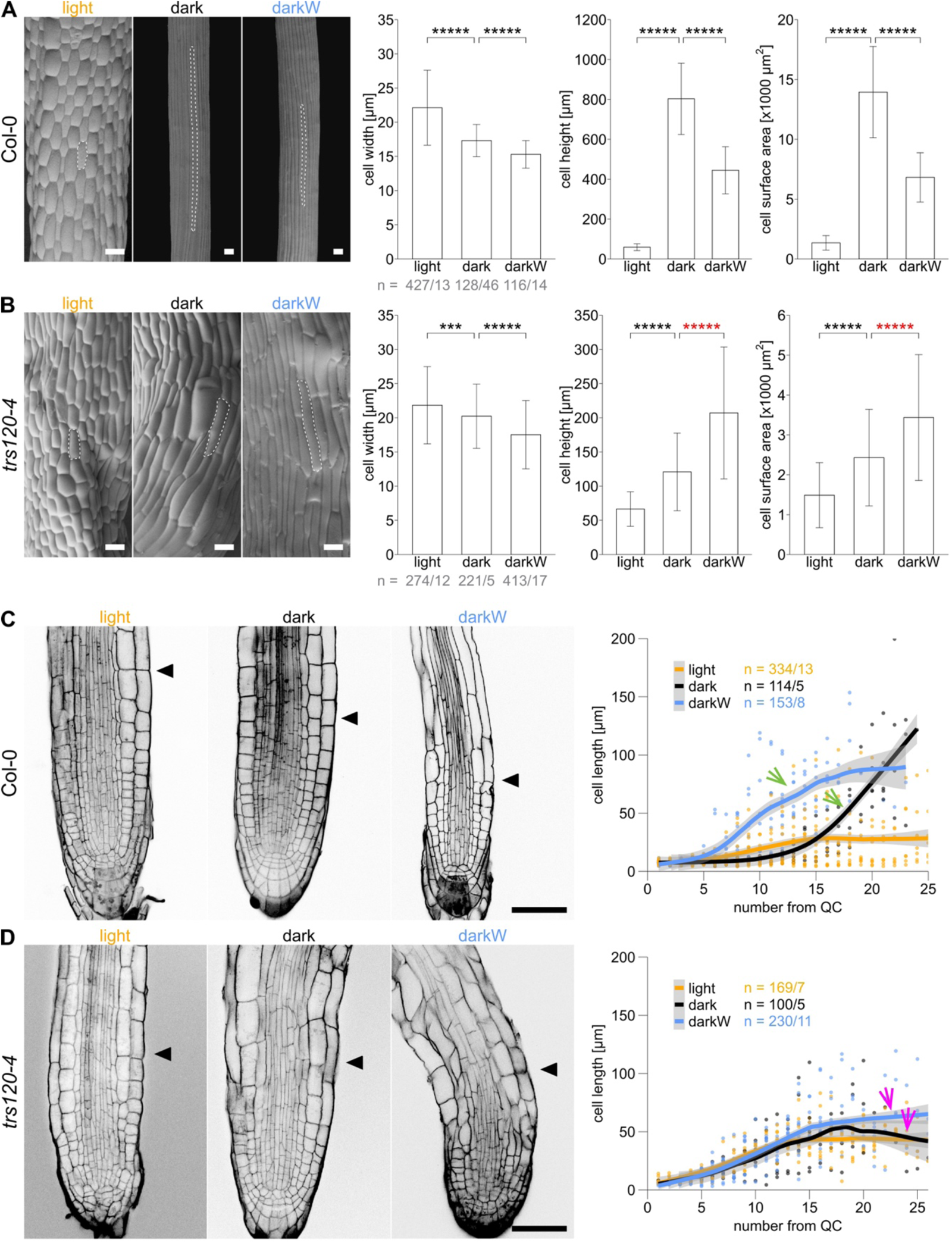
Cellular growth parameters of *trappii* mutants under single versus additive stress conditions. Seedlings were grown in the light (orange), dark (black) or dark with -0.4 MPa water stress (darkW; blue). **A, B.** Scanning electron micrographs (SEM) of hypocotyls of A) Col-0 wild type and B) *trs120-4,* null *trappii* allele. The cell surface area was calculated as the product of cell width and length. *trs120-4* mutants showed the opposite hypocotyl cell height and cell surface adaptations to dark-to-darkW conditions (red asterisks in B) than the wild-type control. Representative SEM images of seedlings grown under the specified conditions are shown (scale bar = 40 µm). Outlines of representative cells are highlighted by white dashed lines. Shown are means ± SD. *P*-values were computed with a two-tailed student’s t-test and are represented as follows: ***: P<0.001; ****: P<0.0001; *****: P<0.00001. **C, D.** Confocal micrographs of mPS-PI stained C) Col-0 wild type and D) *trs120-4* null mutant root tips under different environmental conditions; black arrowheads mark the junction between the meristematic and elongation zones. 10 days after incubation, the cell lengths were measured in single cortex cell files, starting at the cortex/endodermis initials. Cell lengths of consecutive cells were mapped as a function of cell number from the quiescent centre (QC). The fitted lines were generated with Local Polynomial Regression Fitting with the ‘loess’ method in R; grey shading designates the 95 percent confidence interval. Col-0 seedlings grown in the dark with and without water stress show steep slopes (green arrows). In the *trappii* mutant the curves exhibit a minimal to no difference between the different screen conditions (magenta arrows). Scale bar is 50 µm. The sample size (n) is given in each panel as the number of cells/ number of seedlings that were analyzed (A-D). Related to Fig. S11.

### AtTRS120 phosphovariants are functional

To assess the *in vivo* impact of the TRS120 phosphorylation status on intracellular localization, targeted point mutations at the TRS120 AtSK sites (Fig. 2C) were introduced into genomic sequences capable of rescuing the *trs120-4* mutant phenotype (Rybak *et al*, 2014). The TRS120 constructs were fused to a C-terminal GFP tag and we refer to the ensuing site-directed mutants as TRS120 phosphovariants. The constructs were expressed and capable of rescuing the null *trs120-4* allele in the T1 and T2 generations (Fig. S12A-B); upon further propagation, however, signs of silencing were evident in seedlings (Fig. S12C-D). The wild-type TRAPPII complex resides in the cytosol, at the TGN and at the cell plate (Fig. S13; Naramoto *et al*, 2014; Qi *et al*, 2011; Ravikumar *et al*, 2018; Rybak *et al*, 2014). Phosphovariants had a similar appearance, but the phosphomimetic TRS120-SαβγD variant tended to be mostly membrane associated, with almost no detectable cytosolic signal (Fig. S13).

### TRS120-SαβγA and TRS120-SαβγD have opposite effects on seed germination under osmotic stress

We first analyzed the impact of osmotic stress on the germination frequencies of TRS120 phosphovariants. Different TRS120-SαβγA and TRS120-SαβγD primary transformants were plated on media containing 0, 200, or 400 mM mannitol (Fig. 6A-C). As compared with the control (Col-0), the TRS120-SαβγA phosphovariant had higher seed germination rates on mannitol: on day 4 we observed 70% germination in TRS120-SαβγA versus 35% germination for the control on 400 mM mannitol as an osmotic agent (Fig. 6C). In this respect, TRS120-SαβγA was similar to the ABA deficient mutant *aba2-1*, known to be osmotolerant with respect to seed germination (Fig. 6C). In contrast, the TRS120-SαβγD phosphovariant exhibited delayed and reduced germination, with lower maximal germination rates, even in the absence of mannitol (Fig. 6A-C). Indeed, on 400 mM mannitol TRS120-SαβγD mutants reached a maximal germination rate of 60% compared to 90% for the wild type. With respect to its delay in germination, TRS120-SαβγD was similar to the ABA coreceptor higher order mutant *hab1-1 abi1-2 pp2ca-1*, known to be osmosensitive at germination (Fig. 6C). In summary, the non-phosphorylatable TRS120-SαβγA mutations enhanced whereas the phosphomimetic TRS120-SαβγD mutations reduced germination on mannitol.

**Figure 6.**
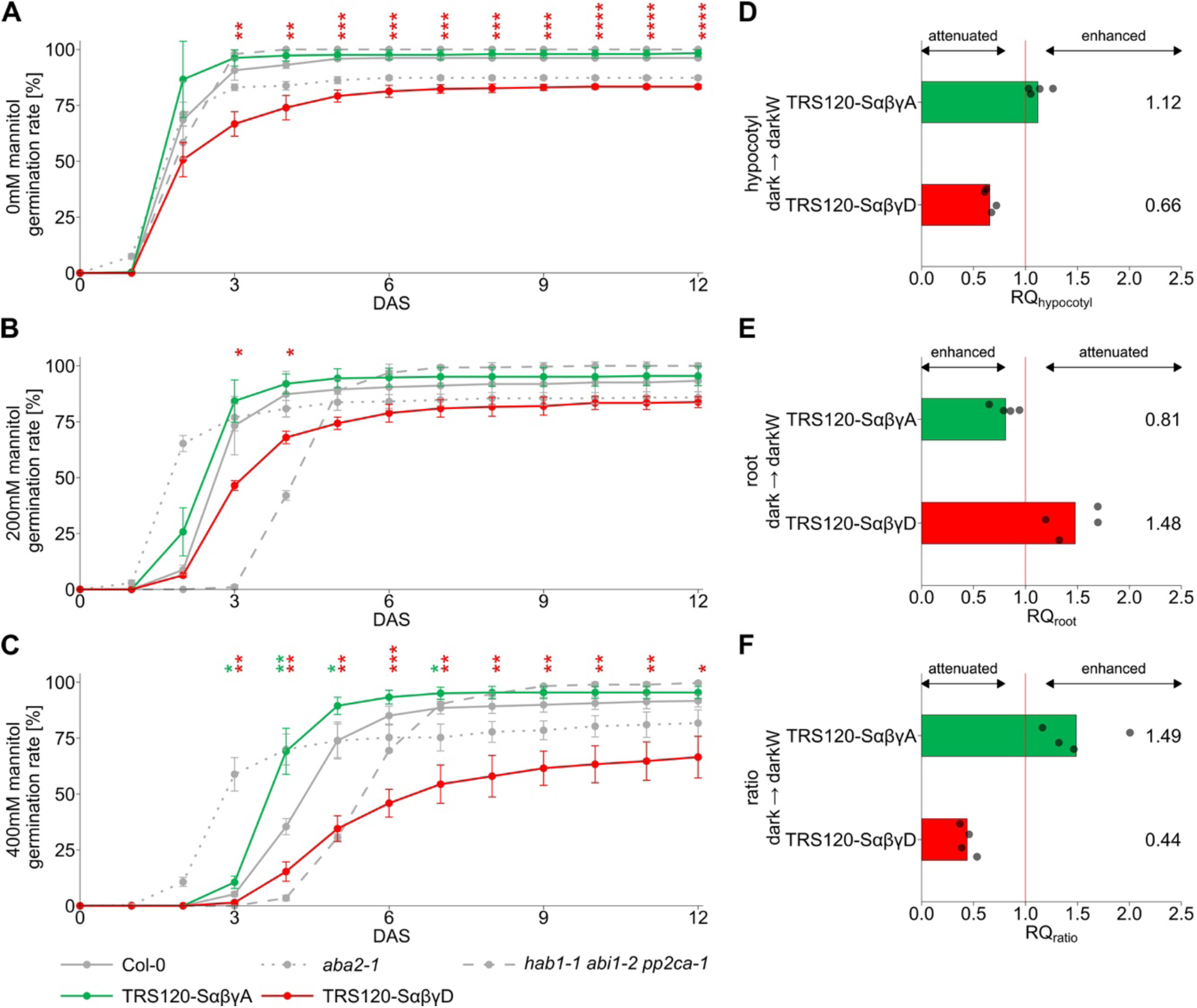
Responses of TRS120 phosphovariants to osmotic or additive stress. Phosphovariants of P_TRS120_::TRS120:GFP (Rybak *et al*, 2014) were in *trs120-4* homozygous backgrounds with the exception of the TRS120-SαβγD variant in panels A-C, which was in a hemizygous *trs120-4*/+ background. **A-C.** Germination of TRS120 phosphovariants under osmotic stress. Effect of A) 0 mM, B) 200 mM and C) 400 mM mannitol on seed germination of the TRS120-SαβγA, the TRS120-SαβγD mutants and the following controls: Col-0 wild type; the ABA deficient mutant *aba2-1* and the ABA PP2C coreceptor triple knock-out line *hab1-1 abi1-2 pp2ca-1*. The non-phosphorylatable TRS120-SαβγA mutation increased seed germination on higher mannitol concentrations, whereas the phosphomimetic TRS120-SαβγD decreased seed germination. Shown are the means ± SD of three technical replicates. Similar results were obtained in three independent biological replicates. Stars indicate statistical significance compared to the wild-type Col-0 calculated with a two-tailed student’s t-test (*: P<0.05; **: P<0.01; ***: P<0.001; ****: P<0.0001). DAS: days after stratification. **D-F**. Response quotients (RQ) of D) the hypocotyl, E) the root and F) the hypocotyl/root ratio of the non-phosphorylatable TRS120-SαβγA and the phosphomimetic TRS120-SαβγD mutants for dark-to-dark with water stress (-0.4 MPa, darkW) adaptation. The divergent hypocotyl and root responses of TRS120-SαβγA and TRS120-SαβγD mutants resulted in a strong hypocotyl/root ratio phenotype under darkW conditions compared to the TRS120-WT control. TRS120-SαβγA showed an enhanced ratio adaption, whereas TRS120-SαβγD mutants had an attenuated ratio response. RQ response quotients are normalized to the rescue mutant TRS120-WT (P_TRS120_::TRS120:GFP in *trs120-4/trs120-4*). A value of 1 (vertical red line) corresponds to an identical adaptation to darkW conditions as the rescue mutant. Note that due to the opposite adaptations of the hypocotyl and root under the additive stress conditions (decrease in hypocotyl and increase in root length from dark-to-darkW), the respective thresholds for attenuated and enhanced responses are opposite. Dots represent biological replicates, of which there were a total of four primary transformants per line. Mean RQ values are given on the right. Related to Fig. S14-S15.

### TRS120 phosphorylation status affects hypocotyl versus root trade-offs under additive stress conditions

In light of the decision phenotypes of *bin2* and *trappii* mutants and of the observation that AtSKs such as BIN2 target TRAPPII, the question arises as to whether shaggy-like kinases regulate differential growth decisions via TRS120 phosphorylation. To test this hypothesis, AtTRS120 phosphovariants homozygous for the null *trs120-4* allele and hemizygous for the phosphovariant construct were studied under different environmental or stress conditions. The etiolation response was not impacted by the phosphorylation status of AtTRS120 (Fig. S14). In contrast, TRS120 phosphomutants had phenotypes under additive stress conditions (Fig. 6D-F; Fig. S15A-C). TRS120-SαβγA non-phosphorylatable variants exhibited an enhanced root response under darkW conditions (Fig. 6E) and a strongly enhanced ratio response to water stress in the dark (mean RQ_ratio_ = 1.49; Fig. 6F). Conversely, the phosphomimetic TRS120-SαβγD mutants exhibited severely attenuated responses to water stress in the dark in all respects (Fig. 6D-F; mean RQ_hypocotyl_= 0.66; mean RQ_root_ = 1.48; mean RQ_ratio_ = 0.44). Thus, the non-phosphorylatable TRS120-SαβγA and the phosphomimetic TRS120-SαβγD mutations had opposite effects on hypocotyl versus root lengths under water stress in the dark, with TRS120-SαβγA exhibiting an enhanced and TRS120-SαβγD an attenuated ratio response (Fig. 6F). Under all environmental conditions, the total seedling length of the phosphovariants did not significantly differ from that of the wild-type AtTRS120 construct (Fig. S15D), which highlights the absence of a growth defect. We conclude that the TRS120 phosphorylation status impacts the seedling’s ability to differentially regulate its hypocotyl and root lengths in response to additive stress conditions. Taken together, the data show opposite impacts of non-phosphorylatable versus phosphomimetic mutations at AtTRS120 AtSK sites in terms of seed germination under osmotic stress and responses to additive stress. This suggests that TRAPPII phosphorylation by shaggy-like kinases mediates adaptive responses to osmotic stress and to light and water availability.

### *bin2* higher order and *trappii* conditional mutants exhibit a synergistic genetic interaction with respect to root gravitropism

To further address the physiological significance of the BIN2-TRAPPII interaction *in vivo*, we deployed higher order mutant analysis. This approach, however, was challenged by (i) the functional redundancy between BIN2 and its homologues (Vert & Chory, 2006; Yan *et al*, 2009), (ii) the semi-dominant nature of *bin2-1*, (iii) the seedling lethality of *trappii* null alleles, as well as (iv) the pleiotropic phenotypes of both *bin2-1* and *trappii* dwarfs (Li *et al*, 2001; Thellmann *et al*, 2010; Ravikumar *et al*, 2018; Garcia *et al*, 2020). To address these challenges, we engineered a conditional *trs120* knock-down allele, named *trs120i*. This features an artificial microRNA that targets 5’ AtTRS120/TRAPPC9 sequences, expressed under an estradiol-inducible promoter (Fig. 7A; Curtis & Grossniklaus, 2003). The *trs120i* construct was introduced into the wild type (Col-0) and into the *bin2-3bil1bil2* triple knock-out mutant. Upon induction, *trs120i* exhibited a mild and *bin2-3bil1bil2* a more pronounced root agravitropism (Fig. 7B-C). The *bin2-3bil1bil2 trs120i* higher order mutant had a more than additively enhanced agravitropic response upon induction as compared to *trs120i* or *bin2-3bil1bil2* alone (Fig. 7B-C, 7E). This was evidenced as primary roots growing in all directions, often against gravity at 180° in *bin2-3bil1bil2 trs120i* (Fig. 7B-C). We observed synthetic enhancement specifically upon estradiol induction and not in the mock control (Fig. 7D, compared to 7E). This is indicative of a synergistic genetic interaction. Furthermore, it suggests that adaptive growth decisions such as gravitropism are mediated by the BIN2/AtSK-TRAPPII interaction.

**Figure 7.**
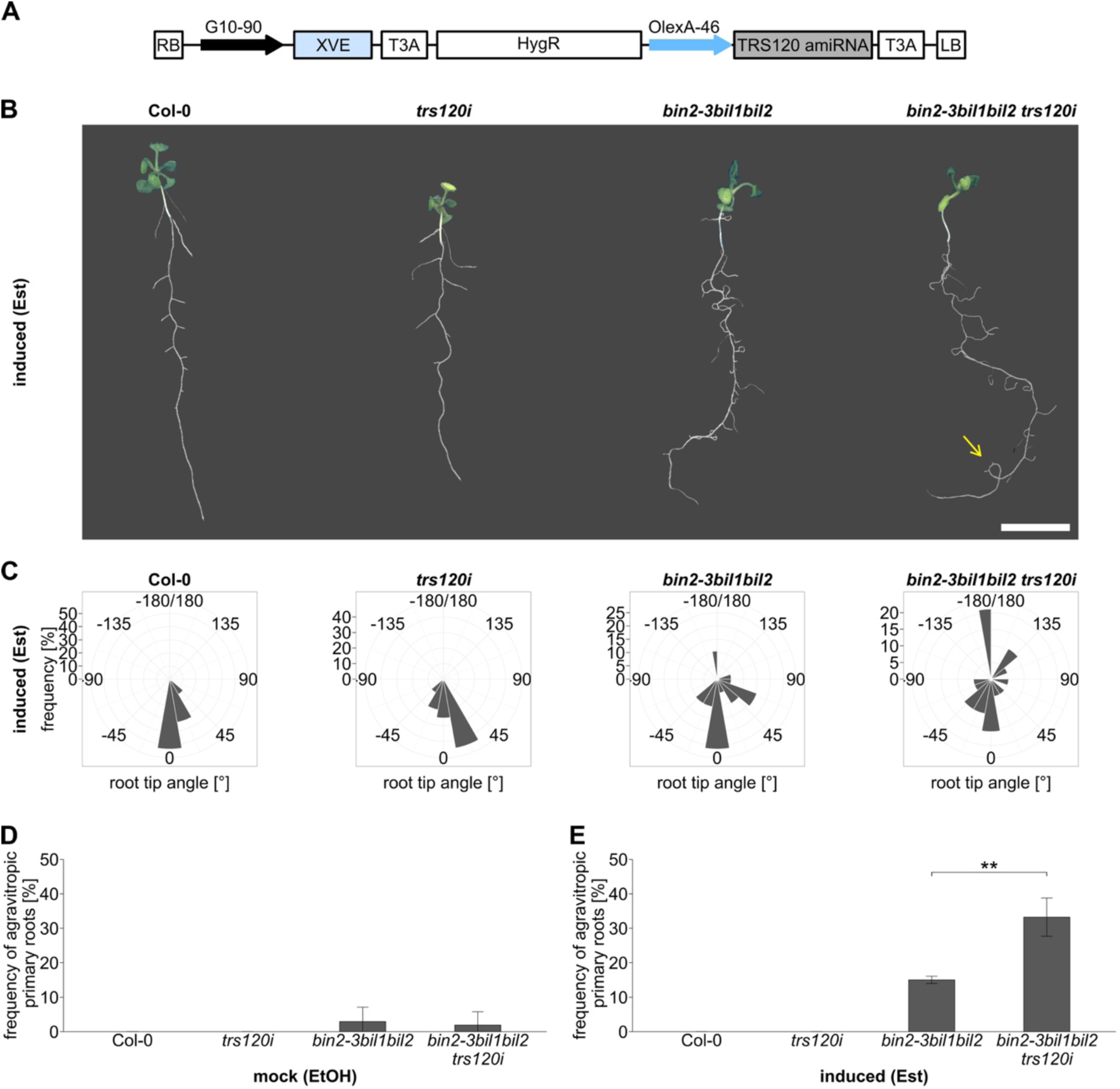
Genetic interaction between *bin2* higher order and *trappii* conditional mutants. Seedlings were transferred to estradiol (Est; induced) or EtOH (mock) containing media and grown vertically for 9 days. **A.** A conditional *trs120* knock-down allele, *trs120i*. This features an artificial microRNA targeting the 5’ end of the AtTRS120/TRAPPC9 coding sequence, expressed under an estradiol-inducible promoter. LB: left border; RB: right border; G10-90: synthetic constitutive promoter; XVE: encoding a chimeric transcription factor composed of the DNA-binding domain of the bacterial repressor LexA (X), the acidic transactivating domain of VP16 (V) and the regulatory region of the human estrogen receptor (E); T3A: poly(A) addition sequence; HygR: hygromycin resistance marker; OlexA-46: estradiol inducible promoter system with multiple copies of the lexA binding domain. **B.** Representative seedling images of Col-0 (wild type), *trs120i* mutant, *bin2-3bil1bil2* and *bin2-3bil1bil2 trs120i* higher order mutants grown on estradiol. Upon induction, *trs120i* exhibited a mild phenotype, with a wavy primary root, and *bin2-3bil1bil2* a more pronounced root agravitropism. In contrast, *bin2-3bil1bil2 trs120i* mutants had an enhanced root agravitropism, growing against gravity or in loops (yellow arrow points to a 360° loop in the primary root). Scale bar is 1 cm. **C.** Visualisation of the primary root growth orientation. Primary root tip angles (excluding lateral roots) of Col-0, *trs120i*, *bin2-3bil1bil2* and *bin2-3bil1bil2 trs120i* seedlings grown on Est were measured and a frequency count with a bin width of 20° was performed. Frequency plots were generated by using a polar coordinate system. A root tip angle of 0° corresponds to vertical root tip growth in line with the gravitropic vector. Primary roots exhibiting looping (see yellow arrow in B) were scored as -180° due to their growth against gravity. A representative plot for each genotype is shown. **D, E.** Quantification of the frequency of agravitropic primary roots upon growth on D) mock (EtOH) or E) induced (Est) conditions. Frequency of primary root tips growing against gravity (angles < – 90° and > 90° in panel C) was counted per experiment. The *bin2-3bil1bil2 trs120i* higher order mutant had a more than additively enhanced agravitropic response upon induction as compared to *trs120i* or *bin2-3bil1bil2* alone (E). Note the synthetic enhancement specifically upon estradiol induction and not in the mock control (Fig. 7D, compared to 7E). This is indicative of a synergistic genetic interaction. Mean ± SD bar graphs; **: P≤0.01 two-tailed student’s t-test. n = 2 independent experiments for Col-0 and *bin2-3bil1bil2*, n = 4 for *bin2-3bil1bil2 trs120i* and n = 6 for *trs120i* with at least 11 seedlings per experiment.

## DISCUSSION

In this study, we explore the role of the TGN in stress responses in Arabidopsis. Our point of entry is the TRAPPII complex, which has been shown to be required for TGN structure and function (Qi *et al*, 2011; Ravikumar *et al*, 2018). We performed proteomic and yeast two-hybrid screens and present several lines of *in vitro* (Y2H, kinase assays) and *in vivo* (IP-MS, pharmacological inhibition) evidence that the TRAPPII subunit AtTRS120/TRAPPC9 is the target of AtSK kinases, including BIN2. We document differential phosphorylation of three distinct AtSK/GSK3 sites by three of the four AtSK clades. The phosphorylation status of AtTRS120 impacted seed germination under osmotic stress as well as adaptive responses to additive stress *in planta*. We show that *bin2* and *trappii* alleles have related phenotypes with respect to an impaired adaptation to additive stress conditions, in this instance achieved by the simultaneous withdrawal of light and water. Furthermore, *bin2* higher order and *trappii* conditional mutants exhibited a synergistic genetic interaction with respect to root gravitropism.

Like other tropisms, root gravitropism requires organ bending as a result, at least in part, of the differential sorting of PIN transporters (Friml *et al*, 2002; Ding *et al*, 2011; Konstantinova *et al*, 2021). PIN2 polarity has been shown to be abolished in *trappii* mutants (Qi *et al*, 2011; Rybak *et al*, 2014) and to require TGN function (Naramoto *et al*, 2014; Qi *et al*, 2011; Rybak *et al*, 2014; Ravikumar *et al*, 2018). A concern when assessing the role of the TGN in adaptive or stress responses is that TGN-related mutants typically have pleiotropic phenotypes as a consequence of their impairment in fundamental processes such as secretion, endocytosis or sorting (Rosquete & Drakakaki, 2018; Ravikumar *et al*, 2018). To distinguish between primary defects in growth on the one hand and adaptive responses on the other, we monitored trade-offs between hypocotyl and root growth under additive stress conditions (Kalbfuß *et al*, 2022). We focus on *echidna* and *trappii* mutants as these have been shown to impact both the structure and the function of the TGN (Boutté *et al*, 2013; Gendre *et al*, 2011, 2013; McFarlane *et al*, 2013; Qi *et al*, 2011; Ravikumar *et al*, 2018). These TGN mutants had hypocotyl and root growth defects, which are characteristic of trafficking mutants (Fig. S10). In addition to their growth defects, *trappii trs120-4* mutants had a significant hypocotyl response but in the wrong orientation as compared to the wild type (denoted by red asterisks in Fig. 4, Fig. 5 and Fig. S11) as well as an insignificant hypocotyl/root ratio response to light and water deprivation (Fig. 4). The *trappii* phenotypes described in this study are difficult to explain as mere growth defects. Rather, the inability to mediate growth trade-offs in response to additive stress provides a compelling argument for TRAPPII as having a role in adaptive growth decisions.

Conceptually, the optimization of hypocotyl-to-root ratios can be deconstructed into four distinct stages: (i) perception, (ii) signal integration, (iii) decision making and (iv) the implementation of resulting actions (Kalbfuß *et al*, 2022). To address the first stage, perception, we looked at, for example, quadruple *phyAphyBcry1cry2* photoreceptor mutants; these failed to adjust their organ lengths to light versus dark conditions but had a highly significant response to additive stress (Fig. 4G-H; Fig. S8A; Fig. S9; Fig. S10G-I; Kalbfuß *et al*, 2022). The opposite was true for *bin2-3bil1bil2* and *trappii* mutants, which were able to respond to single stress factors but not to additive stress (Fig. 4; Fig. 5; Fig. S11; Kalbfuß *et al*, 2022). Shaggy-like kinases integrate a vast number of signaling pathways (Lv & Li, 2020; Youn & Kim, 2015; Li *et al*, 2021; Song *et al*, 2023; Planas-Riverola *et al*, 2019). Accordingly, we have postulated that the *bin2-3bil1bil2* “decision” phenotype under conflict-of-interest scenarios is primarily due to signal integration (Kalbfuß *et al*, 2022). Prime candidates for the fourth stage, execution or implementation of the action, are *echidna* mutants, which are impaired in the formation of secretory vesicles and cell elongation (Boutté *et al*, 2013; Gendre *et al*, 2013; McFarlane *et al*, 2013); these had highly significant adaptive responses – in the correct orientation – despite their growth defects (Fig. 4E). The question arises as to how to categorize *trappii* mutants in the above conceptual framework. Despite having abnormally short hypocotyls or roots, *trappii* mutants still exhibited differential length increase depending on light availability (Fig. S10). This shows that they are able to perceive and respond to light.

If TRAPPII is primarily involved neither in perception nor in execution, this would place it at the signal integration and/or decision-making steps of adaptive growth decisions. The TRAPPII interactome is vast and complex, with – in addition to the expected ontologies describing traffic, transport and cellular organization – a surprising number of signaling components implicated in responses to abiotic cues (Fig. 1; Fig. S1). This suggests a role for TRAPPII in signal integration. A recent study of the BIN2 signaling network describes the expected components of hormone signaling, phototropism and stress responses; surprisingly, the BIN2 signaling network also includes secretion, endocytosis, autophagy, TGN, endoplasmic reticulum (ER), cell wall and cytoskeleton (Kim *et al*, 2023). Thus, there is a considerable compartmental overlap between the BIN2 network and the TRAPPII interactome. Nonetheless, TRAPPII was not reported among the 482 members of the BIN2 signaling network identified by proximity labeling (Kim *et al*, 2023). The discrepancy could be due to the different methods used: yeast two hybrid and proteomics in this study versus proximity labelling and phosphoproteomics in Kim *et al* (2023). However, a multiomics approach identified AtTRS120-S971 (encompassed by our identified β site AtTRS120-S971:S973:S974:S975) as being differentially phosphorylated in response to brassinolide treatment (Clark *et al*, 2021). This provides an additional line of *in vivo* evidence for a BIN2-AtTRS120 interaction and, consistently with our PPZ experiment (Fig. S6), suggests that TRAPPII phosphorylation is, at least in part, brassinosteroid-regulated.

BIN2 is a negative regulator of BR signaling and AtSK11 and AtSK12 have been shown to negatively regulate root growth responses under osmotic stress (Dong *et al*, 2020; Li *et al*, 2021; Song *et al*, 2023). To explore the biological significance of the AtSK-TRAPPII interaction, we probed the phenotypes of TRAPPII phosphovariants. We found that these had an impact on hypocotyl versus root growth trade-offs in response to additive stress (Fig. 6D-F). Furthermore, the non-phosphorylatable TRS120-SαβγA variant had an enhanced root adaptation whereas the phosphomimetic variant decreased root growth under additive stress (Fig. 6D-F), which is consistent with the negative regulation of growth by BIN2 and other AtSKs. We have previously described plant-specific domains or subunits predicted to be at the dimer interface of the Arabidopsis TRAPPII complex (marked in green in Fig. 2D; Garcia *et al*, 2020; Kalde *et al*, 2019). Interestingly, BIN2 interacts with the plant-specific C-terminal domain of AtTRS120, while MAP65-3 interacts with plant-specific C-terminal domains of the TRAPPII-specific subunits CLUB/AtTRS130 and AtTRS120 (Steiner *et al*, 2016). This presents intriguing implications regarding the potential role of the AtSK-TRAPPII module in meeting the unique demands of endomembrane traffic in plants. Plant and animal cells differ in numerous ways, many of which can be attributed to the presence of the plant cell wall. Novel or expanded families of vesicle-trafficking genes are implicated in cell wall assembly (Assaad, 2001). Cell wall deposition and remodeling underly any growth response in plants. Growth in the face of challenging, restrictive environments is a distinctive survival strategy unique to plants (Assaad, 2001). In this study, we look at the tight regulation of differential growth that enables plants to thrive under multiple stress conditions, in the absence of a carbon or energy source.

The observation that the *trappii* “decision” phenotype was not shared by *echidna* mutants raises the question as to what facet of TGN function is required for responses to additive stress. ECHIDNA and TRAPPII have been shown to have overlapping roles in basal TGN functions such as secretion (Ravikumar *et al*, 2018). In addition, ECHIDNA regulates ER stress and immunity whereas TRAPPII has roles in cytokinesis and in the establishment of cell polarity that appear to be independent of ECHIDNA (Liu *et al*, 2023; Ravikumar *et al*, 2018). In the literature, there are few examples of null mutants that are associated with the TGN and that have a positive influence on growth. One such example is ZmBet5L1 (Zhao *et al*, 2023), which encodes a core subunit of the TRAPPII complex (Kalde *et al*, 2019; Thellmann *et al*, 2010). Null alleles of ZmBet5L1 enhance root growth also in the absence of stress (Zhao *et al*, 2023). The TRAPPII complex has two distinct possible molecular functions. First, it has been postulated to act as a multisubunit tethering complex, even though direct evidence is lacking (Ravikumar *et al*, 2017; Brunet & Sacher, 2014; Kim *et al*, 2016; Pinar *et al*, 2019). Second, it has been shown to act as a Rab GTPase Guanine nucleotide Exchange Factor (Rab-GEF; Cai *et al*, 2005; Morozova *et al*, 2006; Pinar *et al*, 2015; Thomas & Fromme, 2016; Riedel *et al*, 2018). Rab-GEFs act as central cellular switches that can activate Rab GTPase cascades, which are in turn crucial for the development of polarity and directional growth in plants (Elliott *et al*, 2020). Indirect *in vivo* evidence suggests that Arabidopsis TRAPPII acts as a GEF for Rab-A GTPases, orthologues of the RAB11/Ypt31 families known to be activated by fungal and metazoan TRAPPII (Qi & Zheng, 2011; Kalde *et al*, 2019). In Arabidopsis, Rab-A GTPases are TGN-associated and comprise a highly expanded class with 26 members (Kalde *et al*, 2019; Elliott *et al*, 2020). In contrast to ECHIDNA, which is required for the biogenesis of secretory vesicles and for growth generically, as a putative Rab-A GEF TRAPPII has an ability to mediate all the highly diverse facets of TGN function. In a judgement-decision model for plant behavior, judgment is a composite of discrimination, assessment, recognition, and categorization (Karban & Orrock, 2018). In this context, it is noteworthy that the most expanded Rab GTPase family is TGN-associated, suggestive of a tremendous degree of discrimination, specialization and subcompartmentalization at the TGN. Atomic structures of yeast and metazoan TRAPPII depict a conserved triangular structure around the central active site chamber in which TRS120/TRAPPC9 and TRS130/TRAPPC10 form tongs that hold the core complex in place (Galindo *et al*, 2021; Mi *et al*, 2022; Bagde & Fromme, 2022). In a cross-kingdom structural alignment, AtTRS120 and CLUB/AtTRS130 aligned along their yeast orthologues in the same overall structure (Fig. S4). In yeast, TRS120/TRAPPC9 has been proposed to comprise a lid that encloses the active site chamber of the TRAPPII GEF (Bagde & Fromme, 2022). Interestingly, the β and γ phosphorylation sites we have identified in AtTRS120/TRAPPC9 face the active site chamber, including the RAB11/Rab-A binding pocket, proposed by Mi *et al* (2022) and Bagde & Fromme (2022) (Fig. S4A). It is, therefore, tempting to speculate that the phosphorylation status of AtTRS120/TRAPPC9 modulates the specificity of the putative GEF activity of Arabidopsis TRAPPII (Kalde *et al*, 2019).

Our study highlights the possible relevance of the AtSKs-TRAPPII interaction for adaptive growth decisions (Fig. 8). Shaggy-like kinases such as BIN2 integrate a vast number of signaling pathways and, together with receptor complexes at the cell surface, comprise a surveillance system fine-tuned to both biotic and abiotic cues (Lv & Li, 2020; Planas-Riverola *et al*, 2019; Youn & Kim, 2015; Li *et al*, 2021; Song *et al*, 2023). Via the cytosol, the AtSKs would then transmit this information by mediating the phosphorylation status of the TRAPPII complex. TRAPPII would, in turn, mediate TGN function. As an early endosome, the TGN is a central hub in the flow of information to and from the cell surface. It is also intimately connected to the late endosome (or the pre-vacuolar compartment) and to the Golgi, which provides complex polysaccharides as building blocks for the deposition of new cell walls. Thus, the AtSK-TRAPPII interaction would integrate all levels of cellular organization. We posit that signal integration and decision-making occur at the AtSK-TRAPPII interface and that, downstream of TRAPPII, Rab GTPase cascades are implicated in implementing decisions reached at the AtSK-TRAPPII module (Fig. 8). Sorting and trafficking decisions at the TGN would enable plants to respond to developmental or environmental signals via differential growth or various forms of movement.

**Figure 8.**
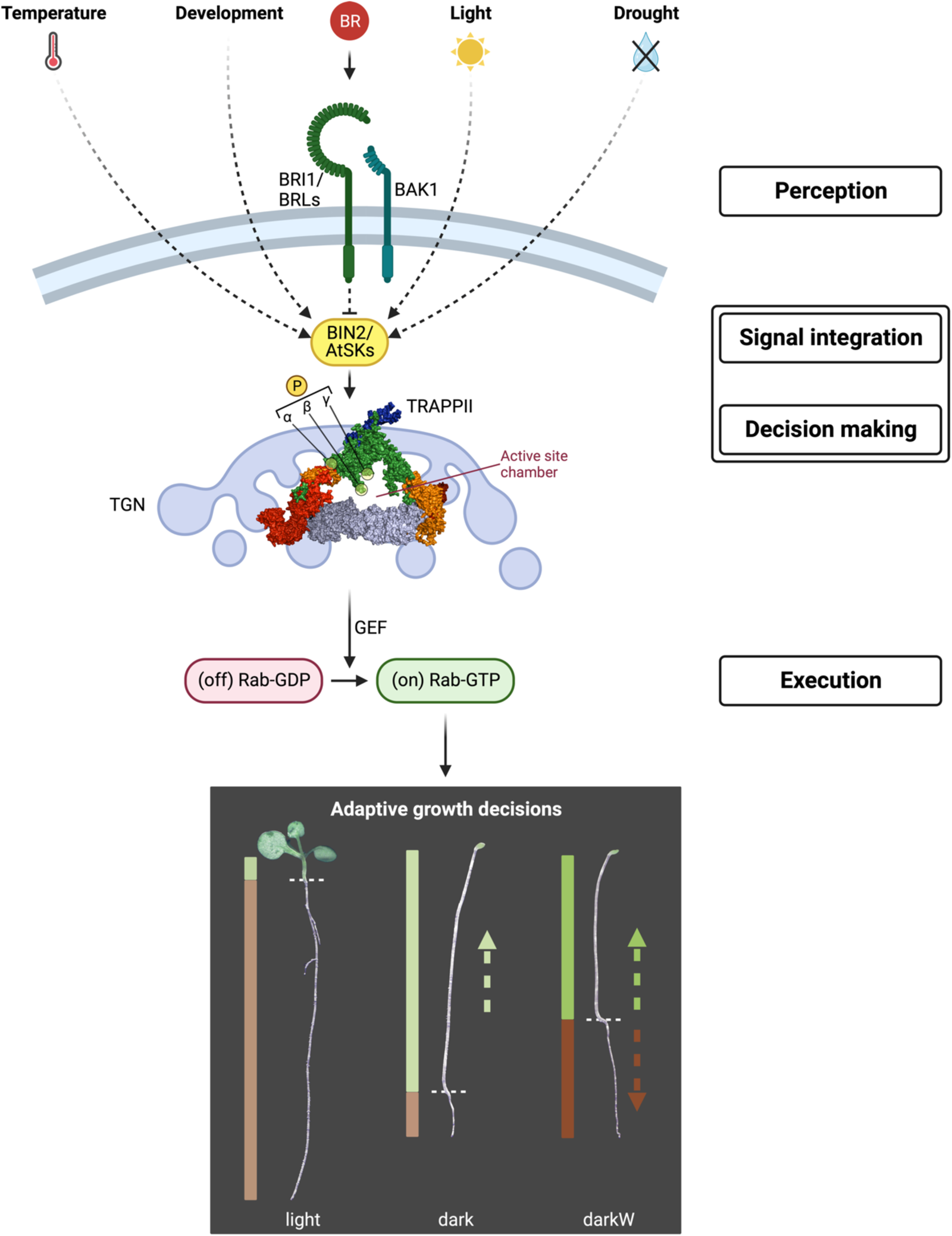
Flow diagram suggesting how BIN2 might mediate plant adaptive responses by phosphorylating the TRAPPII complex. Shaggy-like kinases such as BIN2 integrate a vast number of signaling pathways and, together with receptor complexes at the cell surface, comprise a surveillance system fine-tuned to both biotic and abiotic cues (see text for further detail). BIN2 resides at the plasma membrane (PM) and in the cytosol and TRAPPII resides at the cytosol and in the TGN (Vert & Chory, 2006; Qi *et al*, 2011; Naramoto *et al*, 2014; Rybak *et al*, 2014; Ravikumar *et al*, 2018). We propose that, in the cytosol, the BIN2-TRAPPII interaction serves in the relay of information from the PM to the TGN. We posit that signal integration and decision-making occur at the AtSK-TRAPPII interface. A surface view of the AtTRAPPII complex with AtTRS120 GSK3 phosphorylation sites facing the predicted active site chamber is depicted as AlphaFold structures of AtTRS120/TRAPPC9 and CLUB/AtTRS130/TRAPPC10 mapped onto the cryo-EM-generated structures of yeast TRAPPII (Mi *et al*, 2022; see Fig. S4 for a cross-kingdom structural alignment). Downstream of TRAPPII, Rab GTPase cascades are postulated to be implicated in implementing decisions reached by the AtSK-TRAPPII module. We speculate that the phosphorylation status of AtTRS120 modulates the specificity of the putative TRAPPPII GEF activity, thereby differentially activating distinct Rab GTPase cascades. In the lower panel, we show one example of an adaptive growth decision in 10-day-old seedlings (Col-0 wild type). Bar plots depict hypocotyl (green) versus root (beige-brown) lengths. Light: light-grown seedling with a short hypocotyl and a long root. Dark: a seedling germinated in the dark with a long hypocotyl and a short root. DarkW: Water deprivation in the dark; the hypocotyl is shorter and the root longer than under dark conditions. The dotted green arrow (dark, darkW) represents the incentive for hypocotyl elongation in search of light and the brown arrow (darkW) the incentive for root growth in search of water. Thus, darkW is a “conflict of interest” scenario in which hypocotyl and root growth have competing interests (Kalbfuß *et al*, 2022). Figure generated with BioRender.

Plants exhibit plasticity in their growth, meaning they can adjust their growth patterns in response to environmental cues such as light, temperature, and nutrient availability. It has been argued that plant phenotypic plasticity “is the result of signal integration – a process that requires cell-cell communication, and that results in adaptive forms of movement not to be interpreted as automatic and programmed” (Garzón & Keijzer, 2011). Movement in plants involving organ bending require the polar distribution of morphogens such as auxin. This, in turn, requires the polarized localization of auxin transporters such as PINs. The Arabidopsis TRAPPII complex has been shown to play a pivotal role in a variety of sorting decisions including the polar localization of PIN2 proteins (Qi *et al*, 2011; Rybak *et al*, 2014; Ravikumar *et al*, 2018). How environmental or developmental cues regulate the cargo composition and sorting of TGN vesicles remains unclear. Future experiments will explore the impact of the TRAPPII phosphorylation status on protein sorting, cell polarity, and on the Rab specificity of TRAPPII as a putative GEF. Furthermore, downstream components that govern physiological changes in response to TRAPPII phosphorylation will need to be determined. The presence of the TOR kinase in the TRAPPII interactome is intriguing, as TOR integrates information about nutrient and energy availability, and has recently been shown to regulate actin dynamics as a function of ATP levels (Dai *et al*, 2022); it is tempting to speculate that TOR could provide information on nutrient and energy levels to the AtSK-TRAPPII “decision” module. In plants, the Trans-Golgi Network (TGN) plays a critical role in the relay of information between the cell surface and intracellular compartments. An AtSK-TRAPPII interaction would instruct the TGN, a central and highly discriminate cellular hub, as to how to mobilize and allocate resources to optimize survival under limiting or adverse conditions.

## EXPERIMENTAL PROCEDURES

### Lines and growth conditions

All the lines used in this study are listed in Table S1 (mutant alleles) or in the Supplementary Methods (phosphovariants). Seedling-lethal mutants were propagated as hetero-or hemizygotes. Insertion lines were selected via the TAIR and NASC web sites (Swarbreck *et al*, 2008). Plants were grown in the greenhouse under controlled temperature conditions and with supplemental light, or under controlled growth chamber conditions at the TUMmesa ecotron (16/8 h photoperiod at 180 µmol m^-2^s^-1^). Seeds were surface sterilized, stratified at 4^◦^C for two days and plated on ½ MS medium supplemented with 1% sucrose and B5 Vitamins (Sigma; https://www.sigmaaldrich.com). Plates were incubated at 22°C in constant light (80 µmol m^-2^s^-1^). The root tips of five-day-old plate-grown seedlings were used for confocal microscopy. Seven-day-old plate-grown seedlings were used for co-immunoprecipitation.

### Co-immunoprecipitation (IP)

For CLUB/AtTRS130:GFP we used 3 g of inflorescences per IP experiment. For TRS120:GFP, 3 g of light-grown seedlings were harvested at day 7. For bikinin treatment, seedlings were treated for 30 min with 25 µM bikinin at day 7. For PPZ treatment, 5-day-old seedlings were treated with 2 µM PPZ for 48 h; respective mock treatments were conducted in parallel.

Co-immunoprecipitation experiments were carried out as described previously (Rybak *et al*, 2014). Briefly, seedling lysates were incubated with GFP-trap beads (Chromotek). After washing away all non-binding proteins, 70 µl 2x NuPAGE LDS + 25 mM dithiothreitol (DTT) buffer (ThermoFisher, US) was added and boiled at 70°C for 10 min to denature the bait and all interaction partners. In-gel trypsin digestion was performed according to standard procedures (Shevchenko *et al*, 2006). Briefly, the samples were run on a NuPAGE 4-12% Bis-Tris Protein Gel (Thermofisher Scientific, US) for 3 min. Subsequently, the still not size-separated single protein band per sample was cut out of the gel, reduced (50 mM DTT), alkylated (55 mM chloroacetamide) and digested overnight with trypsin (trypsin-gold, Promega). The resulting peptides were analysed by mass spectrometry. More detailed information on the mass-spectrometric data acquisition is provided in the supporting method section. See data deposition for access to the raw data.

### Molecular techniques and site directed mutagenesis (SDM)

For subcloning for expression in *Escherichia coli* (*E. coli*) or yeast, we used cDNA clones developed by the plant genome project of RIKEN Genomic Sciences Center (Seki *et al*, 1998, 2002). Non-phosphorylatable and phosphomimetic TRS120 phosphosite variant constructs (SαA, SβA, SγA, SαβA, SαγA, SβγA, SαβγA and SαD, SβD, SγD, SαβD, SαβγD)) were generated using the GATEWAY® cloning system and site-specific primers (see supplementary methods for substituted amino acids and used primers). TRS120-T2 cDNA sequences (spanning amino acids 499-1187; Rybak *et al*, 2014) were used as SDM template for protein expression in *E. coli* (kinase assays) and in yeast (Y2H). For *in planta* experiments (confocal microscopy, stress assays), the genomic construct P_TRS120_::TRS120:GFP (Rybak *et al*, 2014) was used as an SDM template. TRS120 phosphovariants were introduced into plants via *Agrobacterium* mediated transformation (see supplementary methods for further details).

### Generation of inducible *trs120* knock-down lines

To generate an inducible *trs120* knock-down mutant, named *trs120i*, an artificial microRNA targeting the 5’ end of AtTRS120 was designed with the Web MicroRNA Designer 3 (Schwab *et al*, 2006). The designed amiRNA sequence (5’-TATAACTCTTACAAGCGGCAT-3’) was introduced in the miR319a precursor and synthesized as a gene strand (Eurofins) with attached attB sites for the GATEWAY® cloning system. The TRS120 amiRNA precursor was cloned into the estradiol inducible pMDC7 vector (Curtis & Grossniklaus, 2003) and introduced into plants via *Agrobacterium* mediated transformation (see supplementary methods for further details and full gene strand sequence).

### Yeast two-hybrid (Y2H)

Y2H pairwise tests were performed as described in Altmann *et al* (2018). Briefly, open reading frames (ORFs) encoding CLUB/AtTRS130 truncations (C2, C3), TRS120 truncations (T1, T3) and BIN2 were transferred by Gateway cloning into the GAL4 DNA-binding domain (DB) encoding Y2H vector pDEST-pPC97, and subsequently transformed into the yeast strain Y8930. These constructs were screened by yeast mating against TRAPPII subunits and truncations thereof (CLUB-C1, -C2, -C3 and TRS120-T1, -T2, -T3) or the TRS120-T2 truncation and its phosphomutants TRS120-T2 SαD, SβD, SγD, SαβD and SαβγD fused to the GAL4 activation domain (AD) in the yeast strain Y8800. Interaction was assayed by growth on selective plates using the *HIS3* reporter, and using 1 mM 3-Amino-1,2,4-triazole (3-AT) to suppress background growth. All candidate interactions were verified by pairwise one-on-one mating in four independent experiments. Only pairs scoring positives in all four assays were considered as *bona fide* interaction partners. The use of low-copy plasmids, weak promoters, the counter-selectable marker cyh2^S^ on the AD-Y plasmid as well as semi-quantitative scoring of quadruplicate tests has been shown to reliably eliminate experimental artifacts and hence false-positives. With the exception of the CLUB-C1 truncation, all TRAPPII truncations and catalytic core subunits used as DB clones in pair-wise tests yielded at least one positive interaction (see also Kalde *et al*, 2019; Garcia *et al*, 2020) and this was used as an internal positive control for the interpretation of negative interaction data.

### *In vitro* kinase assays

Glutathione S-transferase (GST):BIN2 (Li & Nam, 2002) and GST:TRS120-T2 WT or phosphovariants were expressed in *Escherichia coli* (Rosetta-gami^TM^ strain) under constant shaking for 20 h at 25°C or 20 h at 18°C, respectively. The expressed proteins were affinity purified with GST-tags. After sonication of the samples in 1x PBS, 1 mM PMSF, 1 mg/ml lysozyme and 1% Triton X-100, the bacterial cell rests were centrifuged for 30 min at 16,000 g. Supernatants were incubated for 2 h with Glutathione Sepharose^®^ 4 Fast Flow Beads (GE Healthcare) while rotating. After washing the samples five times, the GST tagged proteins were eluted with 10 mM glutathione and concentrated by ultra-filtration.

***In vitro*** kinase assays with radiograph readout were performed as described previously (Kim *et al*, 2012), with some adaptations. In short, ca. 0.1 µg GST:BIN2 and ca. 0.5 µg of GST:TRS120-T2 WT or phosphovariant were incubated in 20 mM Tris pH 7.5, 1 mM MgCl_2_, 100 mM NaCl, 1 mM DTT, 100 µM ATP and 10 µCi ATP[γ-^32^P] for 3 h at 22°C in motion. The kinase reaction was stopped by adding 2x SDS sample buffer and by boiling the samples for 10 min. Protein phosphorylation was analyzed by autoradiographs and SDS-PAGE.

***In vitro*** kinase assays with mass-spectrometry readout were performed to determine specific phosphorylation sites. For each reaction, 10 µg of substrate (TRS120-T2) and different dilutions of the kinase (undiluted, 1:5, 1:10, 1:20 or 1:100 as indicated in the figure panels) were incubated for 15, 30, and 120 min in a kinase buffer (20 mM Tris HCl pH 7.8, 100 mM NaCl, 1 mM MgCl_2_, 1 mM DTT, 1 mM ATP). For the negative controls, one sample with the highest kinase concentration and the longest incubation time was incubated in a kinase buffer without ATP. For the kinase dead control, the kinase was heat-inactivated prior to the incubation with its substrate. To stop the reaction, samples were heated at 95°C for 5 min. After reduction (10 mM DTT), alkylation (55 mM CAA) and protein digestion (trypsin-gold, Promega) the resulting peptides were purified using self-packed StageTips (C18 material, 3M Empore) and analyzed by mass spectrometry. More detailed information can be found in the supporting methods.

### LC-MS/MS data acquisition and analysis

Generated peptides were analyzed on a Dionex Ultimate 3000 RSLCnano system coupled to a Q-Exactive HF-X mass spectrometer (Thermo Fisher Scientific) in data-dependent acquisition mode. Peptide identification and quantification was performed using the software MaxQuant (version 1.6.1.0) (Cox & Mann, 2008; Tyanova *et al*, 2016a) with its built-in search engine Andromeda (Cox *et al*, 2011) and an ***Arabidopsis thaliana*** reference database (Araport). Perseus (Tyanova *et al*, 2016b) and Python 3.0 were used for statistical data analyses. For a targeted analysis of the kinase assay data set, MS1 chromatograms from selected phosphopeptides were extracted and analyzed using the Skyline software (MacLean *et al*, 2010). For detailed information about LC-MS/MS data acquisition and analysis see supporting methods.

### Light and electron microscopy

For scanning electron microscopy, a Zeiss (LEO) VP 438 microscope was operated at 15 kV. Fresh seedlings were placed onto stubs and examined immediately in low vacuum. Confocal microscopes used for imaging were an Olympus (www.olympus-ims.com) Fluoview 1000 confocal laser scanning microscope (CSLM) and a Leica (www.leica-microsystems.com) SP8 Hyvolution CSLM. 40x and 60x water immersion 0.9 numerical aperture objectives (Olympus) were used. Imaging data were acquired using LAS-X software (Leica) and FV10-Ver.2.6.0 software (Olympus). GFP fluorescent proteins were imaged with 488 nm excitation and 500-550 nm emission. 10-day-old seedlings were stained with modified pseudo-Schiff propidium iodide staining (Truernit *et al*, 2008). Excitation Laser was set to 488 nm. The Emission was detected at 520 nm. The junction between the meristematic and elongation zones was determined by marking the first cell in a single cortex cell file that was double the length of the previous cell (González-García *et al*, 2011). For seedling images, a Leica S APO stereo microscope with a Leica MC170 HD camera was used.

### Differential growth decision assay

The differential growth decision experiment was performed as described by Kalbfuß *et al* (2022). In brief, seeds were surface sterilized using a brief 80% ethanol rinse followed by 15 min incubation in sterilization buffer (0.01% SDS, 3% NaOCl). After 5 washes in mQ water, seeds were resuspended in 0.15% agar and stratified in the dark at 4°C for 7 days to break seed dormancy. Afterwards, seeds were plated on squared plates with exactly 45 ml ½ MS medium supplemented with B5 Vitamins (Sigma). Culture media for TRS120 phosphomutants was additionally supplemented with 50 μg/ml kanamycin to select for transgenic seedlings. For -0.4 MPa water deficit, media plates were infused with sterile-filtrated 45 ml 2x PEG-6000 (Merck group; dissolved in liquid ½ MS media) for exactly 24 h at RT and afterwards the PEG-solution completely decanted. Seeds were plated at the interface between the culture media and sterilized plastic strips on the media such that, upon germination, only the root touched the agar. For water stress, seeds in each biological replicate were sown on two plates for technical replicates which were pooled for analysis. After plating, plates were sealed with breathable tape. Plates for dark conditions were wrapped with two layers of thick aluminum foil. All plates were incubated for 10 days at 22°C with permanent light (80 μmol m^-2^s^-1^). Note that the plates were negatively inclined by 4° to promote root growth on the media surface rather than in the agar.

After incubation of 10 days seedlings were transferred onto cold 1.2% agar plates, scanned at 1200 ppi and saved as tiff files. Hypocotyl and root lengths were analyzed with Fiji – ImageJ using the free-hand-tool. The ratio was calculated as hypocotyl length divided by root length.

Root and hypocotyl responses to water deficit in the dark (abbreviated as darkW) were computed as

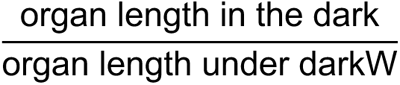

Because the hypocotyl and root have opposite responses to our additive stress conditions (hypocotyl length decreases but root length increases in response to water stress in the dark, as shown in Fig. 4) the RQs move in opposite directions.

The ratio was computed as follows:

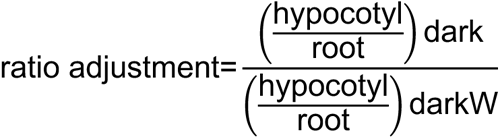

Due to variability between PEG lots and PEG plates (Kalbfuß *et al*, 2022), we normalized each mutant to the corresponding wild-type ecotype (see Table S1) on the same plate. Thus, the normalized ratio adjustment to water stress in the dark (darkW) was computed as follows:

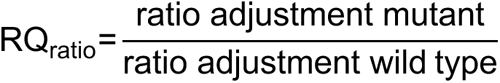

RQ_hypocotyl_ and RQ_root_ as well as light versus dark responses were computed in a similar fashion (Fig. S9, S10, S14). These normalized organ and ratio adjustments were referred to as response quotients (RQ) to light-to-dark or dark-to-darkW conditions. The mean RQ_ratio_ of at least three biological replicates (i.e. the seed stocks from different mother plants) is shown. Please see Supporting Table S3 for thresholds for attenuated, normal, versus enhanced organ or ratio responses. For volcano plots we plotted the mean RQ_ratio_ on the X axis and the median *P*_ratio_-value on the Y axis. Organ and ratio adjustments for light-to-dark comparisons were computed as organ length (or ratio) in the light divided by organ length (or ratio) in the dark. *P*-values were computed with the student’s t-test. Responses were considered to be insignificant for *P*-values ≥ 0.05 and attenuated for *P*-values ≥ 0.00001. The median *P*-value for at least three replicates is shown in the volcano plots.

### Germination assay under osmotic stress

For germination curve experiments, surface-sterilized seeds were stratified in the dark at 4°C for two days. Seeds were plated on ½ MS media supplemented with 0.05% B5 Vitamins (Sigma) and either 0 mM, 200 mM, or 400 mM mannitol (Millipore). Approximately 100 seeds were plated per condition. Plates were incubated at 22°C under constant light conditions (80 μmol m^-2^s^-1^) and rotated every day to avoid light effects. Seed germination rates were recorded every 24 h. Seeds were scored as germinated when the radicle emerged. Data were obtained for three biological replicates, each with three technical replicates.

### Root gravitropism assay

*trs120i* lines were selected on ½ MS media supplemented with 20 µg/ml hygromycin B. Control lines were grown in parallel on media without antibiotics. After 4 days, seedlings were transferred to ½ MS media supplemented with 20 µM β-estradiol (Est; 20 mM stock dissolved in EtOH; Sigma) or 0.1% EtOH as mock control and grown vertically for additional nine days. The primary root tip angles (excluding lateral roots) were analyzed with Fiji – ImageJ. Primary roots exhibiting looping (see yellow arrow in Fig. 7B) were scored as -180° due to their growth against gravity. Primary root growth orientation was visualized as frequency plots in a polar coordinate system in R. To quantify agravitropic roots, the frequency of primary root tips growing against gravity (angles < – 90° and > 90°) was counted per experiment.

### Western blot analysis

Standard Western blot analysis was performed according to Sambrook *et al* (1989). 7.5% SDS-PAGE gels (Mini-PROTEAN TGX gels, Bio-Rad) were blotted onto PVDF Immobilon-FL membranes (Millipore). Polyclonal rabbit anti-GFP antibody (1:2000; A11122, Invitrogen) was used as primary antibody. As secondary antibody, we used a goat anti-rabbit Horseradish peroxidase-conjugated antibody (1:6000; Pierce, Thermo Scientific) together with the SuperSignal West Femto Maximum Sensitivity Substrate (Thermo Scientific). To check for equal loading, blots were stained with Coomassie (10% acetic acid, 50% methanol, 0.25% Coomassie Blue R-250) and subsequently destained (10% acetic acid, 50% methanol).

### Data analysis and image processing

False discovery rates, determined with the standard two-tailed t-test, were set at a cut-off of 1. Images taken with the Leica SP8 microscope were deconvolved using the built-in Huygens Scientific deconvolution software (www.leica-microsystems.com) operated in 2D. For consistency, we selected cortical root tip cells, at a height of 6-22 cells above the quiescent center in the root apical meristem. Images were processed with Adobe photoshop (www.adobe.com) and GIMP (https://www.gimp.org) and analysed with Image J (https://imagej.nih.gov). Statistical analyses were performed in RStudio. Figures were created with RStudio’s ggplot2 package and assembled with Inkscape (https://inkscape.org). Please see the supporting methods for the analysis of mass spectrometry datasets and for multiple sequence and structural alignments.

## Supporting information

Rebuttal

## Acknowledgements

We thank Prof. Wilfried Schwab, Prof. Erwin Grill and other members of the botany department for their support. Thanks to Prof. Arne Skerra, Martin Schlapschy and Veder Garcia for useful suggestions. We thank Yannik Schreckenberg, Andreas Czempiel, Theo Kalmbach, Hermine Kienberger, Nina Lomp and Franziska Hackbarth for technical assistance. We thank the WZW/TUM Centre for Advanced Light Microscopy (CALM), headed by Ramon Torres-Ruiz and Klaus Michel, for access to confocal microscopes, and Prof. Kay Schneitz for access to their Leica S APO stereo microscope. Thanks to Roman Meier at the TUMmesa facility, directed by Leonardo Teixeira and Bálint Jákli, for supporting us with optimal growth conditions for our plants. Jorge José Casal, Sean Cutler and Ueli Grossniklaus shared published resources.

## Data deposition

The CLUB:GFP IP-MS data has previously been published (Kalde *et al*, 2019) and deposited to the ProteomeXchange Consortium via the Proteomics IDEntification (PRIDE) partner repository (Perez-Riverol *et al*, 2019). The dataset can be accessed with the identifier PXD013016. All other IP-MS or MS datasets are being deposited here for the first time. New proteomics raw data (*in vitro* kinase assays and *in vivo* evidences with AtTRS120:GFP IP-MS of bikinin and PPZ treated seedlings) including MaxQuant search results and used protein sequence databases can be accessed via Panorama Public (Sharma *et al*, 2014, 2018) with the following link https://panoramaweb.org/phosphoTRAPP_Ara.url.

## Author contributions

Conceptualization: FFA

Methodology: MA, CL, BA, CHP, PFB, CW, AS, NK, ASM, BB, CM, EF, FFA

Investigation: CW, MA, BA, MAM, AS, NK, ASM, RR, CHP, BB, CM, EF, FFA

Visualization: CW, MA, BA, MAM, AS, NK, ASM, RR, CHP, BB, CM

Funding acquisition: DWE, PFB, ZW, CL, FFA

Project administration: FFA

Supervision: EF, DWE, PFB, ZW, CL, FFA

Writing – original draft: FFA, CW

Writing – review & editing: CW, MA, BA, NK, ASM, RR, CHP, BB, CM, PFB, ZW, CL, FFA

