## Supplementary material for "Regulation of adaptive growth decisions via phosphorylation of the TRAPPII complex in Arabidopsis": Rebuttal

### Full Revision

Review COMMONS

**Manuscript number:** RC-2023-01983
**Corresponding author(s):** Farhah, Assaad

#### General Statements

We are hereby submitting a very thoroughly revised manuscript. We wish to thank the reviewers for their useful suggestions, all of which have been implemented. New approaches or lines of evidence include pharmacological inhibition, an inducible *trappii* allele, genetic interaction data, and cross-kingdom structural alignments. New results are presented in Fig. 7, Fig. S4, Fig. S6, Fig. S8, and Fig. S11B-D. The references have been extended. We have also streamlined the text and worked on its clarity and readability for a broad audience, with a special focus on the figure legends.

#### 1. Point-by-point description of the revisions

Please find below a point-by-point response (in black) to the reviewer comments (in blue).

**Reviewer #1 (Evidence, reproducibility and clarity (Required)):**

Universally present across the eukaryotic world, TRAPPII is a hetero-oligomeric complex that plays a key role in the regulation of the TGN that has often been classified as a multisubunit tethering complex, although conclusive evidence for the tethering role is still lacking. In contrast, it is well established that the complex acts as a GEF for RAB11, primarily by studies carried out with model fungi such as Saccharomyces cerevisiae and Aspergillus nidulans. Atomic structures have revealed that a few amino acids in one of the subunits of the complex suffice to "kick off" GDP from the active site of the substrate RAB. While some of the subunits are necessary to place the RAB nucleotide binding pocket at the right distance from the key TRAPPII residues on the target membrane, it seems unlikely that the sole function played by this Md complex is catalysing the exchange of GDP by GTP in the target GTPase. In this particular regard, our understanding of other potential physiological roles of the complex is utterly incomplete

This very well written manuscript explores the regulation of TRAPPII by phosphorylation, more precisely, the role of phospho-sites in the regulation of TRS120/TRAPPC9. Using co-immunoprecipitation strategies coupled to mass spectrometry, the authors identified a member of the saggy/GSK family of kinases. Given that the interaction of protein kinases with their substrates is supposed to be transient, this is technically sound result that testifies to the impeccable methodology used by authors. They further exploit MS methodology and two hybrid analysis to identify region of TRAPPC9 interacting with the kinase, as well as the phosphorylated residues. The authors close the circle by establishing that substitutions of these residues result in modified responses to abiotic stresses. Thus, a significant merit of this work is opening up the can of regulation by phosphorylation of TRAPP functions.

Experiments/challenges for the future are determining the downstream components that govern these physiological changes in response to phosphorylation of TRAPPII, and expanding these phosphorylation studies to TRAPPIII. This latter complex is involved both in exocytosis and in

autophagy, and it seems plausible that alternation between these two different fates is governed by post-translational modifications of the type studied here.
My suggestions to the authors: there has been a burst of atomic structures of TRAPPs recently, with three papers authored by the Fromme, Munro and Sui labs. It mighty worth comparing the longer Trs120 in these structures with the prediction of the shorter protein from Arabidopsis.
**Response:** We thank the reviewer for pointing this out. We have superimposed the AlphaFold structure of AtTRS120/TRAPPC9 onto the structures from Sui (Monomer) and Fromme (Holodimer). This is shown in Fig. S4 (see also Results p. 7). We write:
Discussion p. 16
"Atomic structures of yeast and metazoan TRAPPII depict a conserved triangular structure around the central active site chamber in which TRS120/TRAPPC9 and TRS130/TRAPPC10 form tongs that hold the core complex in place (Galindo *et al*, 2021; Mi *et al*, 2022; Bagde & Fromme, 2022). In a cross-kingdom structural alignment, AtTRS120 and CLUB/AtTRS130 aligned along their yeast orthologues in the same overall structure (Fig. S4). In yeast, TRS120/TRAPPC9 has been proposed to comprise a lid that encloses the active site chamber of the TRAPPII GEF (Bagde & Fromme, 2022). Interestingly, the β and γ phosphorylation sites we have identified in AtTRS120/TRAPPC9 face the active site chamber, including the RAB11/Rab-A binding pocket, proposed by Mi *et al* (2022) and Bagde & Fromme (2022) (Fig. S4A). It is, therefore, tempting to speculate that the phosphorylation status of AtTRS120/TRAPPC9 modulates the specificity of the putative GEF activity of Arabidopsis TRAPPII (Kalde *et al*, 2019)."

A very minor point is that possibly the most thorough report on the localisation of TRAPPII to the TGN is that co-signed by Mario Pinar and myself in the Journal of cell science
**Response:** We now also cite Mario Pinar and Miguel A. Peñalva, Journal of Cell Science, 2020 https://doi.org/10.1242/jcs.241141.

**Referees cross-commenting**

I have nothing to add to my review, which was made from the point of view of TRAPP researcher. if my colleagues understand that the manuscript is hyperbolic in places, over statements should be removed

Reviewer #1 (Significance (Required)):

TRAPPII is regulated by TRAPPC9 phosphorylation in cruciferae. Convincing evidence. Impeccable presentation and writing.
**Response:** We thank the reviewer for his/her positive assessment, and for his/her TRAPPII perspective, which we have in part incorporated into the introduction and perspective.

**Reviewer #2 (Evidence, reproducibility and clarity (Required)):**

Summary

This manuscript adopted the concept that plants have cognitive ability and proposed the hypothesis that the trans-Golgi network plays a role in the decision process of cells. It investigates how this organelle function in order to reach the right decision when exposed to the combined drought stress and dark or to germination under osmotic stress. The study tests the hypothesis focusing on the on the TRAPPII complex. The authors demonstrated that defects (mutations) on TRAPPII complex cause wrong growth decisions, particularly when seedling are exposed to decisions with trade-offs.

Major comments

The experiments made for this study are enough to support the conclusions, and they were performed with adequate procedures. I just make the comment that follows.
Lines 468-472: The authors propose that "signal integration and decision-making occur at the AtSK-TRAPPII interface". It should be considered whether the decision is made in a specific step and location or if all the cascade of responses is the decision process. It is proposed that TRAPPII makes the decision and the Rab GTPase cascades (or the downstream signals) implement the decision. The authors demonstrated that a defect on TRAPPII causes wrong-decisions, but what would happen if TRAPPII were normal but something downstream was defective and could would proceed the regular process? Would it also lead to wrong growth decisions? I am afraid that any defect may cause wrong decisions because the decision is the full metabolic process and not a single step in the route.

**Response:** We have not performed allocation decision experiments with Rab GTPase mutants as these are typically dominant alleles (dominant negative or constitutively active). However, we have performed allocation decision experiments with perception mutants (*phyaphybcry1cry2* and *pyrpyl* quadruple mutants) and with *echidna*, which we consider to be an “execution” mutant. In contrast to decision mutants (which are characterized by an inability to adapt to additive stress or have opposite responses to stress as compared to the wild type), we observe compromised but overall correct adaptation to additive stress in execution mutants. This is described as follows:

<u>Results p. 9</u>

“We … compared *trappii* mutants to *echidna* mutants, severely impaired in TGN structure and function (Boutté *et al*, 2013; Gendre *et al*, 2013; McFarlane *et al*, 2013). Both *echidna* and *trappii* mutants exhibited a severe impairment in root elongation in the light (Fig. S10C-F; Fig. S10H). In contrast to *trappii*, however, *echidna* mutants had highly significant responses to additive stress that resembled the wild type in all respects (Fig. 4E-H; Fig. S8F). Thus, *echidna* mutants do not qualify as decision mutants.”

<u>Discussion p. 14</u>

“Prime candidates for the fourth stage, execution or implementation of the action, are *echidna* mutants, which are impaired in the formation of secretory vesicles and cell elongation (Boutté *et al*, 2013; Gendre *et al*, 2013; McFarlane *et al*, 2013); these had highly significant adaptive responses - in the correct orientation - despite their growth defects (Fig. 4E).”

**Response:** we write
Results on p. 9
"A question that arises is whether *trappii* mutants are impaired in differential growth decisions as a secondary consequence of primary defects in morphogenesis or cytokinesis (Jaber *et al*, 2010; Rybak *et al*, 2014; Thellmann *et al*, 2010). The etiolation response was severely attenuated in *trappii* mutants, but nonetheless highly significant (P < 0.00001; Fig. S10C-D, S10F-I). Similarly, an attenuated but clear etiolation response has been shown for other cytokinesis-defective mutants including *keule* and *knolle* (Assaad *et al*, 2001). This shows that, despite a severe impairment in cell division and morphogenesis, cytokinesis-defective mutants are nonetheless capable of differential growth."

Line 413: Correct Kim et al (2023).
**Response:** We thank Reviewer 2 for pointing out the wrong citation style. We changed the reference to "Kim *et al* (2023)".

Reviewer #2 (Significance (Required)):
This study offers an important contribution to the discussion on how plants make decisions. The signals and the cascade of stimuli flows through an intricate network. This study demonstrates that on of the streams of information used for growth decisions passes through the Golgi complex.
**Response:** We thank the reviewer for his/her positive assessment.

**Reviewer #3 (Evidence, reproducibility and clarity (Required)):**

Summary:
In the manuscript "Regulation of adaptive growth decisions via phosphorylation of the TRAPPII complex in Arabidopsis" the authors investigate the TRAPPII interactome carried out by an already published IP-MS screen. They study previously identified shaggy-like kinases SK as TRAPII interactors and the phosphorylation sites by Y2H (interactions of wild type, deletion mutants and phosphomutants) and kinase assays (in vitro) and pharmacological inhibition in the subunit AtTRS120. The authors provide a deeper phenotypical analysis of trapii null mutant lines and classification as "decision mutants", based on "limited budget" and "conflict of interest" experiments (previously described) as a starting point of investigations of TGN function in comparison with hormone mutants. Cell elongation is used as a response phenotype. Authors focus on mainly TRS120 and phosphorylation by SK and partly on another TRAPP component, CLUB. Authors study the assays with differing kinases, e.g. Y2H with BIN2, phosphorylation with SK11.

Major comments:
A major issue is that new claims and conclusions are not supported by the new data provided here. The title says "in Arabidopsis", but only components are from Arabidopsis. Interactions,

instead, are studied in this manuscript by Y2H in yeast, and phosphorylation in vitro. The abstract is very misleading and does not distinguish which aspects are studied in vitro and which in vivo.

**Response:** We have extended the *in vivo* parts of the data to include more pharmacological inhibition and genetic interaction data. We specify in the results whether the data are *in vitro* or *in vivo/in planta*. *In vivo* data (Table R1) include IP-MS, pharmacological inhibition, mutant analysis, genetic interactions and the analysis of phosphovariants *in planta*.

**Table R1: *in vitro* versus *in vivo* lines of evidence.** In bold are new lines of evidence not included in the initial submission.

| ***in vivo* evidence** | **Figure** | ***in vitro* evidence or heterologous system** | **Figure** |
|---|---|---|---|
| IP-MS TRAPPC10/CLUB:GFP (AtSK11/12/32) | 1B | Y2H, with wild-type sequences and phosphovariants (BIN2-AtTRS120-T2) | 2 |
| IP-MS TRAPPC9/AtTRS120:GFP (AtSK1; AtSK2; AtSK3; AtSK4) | S2, S3B-D | kinase assay with $^{32}P$ (BIN2-AtTRS120-T2; AtSK11 AtTRS120-T2) | 3A, S5A |
| IP-MS TRAPPC10/CLUB:GFP light-specific interactions not seen in the dark (AtSK11/12) | 1C, S1B | kinase assay with mass spectrometry read out (AtTRS120-T2 SK11, BIL2, SK32, **BIN2**) | 3B, S5B |
| IP-MS TRAPPC9/AtTRS120:GFP with pharmacological inhibition: bikinin and **PPZ** | S6 | | |
| presence of related BR-pathway components in TRAPPII interactome | 1C | | |
| related phenotypes of *trappii* alleles (including ***trs120i)*** and *bin2-3bil1bil2* as (i) "confused" decision and (ii) **agravitropic** mutants | 4, 5, 7 | | |
| opposite phenotypes of non-phosphorylatable versus phosphomimetic variants in physiological and/or growth assays *in planta* | 6 | | |
| **synergistic genetic interaction (*trs120i* with *bin2-3bil1bil2*)** | 7 | | |
| overlapping localization of BIN2 and TRS120 in the cytosol | R1 | | |

New text to describe the new findings include:

<u>Results p. 7</u>

"… seedlings treated with the BR biosynthesis inhibitor PPZ, which should relieve BR-mediated BIN2 inhibition, showed an increased extent of phosphorylation of the TRS120-γ peptide (Fig. S6)."

<u>Results p. 12</u>

"To further address the physiological significance of the BIN2-TRAPPII interaction *in vivo*, we deployed higher order mutant analysis. This approach, however, was challenged by (i) the functional redundancy between BIN2 and its homologues (Vert & Chory, 2006; Yan *et al*, 2009), (ii) the semi-dominant nature of *bin2-1*, (iii) the seedling lethality of *trappii* null alleles, as well as (iv) the pleiotropic phenotypes of both *bin2-1* and *trappii* dwarfs (Li *et al*, 2001; Thellmann *et al*, 2010; Ravikumar *et al*, 2018; Garcia *et al*, 2020). To address these challenges, we engineered a conditional *trs120* knock-down allele, named *trs120i*. This features an artificial microRNA that targets 5' AtTRS120/TRAPPC9 sequences, expressed under an estradiol-inducible promoter (Fig. 7A; Curtis & Grossniklaus, 2003). The *trs120i* construct was introduced into the wild type (Col-0) and into the *bin2-3bil1bil2* triple knock-out mutant. Upon induction, *trs120i* exhibited a mild and *bin2-3bil1bil2* a more pronounced root agravitropism (Fig. 7B-C). The *bin2-3bil1bil2 trs120i* higher order mutant had a more than additively enhanced agravitropic response upon induction as compared to *trs120i* or *bin2-3bil1bil2* alone (Fig. 7B-C, 7E). This was evidenced as primary roots growing in all directions, often against gravity at 180° in *bin2-3bil1bil2 trs120i* (Fig. 7B-C). We observed synthetic enhancement specifically upon estradiol induction and not in the mock control (Fig. 7D, compared to 7E). This is indicative of a synergistic genetic interaction. Furthermore, it suggests that adaptive growth decisions such as gravitropism are mediated by the BIN2/AtSK-TRAPPII interaction."

The Abstract does not mention that this study is based on previously identified interactome data.

**Response:** The CLUB:GFP interactome data (Fig. 1B) was indeed already published in Kalde *et al* (2019). We state this, together with some first novel aspects, in the first paragraph of the results (p. 5) as follows:

"To gain insight into TGN function, we used the TRAPPII interactome as a starting point. We have previously identified TRAPP and EXOCYST subunits, cytoskeletal proteins and Rab GTPases in the TRAPPII interactome (Rybak *et al*, 2014; Steiner *et al*, 2016; Kalde *et al*, 2019; Garcia *et al*, 2020). However, no global meta-analysis of the vast Arabidopsis TRAPPII interactome has been carried out to date."

In contrast, the TRS120:GFP interactome and phosphostatus described in this manuscript have not been published previously and represent data specific to this manuscript. We have clearly described the deposition of data for the different manuscripts in the data deposition section (p. 23) as follows:

"The CLUB:GFP IP-MS data has previously been published (Kalde *et al*, 2019) and deposited to the ProteomeXchange Consortium via the Proteomics IDEntification (PRIDE) partner repository (Perez-Riverol *et al*, 2019). The dataset can be accessed with the identifier PXD013016. All other IP-MS or MS datasets are being deposited here for the first time."

o The figure legends are often not sufficiently detailed to understand what exactly is represented. Therefore, it is not possible to judge in every case whether experiments are supported by data. E.g.

Fig. 1: A, describe which data were used and which control for IP-MS had been taken into account. B, this is a plot, please describe what is represented. Explain better why Shaggy kinases were chosen. C, explain the principle and what is represented. How is this experiment controlled and how is it ensured that negative results are not caused by absent proteins.

**Response:** The figure legend (see p. 29) specifies that:

- CLUB:GFP IP-MS data from inflorescences were used.
- The empty vector cassette (soluble GFP) was used as a negative control.
- A volcano plot is represented in Fig. 1B. On the X axis: The ratio (or fold-change) was calculated for each protein as the average intensity of the signal in the experiment divided by its average intensity in the negative control. The Y axis depicts the *P*-values of the signal in the experiment versus the control, based on three biological replicates and depicted along a negative log10 scale.

Shaggy-like kinases were chosen because they were the only positive TRAPPII-Kinase interaction identified in pairwise Y2H tests, as specified in the legend to Fig. 2B (see p. 30). For the Y2H (previously Fig. 1C) please see our response to the next reviewer comment on Fig. 2.

Fig. 2: Indicate the phosphorylation sites in the other subfigures. Fig. 2E: How was it generated, explain what is seen. Since this is the only figure illustrating the protein complex of TRAPP, this figure should be more thoroughly prepared and labeled. I recommend a better visualized protein complex. As before, Fig. 2F remains unclear.

**Response:** We have now labelled all the subunits of the TRAPPII complex. The model shows a TRAPPII dimer and highlights the plant-specific sequences, including the TRIPP subunit, in green. Structural models of yeast and metazoan TRAPPII monomers and dimers confirm our Arabidopsis dimer model, which was generated based on extensive pairwise Y2H tests with the Arabidopsis core and TRAPPII-specific subunits (Kalde *et al*, 2019; Garcia *et al*, 2020). The negative controls for Y2H are the empty vector DB cassettes tested against the different AD constructs. As positive controls, we used TRAPPII-specific subunits, known to interact with TRS120-T2 (Kalde *et al*, 2019). This is specified in the figure legend (see p. 29-31).

The hybrid proteins for Y2H are expressed from low copy CEN vectors and using the weak minimal ADH2 promotor. These features are intended to increase the quality of interaction identification by preventing protein aggregation due to high expression levels; at the same time, however, also due to the small AD domain, we are not able to detect proteins by Western blot. Nonetheless, all DB clones reported in the study have positive interactions in parallel experiments (Table R2), with the exception of CLUB-C1, which spans the first 400 amino acids of the protein. This shows that the constructs are expressed.

We have now added the following to the text:

Methods p. 19

"The use of low-copy plasmids, weak promoters, the counter-selectable marker $cyh2^S$ on the AD-Y plasmid as well as semi-quantitative scoring of quadruplicate tests has been shown to reliably eliminate experimental artifacts and hence false-positives. With the exception of the CLUB-C1 truncation, all TRAPPII truncations and catalytic core subunits used as DB clones in pair-wise tests yielded at least one positive interaction (see also Kalde *et al*, 2019; Garcia *et al*, 2020) and this was used as an internal positive control for the interpretation of negative interaction data."

**Table R2: Internal positive controls in our Y2H dataset.** The strength of the interaction in quadruplicate tests is scored semi-quantitatively; in the table the number of + signs depict the strength of the interaction. See Fig. 2F. TRS120-DB auto activates (red text). T1, T2 and T3 refer to TRS120 and C1, C2 and C3 to CLUB truncations.

| | Y2H AD clones | | | | | | | | Reference |
|---|---|---|---|---|---|---|---|---|---|
| | Bet3 | Trs23 | Trs31 | Tca17 | T2 | C2 | TRIPP | EMPTY | |
| DB | | | | | | | | | |
| Bet5 | | +++++++ | | | | | | | Kalde *et al*, 2019 |
| Bet3 | | | ++ | | | | | | Kalde *et al*, 2019 |
| Trs31 | | +++ | | | | | | | Kalde *et al*, 2019 |
| Tca17* | | | | | | + | | | Kalde *et al*, 2019 |
| Trs33 | +++++ | | | +++++ | | | | | Kalde *et al*, 2019 |
| T1 | | | | | +++ | | +++ | | Kalde *et al*, 2019; Garcia *et al*, 2020 |
| T2 | | | | | | | | ++ auto-activation | Steiner *et al*, 2016 |
| T3 | | | | | +++ | | +++ | | Kalde *et al*, 2019; Garcia *et al*, 2020 |
| C1 | | | | | | | | | |
| C2 | | | | | +++++++ | | +++++++ | | Kalde *et al*, 2019; Garcia *et al*, 2020 |
| C3 | | | | | +++++++ | | +++++++ | | Kalde *et al*, 2019; Garcia *et al*, 2020 |

Fig. 3: Please add a figure illustrating the mutations. 3C: what has been diluted? Other examples are found in other figures.

**Response:** The mutations are now depicted in red in Fig. 2C. We now specify in the figure panel and in the legend that the kinase was diluted (Fig. 3B; previously Fig. 3C).

Fig. 4: Shouldn't the wild type be compared with all the mutants? Then statistics have to be conducted accordingly. Better explain G and H. If there are quotients, explain of what exactly.

**Response:** A direct comparison between the mutant and wild type is now provided in violin plots in Fig. S8 (new). The response quotients (RQs) allow us to compare mutants in different genetic backgrounds and are now explained more thoroughly, as follows:

Results p. 8

"In addition to comparing organ lengths, each mutant line was normalized to its corresponding wild-type ecotype on the same (PEG) plate, which helped us to take the variability between PEG plates and experiments into account and enabled us to pool biological replicates (see methods for further detail). To this end, the response to water stress in the dark was represented as a normalized response quotient (RQ). The RQ is an indication of how well each mutant responds to a given combination of stress cues and indicates how much each line deviates from the wild type. A value of 1.0 means that the mutant line behaves exactly like its corresponding wild type."

Methods p. 21

"… the normalized ratio adjustment to water stress in the dark (darkW) was computed as follows:

$$RQ_{ratio} = \frac{\text{ratio adjustment mutant}}{\text{ratio adjustment wild type}}$$

$RQ_{hypocotyl}$ and $RQ_{root}$ as well as light versus dark responses were computed in a similar fashion (Fig. S9, S10, S14). These normalized organ and ratio adjustments were referred to as response quotients (RQ) to light-to-dark or dark-to-darkW conditions. The mean $RQ_{ratio}$ of at least three biological replicates (i.e. the seed stocks from different mother plants) is shown."

Fig. 5. Same as before. How do I see that there is a phenotype? There is no comparison with wild type. It is also unclear to which values the statistics refer to.

**Response:** To better enable the reader to see the phenotype(s), we now outline a representative cell in each figure panel with a white dashed line. The wild type is shown in the upper panel Fig. 5A. A direct comparison between the wild type and *trappii* mutants is now provided in an ANOVA analysis in Fig. S11 (new). The statistics in Fig. 5A,B refer to a pairwise comparison between light versus dark and dark versus darkW (water stress in the dark), as indicated by the bars above the graphs. The ANOVA analysis in Fig. S11 corroborates the pair-wise tests, reiterating the same conclusions.

Fig. 6: Please guide the reader through the figure and experiment.

**Response:** In a survey of root tip growth responses to our single versus additive stress conditions, we found that the dynamic changes in cell length across the root meristem were a very good representation of the different growth strategies of the wild type under different environmental conditions. Under additive stress, cells appear to exit the meristem very early on in the wild type (see green arrows pointing to steep slopes in Fig. 5C, previously Fig. 6A-D). In *trappii,* however, this was not observed and all growth curves were similar (see magenta arrows pointing to flat slopes in Fig. 5D, previously Fig. 6E-H). This is now better described in the text, as follows:

Results p. 10

"We have recently shown that root growth in response to water stress in the dark is due to a combination of cell division and rapid exit from the meristem (Kalbfuß *et al*, 2022). An early exit from the meristem can be visualized as cell elongation in cells close to the quiescent center. This was observed under dark and darkW conditions in the wild type (green arrows in Fig. 5C) but not in the *trappii* mutant *trs120-4* (magenta arrows in Fig. 5D). While the curves differed under the different environmental conditions in the wild type (Fig. 5C), these were fairly similar regardless of the environmental cue in *trs120-4* (magenta arrows in Fig. 5D; note that the grey shading, which designates the 95% confidence interval, overlaps)."

Fig. 8: I miss the connection with other shaggy-like kinases. This summary could be more complete. What about phosphorylation sites?

**Response:** We have added the other shaggy-like kinases as well as the three phosphorylation sites. To complete the summary, we now represent a surface view of the predicted TRAPPII structure in 3D (please see the cross-kingdom structural alignment in Fig. S4, new).

o Line 133-134: "we focus on the TRAPPII complex as a starting point as it is required for all aspects of TGN function, including the sorting of proteins such as PINs to distinct membrane domains" I did not find an obvious connection to the PIN transporters as well as clear data to TGN functions. This sentence was for me misleading about the context of this manuscript.

**Response:** We have now extended the analysis to include new conditional alleles of *trappii* as well as a quantification of root gravitropism. We write:

Discussion p. 13

"Furthermore, *bin2* higher order and *trappii* conditional mutants exhibited a synergistic genetic interaction with respect to root gravitropism.

Like other tropisms, root gravitropism requires organ bending as a result, at least in part, of the differential sorting of PIN transporters (Friml *et al*, 2002; Ding *et al*, 2011; Konstantinova *et al*, 2021). PIN2 polarity has been shown to be abolished in *trappii* mutants (Qi *et al*, 2011; Rybak *et al*, 2014) and to require TGN function (Naramoto *et al*, 2014; Qi *et al*, 2011; Rybak *et al*, 2014; Ravikumar *et al*, 2018)."

o Figure 1C: A supporting Western Blot control is needed, to fully validate the missing interaction of BIN2 with the truncated variants of TRS120 and CLUB. Additionally, swapping the constructs from DB to AD and vice versa will provide a better set-up of the interaction screen. This should be easily done in a few weeks.

**Response:** For an explanation of quality control, expression of the constructs, and how we avoid false positives in the Y2H screen, please see our response to Fig. 2 above.

TRS120-DB auto activates, which is why we cannot do the reciprocal orientation for TRS120, which would indeed be important (see Table R2 above).

o Line 431-432: "This presents intriguing implications regarding the potential role of the AtSK-TRAPPII module in meeting the unique demands of endomembrane traffic in plants." Why do the authors come to this assumption? Further discussion is needed here.

**Response:** We have elaborated on this point as follows:
Discussion p. 15
"Plant and animal cells differ in numerous ways, many of which can be attributed to the presence of the plant cell wall. Novel or expanded families of vesicle-trafficking genes are implicated in cell wall assembly (Assaad, 2001). Cell wall deposition and remodeling underly any growth response in plants. Growth in the face of challenging, restrictive environments is a distinctive survival strategy unique to plants (Assaad, 2001). In this study, we look at the tight regulation of differential growth that enables plants to thrive under multiple stress conditions, in the absence of a carbon or energy source."

o Figure 2F: What serves as positive controls? What is the purpose of showing every panel between each TRS120-T2 variant with CLUB-C2, CLUB-C3, TRS120-T1 and TRS120-T3 and not only interactions between BIN2 and the TRS120-T2 variants? Why are there six negative controls as it is every time the same control?
**Response:** The positive control is the WT TRS120-T2 truncation (first row), which interacts with the TRAPPII-specific subunits CLUB and TRS120 as well as with BIN2. The Y2H results of the phosphovariants with all the TRAPPII truncations were shown to show that our phosphovariants (i) exhibit the same interactions with CLUB and TRS120 truncations as the wild-type TRS120 construct and (ii) that they are expressed. The negative controls are now represented 3-4 times, one for each of the four experiments, or three of the four replicate experiments in the case of Fig. 2B. This is specified in the figure legend (see p. 29-31).

Clearly, title, abstract and statements have to be formulated differently. The discussion should contain a limitations paragraph in which the authors detail that conclusions are based on in vitro, yeast and plant IP-MS screening data, and they should describe approaches how the study can be continued in the future. Which alternative explanations are possible. Are SKs and TRAPP expressed and present in the same locations?
**Response:** We would please like to refer the reviewer to Table R1 for a list of the extensive *in vivo* evidence. We specify in the results whether the line of evidence is *in vivo* or *in vitro*. For localization, we have studied the BIN2:YFP construct from Kim *et al* (2009) and TRS120:GFP and find that they overlap in the cytosol (Fig. R1).

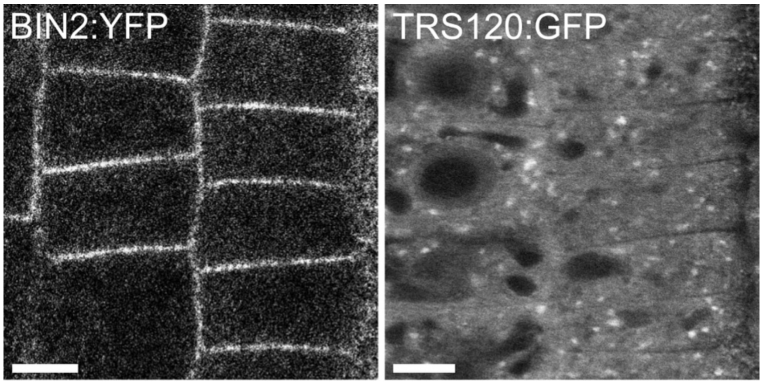


**Fig. R1. BIN2:YFP and TRS120:GFP may overlap in the cytosol.** BIN2:YFP localizes to the plasma membrane and cytosol, and is also seen in the nucleus (Vert & Chory, 2006). TRS120:GFP localizes to the TGN and to the cytosol (Qi *et al*, 2011; Rybak *et al*, 2014; Ravikumar *et al*, 2018; Naramoto *et al*, 2014). Scale bars represent 5 µm.

- Demonstrating interactions and phosphorylation by other approaches in vivo
- demonstrating effects of TRAPP phosphomutants and lack of kinases in vivo

**Response:** The phenotypes of phosphomutants *in planta* are presented in Fig. 6 (previously Fig. 7); kinase knock-outs *in vivo* are addressed in Fig. 7 (new), where we present genetic interaction data (see also Table R1).

- o Figure S9: Is it not a loss of chlorophyll instead of GFP? Does not look like a fluorescent image.

**Response:** We thank the reviewer for pointing this out. We now specify in the legend of Fig. S12 (previously Fig. S9) that these are not fluorescent images, just images taken with a binocular microscope of seedlings with white sectors due to loss of chlorophyll, likely due to silencing.

o Lacking information of pH of in vitro kinase assay solution with Mass-spectrometry.

**Response:** We thank the reviewer for pointing this omission out. The pH value is now specified in the methods section (see p. 20).

o What is the purpose of transferring 10 days old seedlings to fresh plates for scanning? Needs additional information for understanding, at the moment it sounds more like unnecessary extra stress for the seedlings.

**Response:** The seedlings often grow in ways that are difficult to trace for measuring organ lengths. Transferring them enables us to make sure that we can follow the root or shoot.

o Why are seedlings grown under constant light?

**Response:** These happen to be the settings for a shared tissue-culture room, they work and we did not wish to change them mid-experiment.

o Figure 5: It will be good to use ANOVA for statistics here. I personally doubt the high significance of some parameters, e.g. for club-2 cell width and cell surface area between dark and darkW due to the high standard errors. Rechecking with the original values is necessary. Why is there no comparison between wild-type and the two mutants?
**Response:** We now add an ANOVA test in Fig. S11B-D. Please note that in Fig. 5 the scales of WT vs. *trappii* mutants are different and, therefore, we are comparing the adaptation between different conditions (up vs. down between two conditions).

o Figure 7A - C, statistic is probably not correct. For example: in A statistical differences with $P<0.001$ between wild-type (~100 %) and TRS120SαβγD (~80 %), in C statistical difference of only $P<0.05$ between wild-type type (90 %) and TRS120SαβγD (60 %)
**Response:** We would please like to draw the reviewer's attention to the different error bar sizes. For 0mM mannitol we have error bars of only ±0.6% (Col) and ±0.9% (TRS120SαβγD) whereas ±2.8% (Col) and ±9.3% (TRS120SαβγD) for 400mM mannitol. Bigger error bars means bigger variation, leading to higher *P*-values.

o No information on IP-MS replicate numbers mentioned.
**Response:** We now specify that three replicate IP-MS experiments were performed.

o Also see comments above to figures
Minor comments:
o Figure 4, S7, S8, S11 and S12: It will be helpful to support the data with images of the seedlings.
**Response:** To help visualize important points, we now include seedling images in Fig. 4, Fig. 7, Fig. 8 and Fig. S10 (new Figure numbers).

o The introduction is quite lengthy with unnecessary information, e.g. about PIN transporters, but useful information about shaggy-like kinases and connection to brassinosteroid signaling is lacking.
**Response:** We have deleted unnecessary detail in the introduction and added background on the decision screen and on TRAPPII instead. AtSKs/BIN2 arise from interaction screens in the results; to avoid anticipating our results, they are introduced later. PINs are now integrated throughout the text in the context of gravitropism.

o Figure 1C: In the figure legend is no explanation of abbreviation "Co"; no explanation of BET3, TRS31, Tca17 and TRIPP; no indication that spots come from different plates (just visible by different brightness of the squares). Why are there eleven negative controls as it is every time the same control?
**Response:** We thank the reviewer for pointing this out and have corrected the figure to have three or four of the controls (one for each of four replicates) side by side, under the full title "control" rather than "Co.". The core subunits are now labelled and listed in the figure panel (Fig. 2D) and legend (see p. 30).

o Figure 1C is specifically for BIN2, but BIN2 was not identified in the IP-MS screen represented in Figure 1B. Why does 1C not focus on SK11/12/32, identified in 1B?

**Response:** AtSK11 was identified in CLUB:GFP IP-MS (Fig. 1B, 1C), and razor peptides common to BIN2 and its homologues were identified in TRS120:GFP IP-MS, meaning a different bait (Fig. S3D). Kinase - substrate interactions are transient and conditional. They are therefore difficult to detect. In addition, BIN2 belongs to a family with 10 members, so unique peptides are rare. The issue is likely one of both sensitivity and selectivity: we do not detect BIN2 specifically in IP-MS, but only in the context of a group of homologous proteins. However, and as stated in the results and Fig. 2 legend, the BIN2-TRS120 interaction was the only binary kinase-TRAPPII interaction we identified in Y2H, which is why this is shown in Fig. 2B. We list the different lines of evidence for the different AtSKs in Table R3:

**Table R3: AtSK-TRAPPII interactions.** * refers to razor (as opposed to unique) peptides.

| clade | Kinase | interacts with | Evidence | Figure |
|---|---|---|---|---|
| I | AtSK11 | CLUB:GFP | IP-MS | Fig. 1B |
| | | TRS120:GFP | IP-MS* | Fig. S3B-D |
| | | TRS120-T2 | in vitro kinase assay | Fig. 3B; Fig. S5A |
| | AtSK12 | CLUB:GFP | IP-MS | Fig. 1B |
| | | TRS120:GFP | IP-MS* | Fig. S3B-D |
| | AtSK13 | TRS120:GFP | IP-MS* | Fig. S3B-D |
| II | AtSK21/BIN2 | TRS120:GFP | IP-MS* | Fig. S3D |
| | | TRS120-T2 | Y2H | Fig. 2B, 2F |
| | | TRS120-T2 | in vitro kinase assay | Fig. 3A; Fig. S5B |
| | AtSK22/BIL1 | TRS120:GFP | IP-MS* | Fig. S3D |
| | AtSK23/BIL2 | TRS120:GFP | IP-MS* | Fig. S3D |
| | | TRS120-T2 | in vitro kinase assay | Fig. 3B |
| III | AtSK31 | TRS120:GFP | IP-MS* | Fig. S3C |
| | AtSK32 | CLUB:GFP | IP-MS | Fig. 1B |
| | | TRS120:GFP | IP-MS* | Fig. S3C |
| | | TRS120-T2 | in vitro kinase assay | Fig. 3B |
| IV | AtSK41 | TRS120:GFP | IP-MS* | Fig. S3B |
| | AtSK42 | TRS120:GFP | IP-MS* | Fig. S3B |

o Figure 1C shows several truncated variants of TRS120 and CLUB, a schematic overview as represented in Figure 2A will be helpful for the understanding of 1C. Order of variants should be the same (now: in 1C first TRS than CLUB in 2A first CLUB than TRS).

o Figure 1C: Interaction of TRS120 full-length with BIN2 is missing in this figure but is presented in Figure 2F.

**Response:** Full length TRS120:GFP constructs did not work in Y2H in our hands. In both figure panels (Fig. 2B, 2F new numbers), interactions with the TRS120-T2 truncation are shown. We have corrected the panels to fit the same sequence of truncations, as is represented in Fig. 2A.

o Result of Figure 2F is described after Figure 3. Better arrangement of Figures or text is needed here.
**Response:** We thank the reviewer for pointing this out and have restructured the text accordingly (Fig. 2 mentioned before Fig. 3).

o Figure 3A: Why was AtSK11 and not BIN2 used for the main figure? Better change Figure 3A with Figure S4 to keep the focus on BIN2. No explanation of the result in the text.
**Response:** We thank the reviewer for pointing this out and have revised the figures accordingly, with BIN2 *in lieu* of AtSK11 in the main Fig. 3A. We have also added a conclusion sentence to the results and legends, as follows:

Results p. 7
"This showed that BIN2 and AtSK11 phosphorylated AtTRS120-T2 *in vitro,* with a preference for wild-type sequences over non-phosphorylatable substrates such as AtTRS120-SαβγA (Fig. 3A; Fig. S5A)."

o Figure 3A: In the figure legend is no explanation of abbreviation CBB. What are the non-phosporylated variants? Where are they shown? Description sounds that only TRS120-T2-SαβγA versus TRS120-T2 WT was tested by t-test is this correct? And if yes, why?
**Response:** CBB is now spelt out in the legend. We now specify that non-phosphorylatable S to A TRS120-T2 variants were used as negative controls. All phosphovariants were compared to TRS120-WT but only SαβγA showed a significant difference in the AtSK11 kinase assay with radioactive substrates (Fig. S5A, new figure number, formerly Fig. 3A).

o No need for Figure 3B, information was already given in Figure 2A + B.
**Response:** Figure 3B is now merged with Fig. 3C.

o Figure 3C: Why BIL2 for Clade II and not BIN2?
**Response:** BIN2 kinase assays with mass spectrometry readout are now presented in Fig. S5B.

o Figure 4: Why are A-E not directly compared to wild-type but trs120-4 as seen in 4F? What is the purpose of using different types of diagram?
**Response:** We now have a full figure of violin plots (Fig. S8) to show (i) the distribution of data points and (ii) direct comparisons to the corresponding wild-type ecotypes (in grey) for all the lines.

o Figure 4H: Why are phyAphyBcry1cry2 and pyrpyl1pyl2pyl4 depicted? No description in the text.
**Response:** These are interesting as perception mutants, as compared to decision/integration or execution mutants. They are now referred to in the text as follows:
Results p. 9
"… the near-wild-type phenotype of higher order null mutants impaired in the perception of light or water stress (*phyAphyBcry1cry2* and *pyr1pyl1pyl2pyl4* in Fig. 4G-H)."

Discussion p. 14

"To address the first stage, perception, we looked at, for example, quadruple *phyAphyBcry1cry2* photoreceptor mutants; these failed to adjust their organ lengths to light versus dark conditions but had a highly significant response to additive stress (Fig. 4G-H; Fig. S8A; Fig. S9; Fig. S10G-I; Kalbfuß *et al*, 2022). The opposite was true for *bin2-3bil1bil2* and *trappii* mutants, which were able to respond to single stress factors but not to additive stress (Fig. 4; Fig. 5; Fig. S11; Kalbfuß *et al*, 2022)."

o Figure 6: Confusing order of given information in the figure legend. Sentence one belongs to D and H only, second sentence describes whole figure.

**Response:** The figure is now merged with Fig. 5 and the legend streamlined (see p. 32-33).

o Figure 6D + H, color difference between black and blue is hard to see, better change one into e.g. red.

**Response:** We thank the reviewer for pointing this out and have adjusted the shade of blue, which we associate with water stress, for optimal contrast.

o Figure 7D - F wrong indication of D to F, named in the description as A) - C). Why is E different to D in F (D and F: 0-1 is attenuated, >1 enhanced; E the other way around).

**Response:** We thank the reviewer for pointing out the wrong panel designation in the figure legend, which we have corrected. Root and hypocotyl responses to water deficit in the dark (abbreviated as darkW) were computed as

$$\frac{\text{organ length in the dark}}{\text{organ length under darkW}}$$

Because the hypocotyl and root have opposite responses to our additive stress conditions (hypocotyl length decreases but root length increases in response to water stress in the dark, as shown in Fig. 4) the RQs move in opposite directions. This is now explained in the methods (see page 21) and figure legends. Additionally, we have added Supplementary Table 3 (new), which describes the thresholds for attenuated and enhanced response quotients for organ and ratio adjustments.

o Figure S9A: Indication of protein size on the Coomassie gel is missing and the respective position of 160 kDa is not visible on the gel.

**Response:** Protein sizes are given as kDa on the left in the figure. Because no protein band was visible for TRS120:GFP on the Coomassie stain (see Fig. R2; arrow points to approximately 157kDa, where the TRS120:GFP band would be expected) we used rubisco as a loading control.

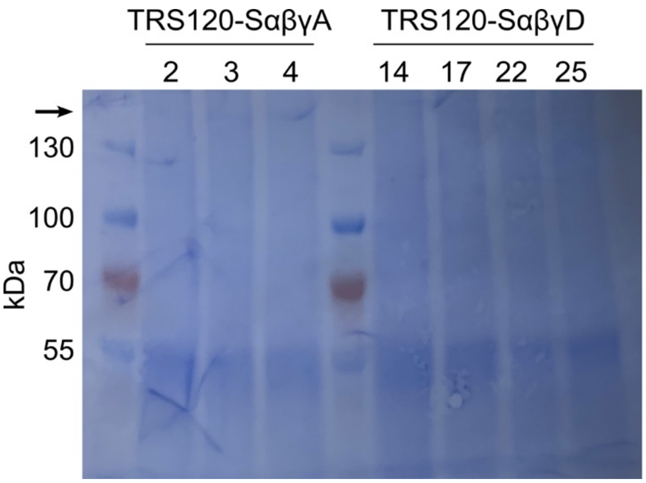


**Fig. R2. Full Coomassie-stained membrane.**

o Figure S12D: No explanation of the color code in the figure legend.

**Response:** The color code, which is consistent throughout the manuscript, is now mentioned in the legend of Fig. S15D (previously Fig. S12D). Briefly, the hypocotyl is depicted in green, the root in beige-brown and the hypocotyl/root ratio in blue shades. For the different conditions (light; dark; dark with water stress (darkW)), we use an increasing opacity gradient of 50% for dark and 100% for darkW conditions.

o Consistent labelling and layout of all Figures and Supplemental Figures will be helpful. E.g., Figure 3A and S4; in S8A-E + S11A-C bars of different conditions have the same color.

**Response:** The represented autoradiographs (Fig. 3A; Fig. S5A (now)) are now consistently labeled. The hypocotyl and root bars are colored according to the color code in the manuscript mentioned above (please see response to previous comment).

Most of the figure legends are quite shortly described and lack information about what kind of data is presented.

**Response:** We have thoroughly revised the figure legends to be more complete and explanatory.

o YFP parameters are described in material and methods, but no YFP construct appeared in the manuscript to my knowledge.

**Response:** We thank the reviewer for pointing this out. YFP was only used for the rebuttal (Fig. R1) and has therefore been removed from the methods section.

• Lines 243-245: Text is nearly identical to Kalbfuß et al., 2022.
• Lines 246-254: Text is identical to Kalbfuß et al., 2022.

**Response:** We have edited the text further to minimise overlap.

Please see the above and below comments to figures and figure legends.

Overall, the manuscript may have very interesting data and new findings. It is very interesting that the authors study the regulation of a protein complex that may mediate environment responses and intracellular Golgi functions. However, it is very difficult to follow and understand the ideas and concept of the manuscript. This manuscript is based on a previously published interactome study by Rybek et al. 2014, Steiner et al. 2016, Kalde et al. 209. Moreover, a physiological approach is published in Kalbfuß et al. 2022. The outcomes and conclusions from these previously published manuscripts and the emanating open questions addressed here should be clearly described in the introduction. This is currently not the case. Moreover, many experimental approaches and results (e.g. figures, figure legends) are not properly described. Overall, it is therefore not possible to understand the manuscript without studying in depth all other manuscripts. Before the manuscript can be more thoroughly judged, it is necessary that the authors rewrite the manuscript, reorganize it and explain better their ideas and approaches. It is also necessary to explain and define unusual terms such as "decision mutants", "limited budget" and "conflict of interest" experiments, which are crucial for the understanding. The importance of the TRAPPII complex should be illustrated using specific physiological examples and the context in which this complex is studied here has to be explained. Before this is not corrected, the following assessment will remain rather incomplete.
Another complication is that two subunits of TRAPP were studied and different types of SKs, however, authors did not systematically analyze all interactions. At least it should be thoroughly described, and a flow chart would be helpful as supplemental figure clearly describe which types of proteins were tested in the different assays.
**Response:** We kindly refer the reviewer to Table R3 for a list of different AtSK-TRAPPII interactions identified in the study.

The introduction is not well written. It is very lengthy, however the important messages from previous publications are left out. Thus the open question is not understandable (see above).
**Response:** We have streamlined the introduction and now explicitly define novel terms such as "decision mutants", "limited budget" and "conflict of interest". We also pose the open questions more clearly; these include our incomplete understanding of

- how environmental stimuli are translated into sorting decisions
- the role of the plant TGN and of AtTRAPPII in stress responses and in decision-making processes
- the physiological role of the TRAPPII complex across kingdoms

Instead, the results parts start with introduction again. Explanations are also lacking in every result paragraph on the approach and expected data.
**Response:** We now provide more information on the approach in each section, while avoiding an anticipation of the results.

The Discussion is also not very well written. It is much focused on physiological and molecular actions and consequences in plants. However, there should be at first a technical discussion on the relevance since in the study is based on in vitro and heterologous expression data, and the physiological analysis was only conducted with knockouts but not phosphomutants.

Therefore, the link between the protein interaction and physiological functions needs to be worked out.

**Response:** The phenotypic analysis of phosphomutants is presented in Fig. 6 (previously Fig. 7), where we show opposite impacts of the non-phosphorylatable and the phosphomimetic variants in terms of seed germination under osmotic stress as well as responses to additive stress *in planta*. In addition, the new high order mutant analysis of *bin2-3bil1bil2 trs120i* (Fig. 7, new) suggests an additional physiological implication of the BIN2/AtSK-TRAPPII interaction in root gravitropism (see also Table R1).

**Referees cross-commenting**

My colleague and I have read thoroughly the manuscript and found a number of issues which we indicated in our review. These points can be fixed by the authors, if they formulate more carefully and remove the overstatements. They should also work on reorganizing and including more explanations.

Reviewer #3 (Significance (Required)):

The following aspects are important:

One new aspect of this story is the validation of interaction of TRAPII subunits as substrate for AtSKs and their action as phosphorylation agents shown in vitro. The other new aspect is the phenotypical characterization of trapii mutants under stress-conditions (grown in darkness) and additive stress (with additional drought stress). The potential interaction with brassinosteroid signaling via BIN2 is intriguing.

A strength is that a new interaction is further studied. A weakness is that the studies are primarily conducted in yeast and in vitro, leaving open how relevant this process is in plants. A strength is further studies and phenotypic analysis of trapii mutant effects. A weakness is that this mutant analysis is disconnected from the action of SKs.

**Response:** Higher order *bin2* null and *trappii* null or inducible alleles have a number of related phenotypes, including

- aberrant organ and cellular hypocotyl and root responses in the allocation decision screen (Fig. 4, Fig. 5 this study cf. Kalbfuß *et al*, 2022)
- root gravitropism (Fig. 7)

Please see Table R1 for further *in vivo* evidence.

Further, the writing should be improved and more clear (see comments above). The introduction gives the impression of a stronger investigation of TGN function, which is from my point of view not the case and should be reformulated and/or put into a deeper context with known literature.

**Response:** We address TGN function with some of the most adequate mutants available to us: *echidna* and *trappii*.

The authors switch several times between the different TRAPPII subunits and shaggy-like kinases in the main figures which made it for me very confusing. I believe that rearranging some data/figures will improve the understanding of the story. The text is also lacking explanations of many abbreviations and gene names which caused more difficulties in understanding the story and slowed down the reviewing process. From my point of view it seems to be necessary to read the often cited Kalbfuß et al., 2022 publication before, as many important technical aspects and scientific background, e.g. the reason to use specific control mutants, are well explained there, but are lacking in this manuscript and needs improvement.

Based on the cited literature in this manuscript the direction of the story with "limited budget" and "conflict of interest" situations to classify mutants methodically seems to be a recently emerged approach. Apart from that this manuscript provides only new impact on TRAPII and AtSKs specific knowledge based on well-established and frequently used techniques that address the problem in vitro and in a heterologous system. Therefore, this story will be interesting for researchers specialized in stress responses, TGN and growth defects as well as important for basic research. Limitations in interpretation are present.
**Response:** *In vivo* data (Table R1) include IP-MS, pharmacological inhibition, mutant analysis, genetic interactions and the analysis of phosphovariants *in planta*. These enable us to reach the conclusions that are carefully worded in the title and abstract.

Our field of research is related to nutrition-regulated processes especially in Arabidopsis with a strong methodological background in interactomics, physiological, morphological and molecular responses and biochemical approaches and microscopy.

**Response:** We thank Reviewer 3 for critically reading our manuscript and providing detailed and specific feedback, which we feel will considerably have improved the readability of the text.
