## Supplemental Information for "Regulation of adaptive growth decisions via phosphorylation of the TRAPPII complex in Arabidopsis"

**SUPPORTING INFORMATION**

**Figure S1.** GO term enrichment and selected brassinosteroid signaling components in IP-MS with the TRAPPII-specific CLUB:GFP subunit as bait.

**Figure S2.** Fragment ion mass spectra of AtTRS120 phosphopeptides found in AtTRS120:GFP IP-MS.

**Figure S3.** Fragment ion mass spectra of AtSK/GSK3 kinases found in AtTRS120:GFP IP-MS.

**Figure S4.** Cross-kingdom structural alignment of TRAPPII.

**Figure S5.** AtSK11 and BIN2 phosphorylate AtTRS120: *in vitro* kinase assays.

**Figure S6.** Pharmacological inhibition or enhancement of TRAPPII phosphorylation by AtSKs *in vivo*.

**Figure S7.** Cytokinesis and protein sorting in *bin2-1*.

**Figure S8.** Response to single versus additive stress: violin plots.

**Figure S9.** Response to additive stress: response quotients and volcano plots for the hypocotyl and root.

**Figure S10.** Light responses in *bin2-3bil1bil2*, *trappii* and *echidna* mutants.

**Figure S11.** Cellular hypocotyl parameters of *trappii* mutants under single and additive stress conditions.

**Figure S12.** Characterization of TRS120:GFP phosphovariants.

**Figure S13.** Localization of TRS120:GFP phosphovariants.

**Figure S14.** The etiolation response in TRS120 phosphovariants.

**Figure S15.** Role of TRS120 phosphorylation status on adaptive growth responses under additive stress conditions.

**Movie S1.** 3D projection of cross-kingdom structural alignment of TRAPPII.

**Table S1.** Mutant lines used in this study.

**Table S2.** Selected brassinosteroid signaling proteins in the CLUB interactome.

**Table S3.** Thresholds for attenuated, normal and enhanced response quotients.

**Supporting Methods.**

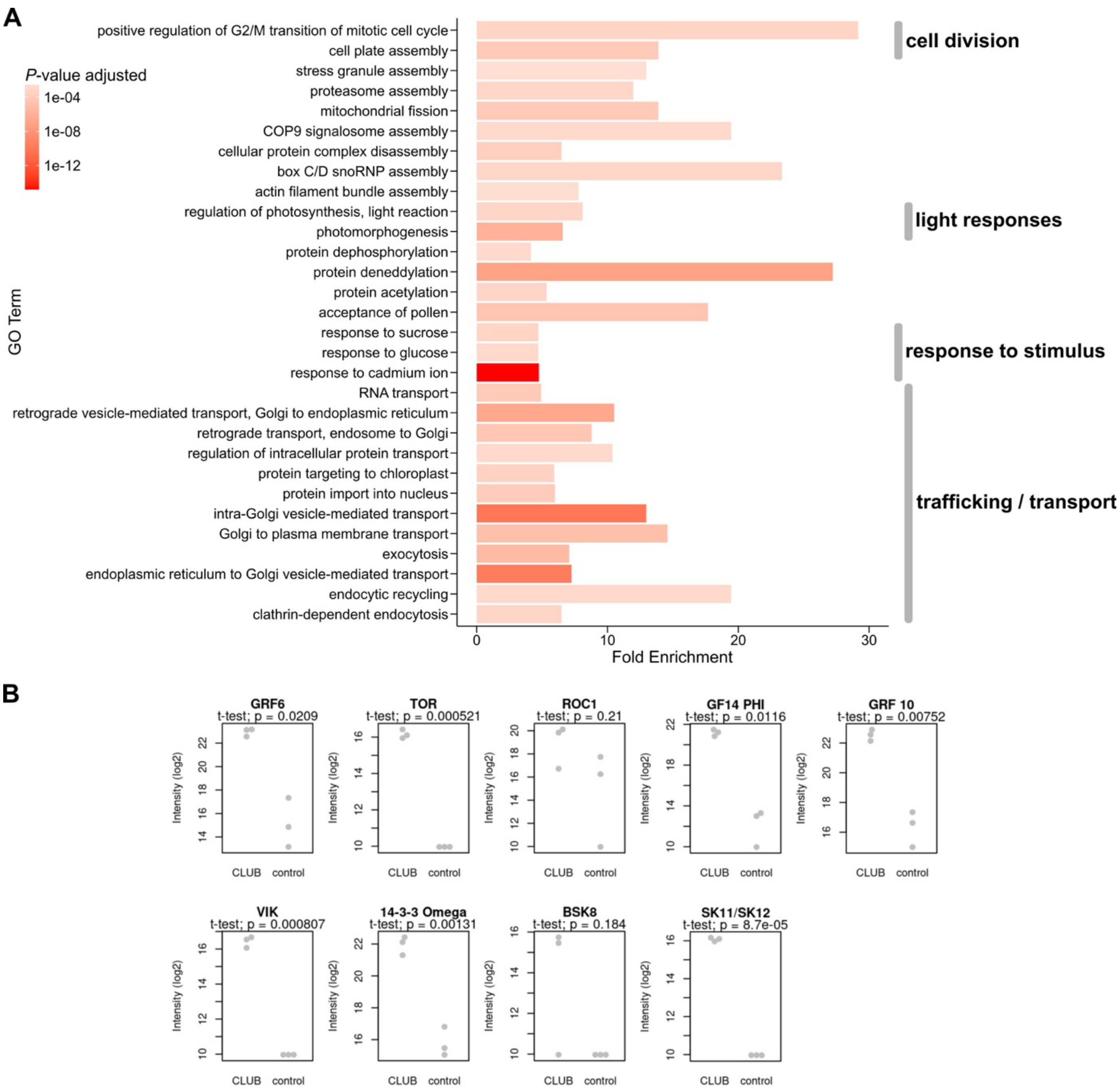


**Figure S1. GO term enrichment and selected brassinosteroid signaling components in IP-MS with the TRAPPII-specific CLUB:GFP subunit as bait.**
Light-grown inflorescences were used (see Fig. 1B-C; Kalde *et al*, 2019). As control the soluble GFP empty vector was used. Three replicates were carried out for the IP-MS experiments.
**A.** Gene ontology (GO) term enrichment analysis of the TRAPPII interactome. Depicted are highly abundant (fold enrichment ≥ 4) and significant (FDR-adjusted *P*-value ≤ 0.003) GO term associations of biological processes of level 0. The length of each bar corresponds to the fold enrichment of GO terms associated with detected proteins, while the color intensity indicates the significance given as the *P*-value adjusted for the false discovery rate (FDR). The GO term enrichment analysis was carried out with high confidence interactors (intensity ratio > 8 and *P*-value ≤ 0.02; see Fig. 1B). Note that the majority of GO terms are associated with trafficking and transport (12/30). GO terms associated with response to stimulus, light responses and cell division were also enriched.
**B.** Individual intensity based absolute quantification (iBAQ) values in log2 scale of brassinosteroid related proteins (highlighted in Fig. 1C) found in each CLUB:GFP IP-MS replicate. Selected brassinosteroid-related proteins were differentially enriched over light- versus dark-grown seedlings in a different IP-MS experiment. Dots clustered at the bottom of the graphs represent peptides that were not detected (or detected as very low intensity) in the sample. *P*-values were obtained using the Welch's t-test. Related to Fig. 1, Table S2.

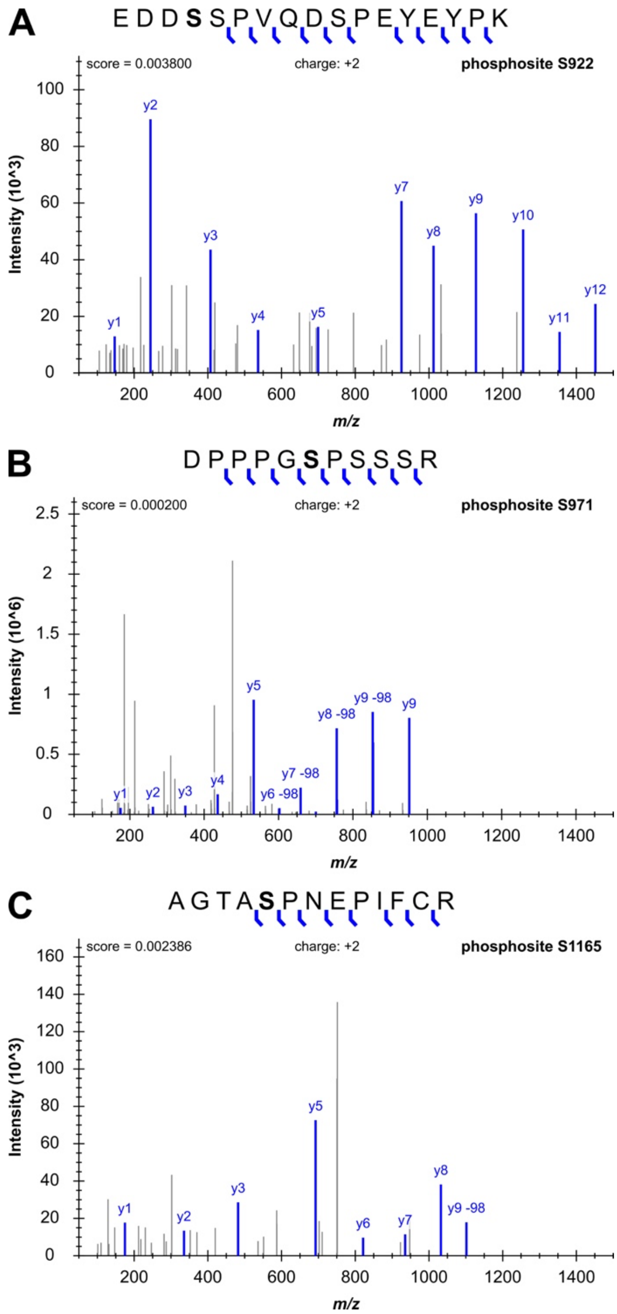


**Figure S2. Fragment ion mass spectra of AtTRS120 phosphopeptides found in AtTRS120:GFP IP-MS.**

Seedlings were grown in the light.

**A.** α-phosphosite of AtTRS120 at amino acid position S922. The spectrum with the phosphorylated S922 residue was identified with a 1% FDR, as can be seen in the deposited Skyline library. Note that the depicted y-ion series does not show the phosphorylation at S922.

**B.** β-phosphosite of AtTRS120 at amino acid position S971.

**C.** γ-phosphosite of AtTRS120 at amino acid position S1165.

The y-ion series is highlighted in blue and non-annotated fragment ions in gray. Phosphorylated serines are written in bold letters. These sites are annotated as GSK3 sites in the PPSP (https://www.phosphosite.org) database. Note that not all phosphosites we detected *in vivo* (highlighted in yellow in Fig. 2C) are depicted here. Spectra are taken from the deposited Skyline library, resulting from a MaxQuant search. At least three replicates were carried out for the IP-MS experiments. Related to Fig. 2.

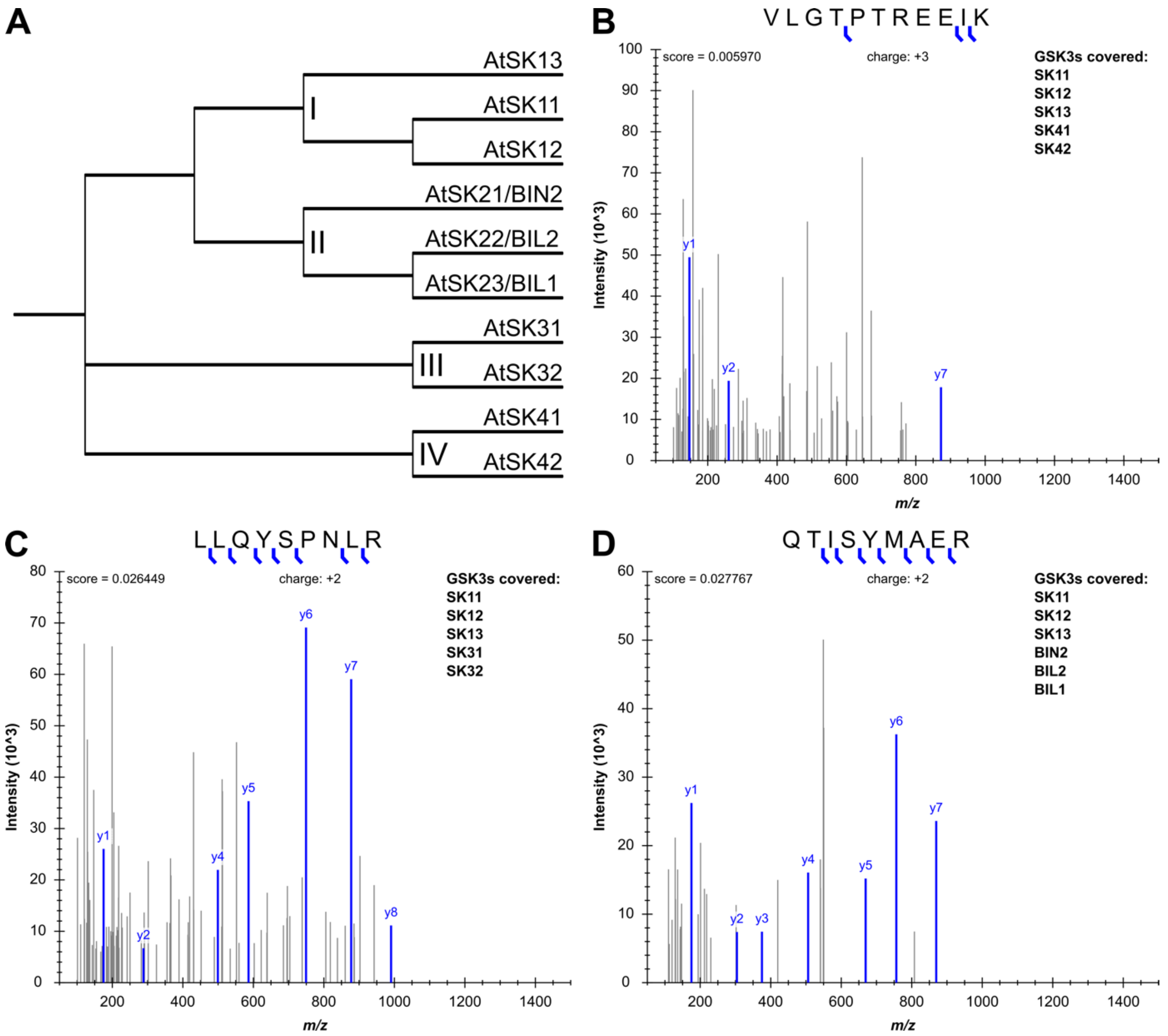


**Figure S3. Fragment ion mass spectra of AtSK/GSK3 kinases found in AtTRS120:GFP IP-MS.** The AtSK/GSK3 kinase family in Arabidopsis consists of 10 isoforms. Therefore, mass-spectrometric evidence of different peptides can account for the presence of different kinases in a co-immunoprecipitation with light-grown TRS120:GFP seedlings as bait. The y-ion series is highlighted in blue, non-annotated fragment ions in gray.

**A.** The Arabidopsis genome encodes ten shaggy-like kinases, which are classified into four clades. Multiple sequence alignment in UniProt using full-length protein sequences.

**B.** VLGTPTREEIK is shared by AtSKs in clades I and IV.

**C.** LLQYSPNLR is shared by AtSKs in clades I and III.

**D.** QTISYMAER is shared by AtSKs in clades I and II.

Spectra are taken from the deposited Skyline library, resulting from a MaxQuant search. At least three replicates were carried out for the IP-MS experiments. Related to Fig. 2.

A

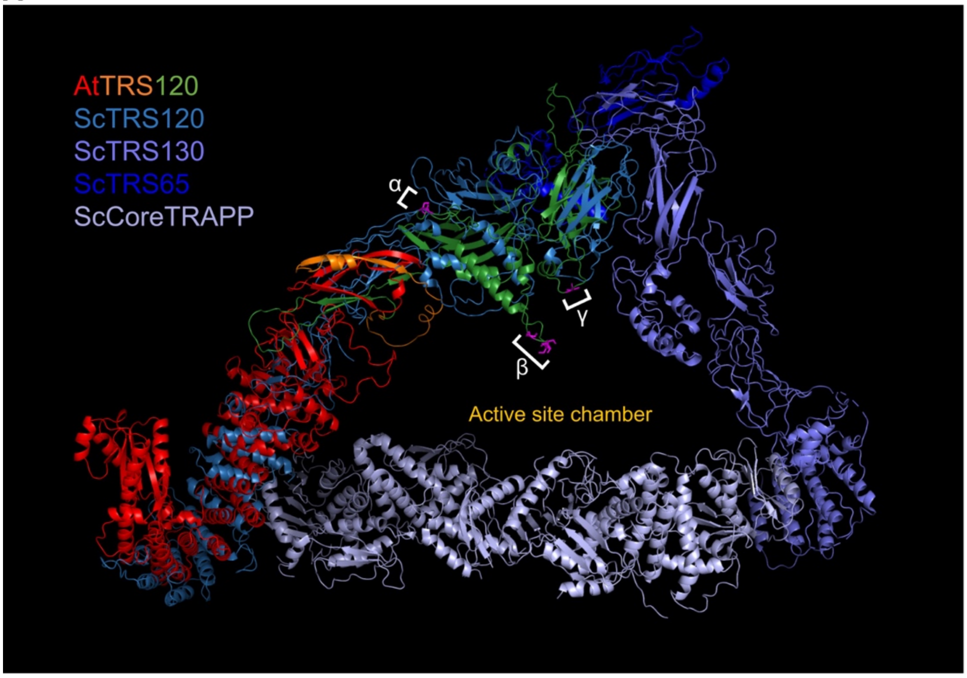


B

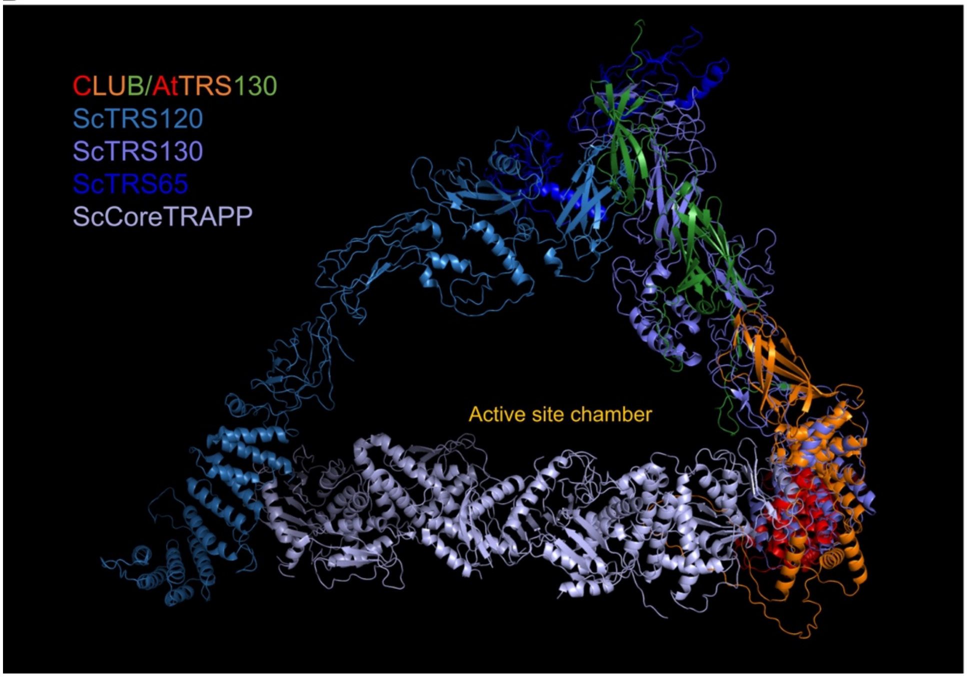


**Figure S4. Cross-kingdom structural alignment of TRAPPII.**

AlphaFold predictions (Jumper *et al*, 2021; Varadi *et al*, 2022) of *Arabidopsis thaliana* (At) **A.** AtTRS120 and **B.** CLUB/AtTRS130 aligned with the open formation of the TRAPPII monomer structure without substrate in *Saccharomyces cerevisiae* (Sc) resolved with cryo-electron microscopy (*in vitro*) from Mi *et al* (2022). Alignments were performed with the align algorithm in PyMOL. Reported root-mean-square deviations (RMSD) were 16.966 Å (3775 atoms) for AtTRS120/TRAPPC9 and 12.953 Å (1660 atoms) for CLUB/AtTRS130/TRAPPC10. The ScTRAPPII monomer is depicted in different shades of blue (ScCoreTRAPP with ScTCA17: blue-grey; ScTRS130: purple; ScTRS120: light blue; ScTRS65: dark blue). AtTRS120 and CLUB/AtTRS130 are colored based on their sequence conservation (red: conserved sequences; orange: intermediate conservation; green: plant-specific sequences - as depicted in Fig. 2A, 2D). Note that the β and γ phosphorylation sites of AtTRS120 (magenta sticks) face inwards, toward the active site chamber (including the RAB11/Rab-A GTPase binding pocket) proposed by Mi *et al* (2022) and Bagde & Fromme (2022). Related to Fig. 2, Movie S1.

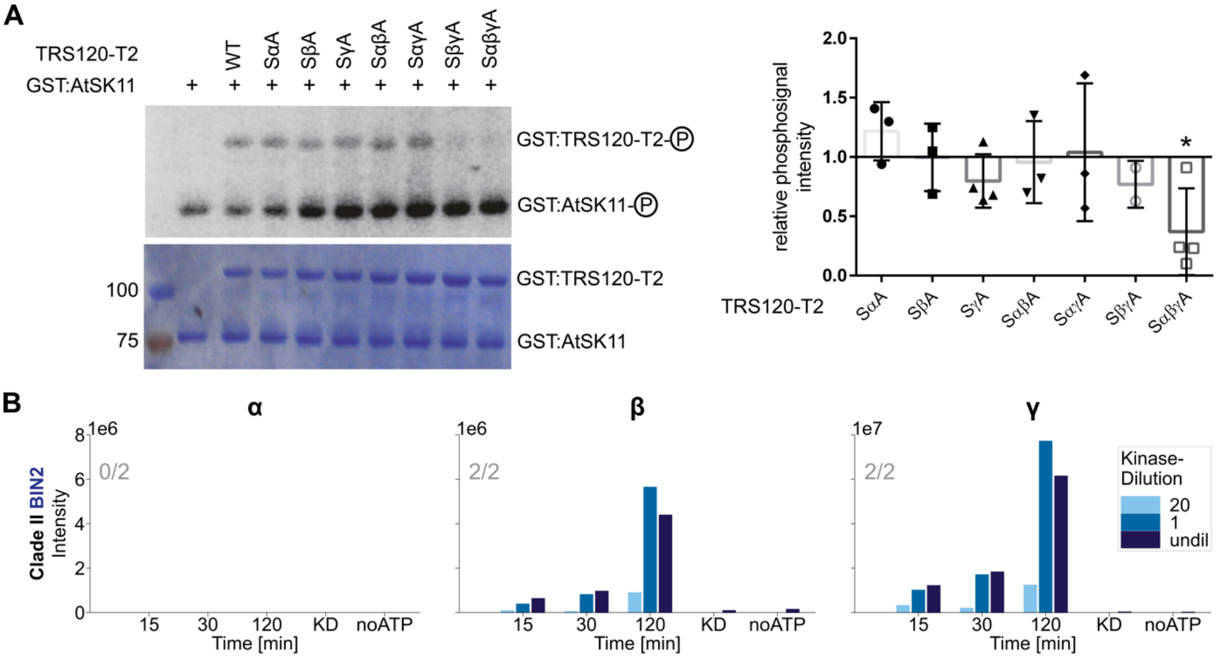


**Figure S5. AtSK11 and BIN2 phosphorylate AtTRS120: *in vitro* kinase assays.**

**A.** *In vitro* kinase assays using GST:AtSK11 (72 kDa) and GST:TRS120-T2 (100 kDa). The change of phosphosignal is shown in a representative autoradiograph (upper panel) and the loaded protein amount in the corresponding CBB (Coomassie stain, lower panel). Non-phosphorylatable S to A TRS120-T2 variants were used as negative controls. The means ± SD of phosphosignals were normalized to the protein amount and related to non-mutated TRS120-T2 wild-type control. Note that AtSK11 phosphorylated AtTRS120-T2 *in vitro*, with a preference for wild-type (WT) sequences over non-phosphorylatable AtTRS120-SαβγA. n = 3 independent experiments; *: $P<0.05$ for significant differences to TRS120-T2 WT (set at 1.0 right panel) determined by using a one sample two-tailed t-test.

**B.** *In vitro* kinase assay with mass-spectrometry readout using BIN2 as kinase and TRS120-T2 truncation as substrate. Dilution series of the kinase are depicted in different shades of blue. BIN2 phosphorylated the β and γ site of TRS120 with a preference for the γ site (1e7 for TRS120-γ versus 1e6 for TRS120-β on the Y axis). Samples incubated for 120 min in a kinase buffer without ATP, or samples in which the kinase was heat inactivated (KD), served as negative controls. The numbers in grey in each plot denote the number of times the phosphorylation event was seen in the given number of independent replicates. Related to Fig. 3.

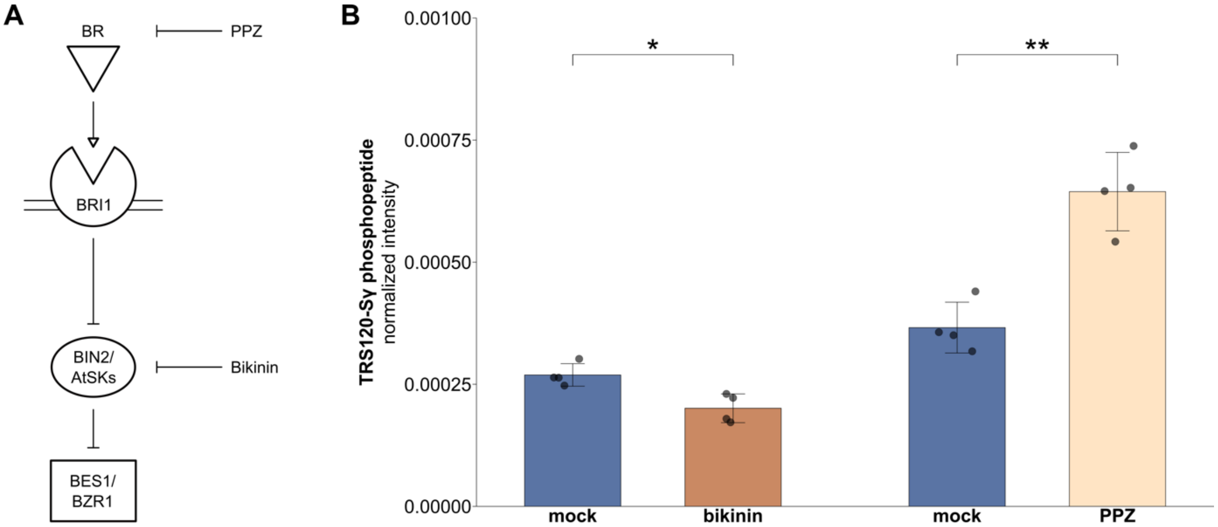


**Figure S6. Pharmacological inhibition or enhancement of TRAPPII phosphorylation by AtSKs *in vivo*.**

**A.** Bikinin is an inhibitor of shaggy-like kinases (AtSKs), whereas PPZ is a BR biosynthesis inhibitor that relieves BR-mediated BIN2 inhibition.

**B.** Impact of the pharmacological inhibitors bikinin and PPZ on the TRS120 phosphorylation status *in vivo*. IP-MS was carried out on light-grown TRS120:GFP seedlings treated with bikinin, PPZ, or the respective mock-controls. The phosphorylated peptides were further analyzed via Skyline (MacLean *et al*, 2010). Normalized intensities were calculated as the ratio of the TRS120-Sγ phosphopeptide intensity over the sum of all TRS120 peptide intensities found in the respective experiment. The extent of phosphorylation at the TRS120-γ site was significantly decreased by bikinin. In contrast, PPZ treatment increased the phosphorylation of the TRS120-Sγ peptide *in vivo*. *P*-values were computed with a two-tailed student's t-test (*: $P<0.05$; **: $P<0.01$). Mean ± SD of 4 replicates are shown for the control and treatment. Related to Fig. 2-3.

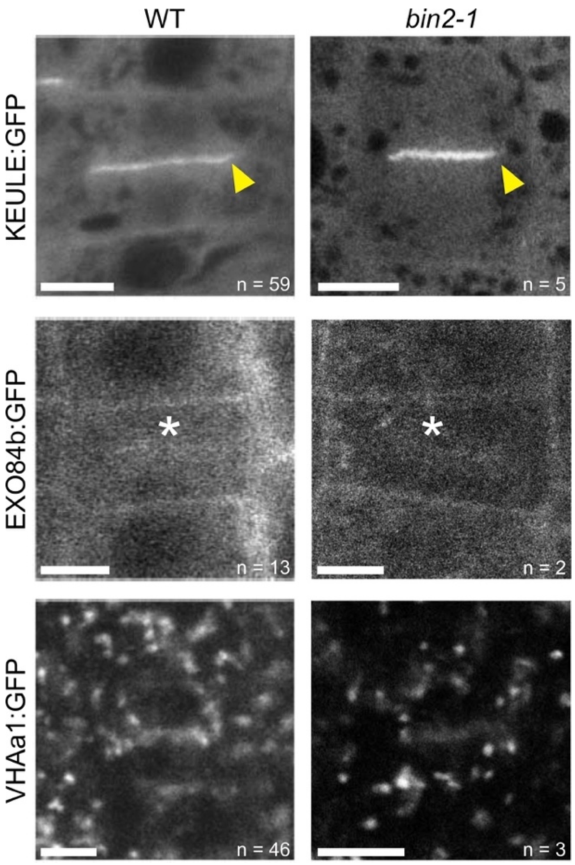


**Figure S7. Cytokinesis and protein sorting in *bin2-1*.**
Localization patterns of diverse markers in root tip cells of light-grown seedlings. Confocal Scanning Laser Microscopy. A protein that resides at the cell plate during cytokinesis in the wild type, $P_{KEU}$::KEULE:GFP (Steiner *et al*, 2016b), was also seen at the cell plate in *bin2-1* mutants (yellow arrowhead). Conversely, markers that are largely excluded from the cell plate (white star) in the wild type ($P_{EXO84b}$::EXO84b:GFP (Fendrych *et al*, 2010), $P_{VHAa1}$::VHAa1:GFP (Dettmer *et al*, 2006)) are also excluded in *bin2-1* mutants. Cell plate formation and morphology in *bin2-1* also did not visibly differ from the wild type. Sample numbers n = number of imaged cells. Scale bars represent 5 μm.

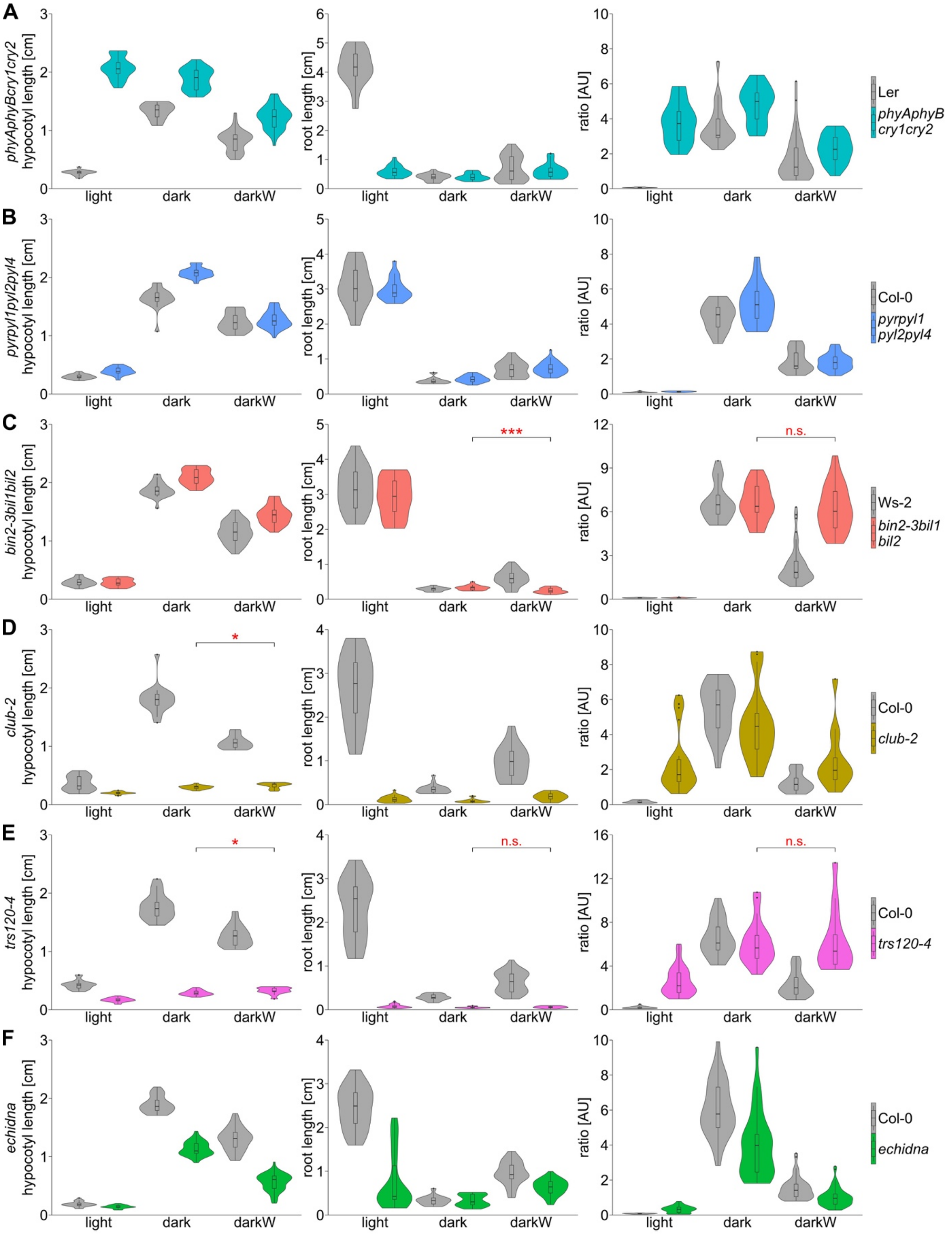


**Figure S8. Response to single versus additive stress: violin plots.**

Seedlings were germinated on ½ MS in the light (light), in the dark (dark) or in dark with -0.4 MPa water stress (darkW). Violin plots depict a direct comparison of organ lengths or the hypocotyl/root ratio between the wild type and **A.** *phyAphyBcry1cry2*; **B.** *pyrpyl1pyl2pyl4*; **C.** *bin2-3bil1bil2*; **D.** *club-2*; **E.** *trs120-4*; **F.** *echidna* in the same experiment and on the same PEG plates. The same datasets as shown in Fig. 4B-E and Fig. S10B-E were used. Decision phenotypes of *bin2-3bil1bil2* and *trs120-4* mutants are highlighted as follows: *bin2-3bil1bil2* showed an opposite root adaptation (red asterisk) and no hypocotyl/root adaptation (non-significant (n.s.) in red) to the multiple stress conditions of darkW. *trappii* null mutants, *club-2* and *trs120-4*, had an attenuated, but significant etiolation response but failed to correctly adjust their hypocotyl length under darkW (red asterisks). Additionally, *trs120-4* mutants showed no root and no hypocotyl/root ratio adaptation (n.s. in red) to darkW conditions. At least 3 experiments were performed for each line, and a representative one is shown here on the basis of RQ and *P*-values. *P*-values for decision phenotypes were computed with a two-tailed student's t-test (*: P<0.05; ***: P<0.001) Related to Fig. 4, S10.

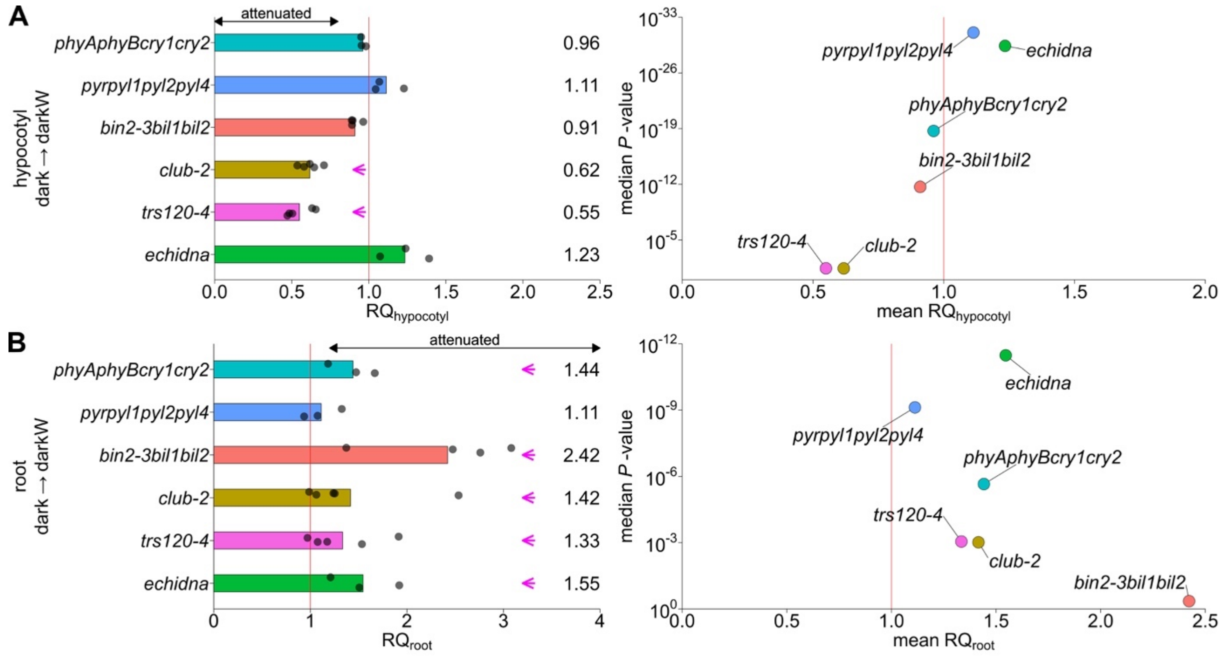


**Figure S9. Response to additive stress: response quotients and volcano plots for the hypocotyl and root.**

Response quotients (RQ, left) and volcano plots (right) of the hypocotyl and root adaptations to dark-to-dark with water stress (-0.4 MPa, darkW). RQ response quotients are normalized to the respective wild-type response; a value of 1 (vertical red line) indicates that the response to a shift from dark to darkW is identical to that of the respective wild-type ecotype. Each replicate is represented by a dot. Mean RQ values are given on the right. Volcano plots show the mean RQ depicted on the X axis and the median *P*-value of the response on the Y axis (negative log scale).

**A.** Hypocotyl responses to dark versus darkW conditions. Note that *club-2* and *trs120-4* had attenuated responses ($RQ_{hypocotyl}$ < 0.8, magenta arrows).

**B.** Root responses to dark versus darkW conditions. Note that the triple *bin2-3bil1bil2* knock out (from Kalbfuß *et al*, 2022) had the strongest attenuated $RQ_{root}$ phenotype (mean $RQ_{root}$ = 2.42, magenta arrow) with an insignificant root response. *phyAphyBcry1cry2*, *club-2*, *trs120-4* and *echidna* mutants show also an attenuated root response ($RQ_{root}$ > 1.2, magenta arrows). Quadruple *pyrpyl* mutants did not have a phenotype, possibly because only four genes are knocked out in a family with fourteen members (Ma *et al*, 2009; Park *et al*, 2009). Due to the opposite adaptations of the hypocotyl and root under the additive stress conditions (decrease in hypocotyl and increase in root length from dark-to-darkW), the respective thresholds for attenuated responses are opposite (Fig. S9A, compared to S9B). Related to Fig 4.

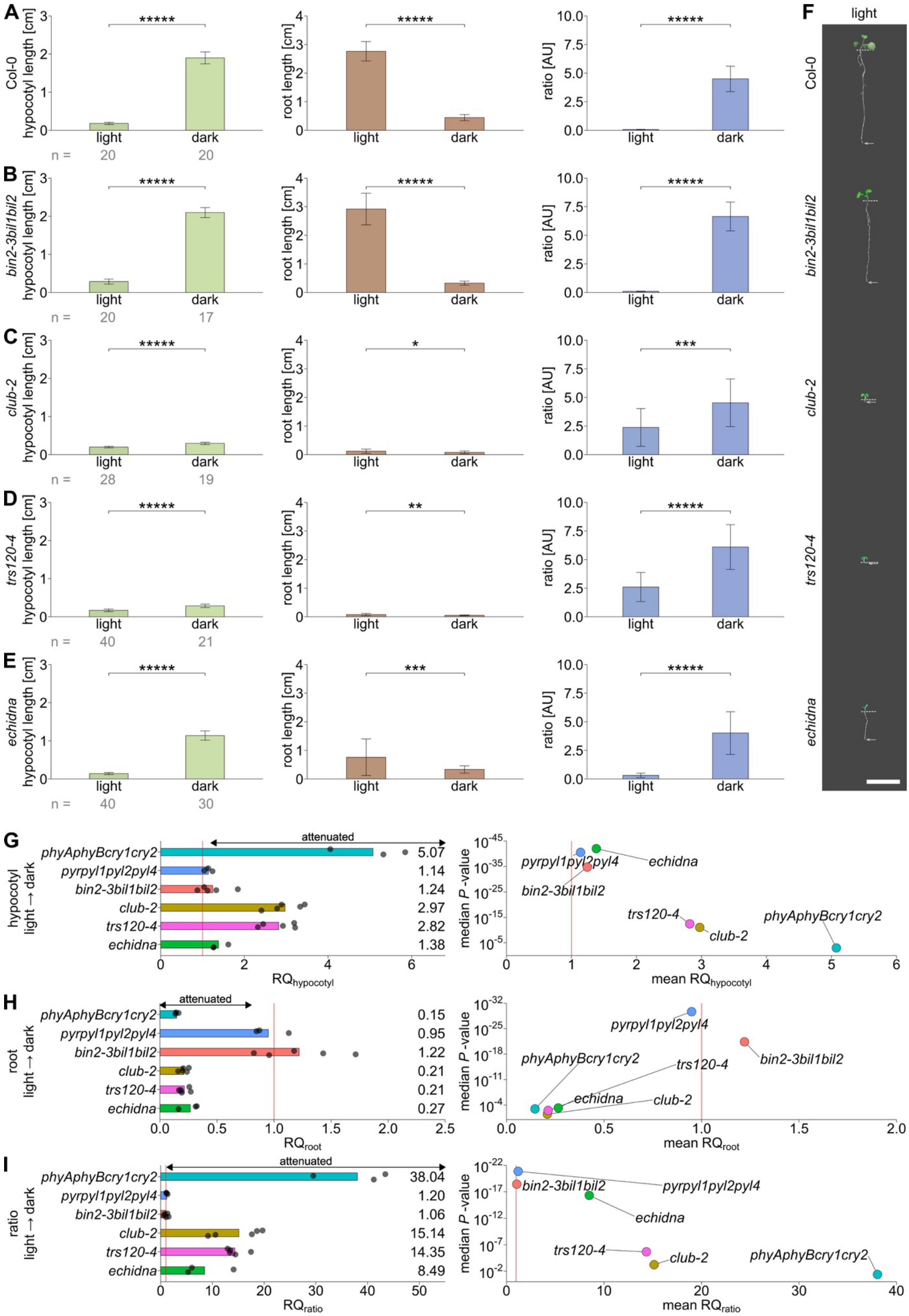


**Figure S10. Light responses in *bin2-3bil1bil2*, *trappii* and *echidna* mutants.**
Seedlings were germinated on ½ MS in the light or dark **A.** Col-0 (wild type). **B.** BR signaling mutant *bin2-3bil1bil2* triple knockout (from Kalbfuß *et al*, 2022). **C.** *club-2*, a null *trappii* allele. **D.** *trs120-4*, a null

*trappii* allele. **E.** *echidna* null allele, impaired in TGN structure and function. At least 3 experiments were performed for each line, and a representative one is shown here on the basis of RQ and *P*-values. **F.** Representative images of light-grown seedlings of Col-0 (wild type) and mutants shown in A-E. Dotted lines mark the hypocotyl-root junction, whereas arrows point to the end of the root. For corresponding dark-grown seedlings see Fig. 4F. Scale bar is 1 cm. **G-I.** Response quotients (RQ, left) and volcano plots (right) of G) the hypocotyl, H) the root and **I**) the hypocotyl/root ratio responses to light versus dark. RQs are normalized to the wild-type quotient; a value of 1 (vertical red line) indicates that the response to a shift from light to dark is identical to that of the respective wild-type ecotype. Each replicate is represented by a dot. Volcano plots show the mean RQ depicted on the X axis and the *P*-value of the response on the Y axis (negative log scale; a median of all replicates was used). *trappii* mutants *trs120-4* and *club-2*, the TGN mutant *echidna* and the higher order light perception null mutant *phyAphyBcry1cry2* show a severely attenuated etiolation response (mean $RQ_{hypocotyl}$ and mean $RQ_{ratio}$ $\gg 1.2$ as well as $RQ_{root} \ll 0.8$). Note that due to the opposite adaptations of the hypocotyl and root from light-to-dark conditions (increase in hypocotyl, decrease in root length and therefore increased hypocotyl/root ratio), the respective thresholds for attenuated responses are opposite (Fig. S10G, **I**, compared to S10H). The number (n) of seedlings measured per condition is in grey below the mean ± SD bar graphs. *P*-values were computed with a two-tailed student's t-test (*: P<0.05; **: P<0.01; ***: P<0.001; ****: P<0.0001; *****: P<0.00001). Ecotypes are described in Table S1. Related to Fig. 4.

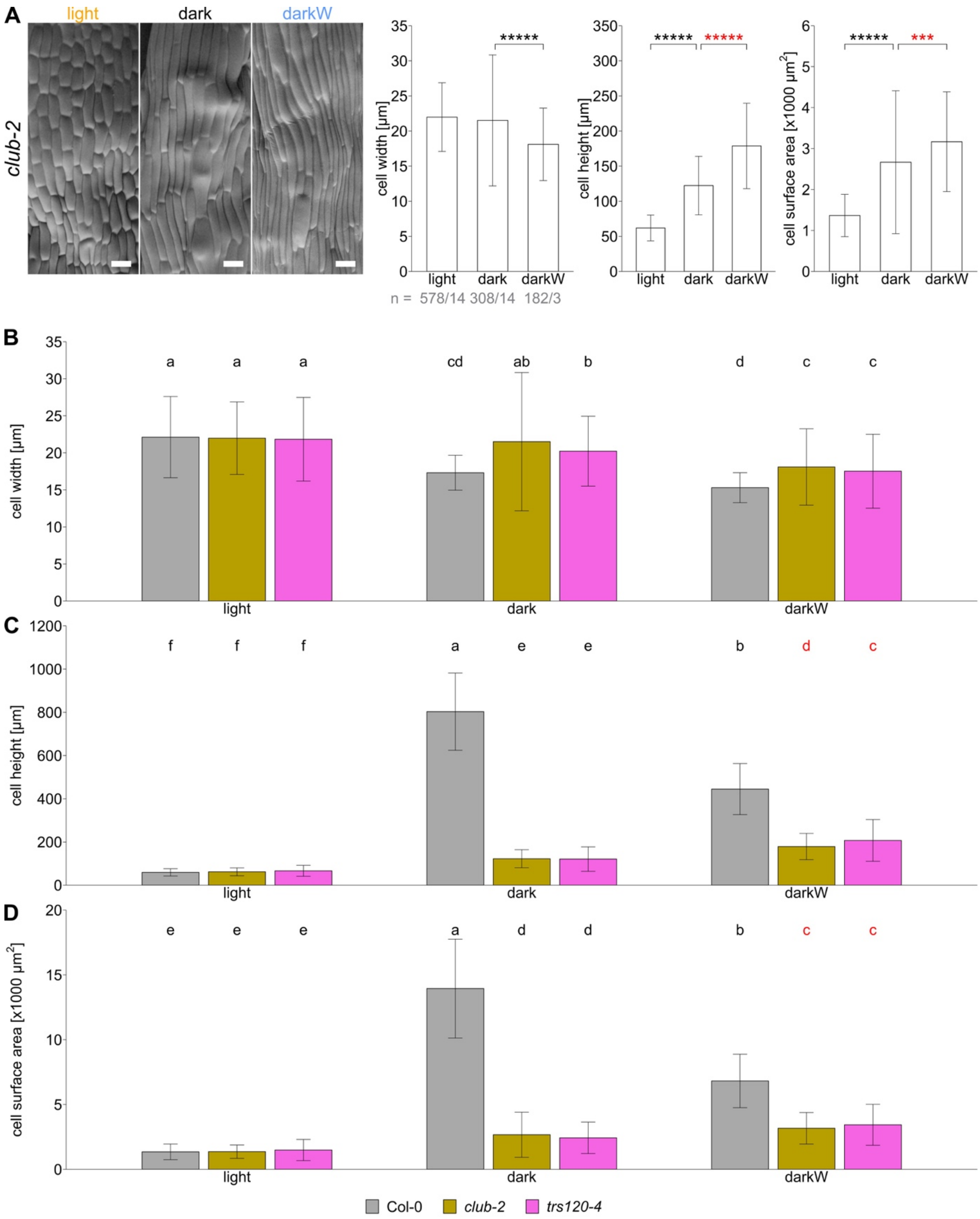


**Figure S11. Cellular hypocotyl parameters of *trappii* mutants under single and additive stress conditions.**

Seedlings were grown in the light (orange), dark (black) or dark with -0.4 MPa water stress (darkW; blue).

**A.** Scanning electron micrographs (SEM) of hypocotyls of *club-2* mutants. The cell surface area was calculated as the product of cell width and length. *club-2* mutants showed, identical to *trs120-4* (Fig. 5B), the opposite hypocotyl cell height and cell surface adaptations to dark-to-darkW conditions (red asterisks) than the wild-type control (Fig. 5A). Representative SEM images of seedlings grown under the specified conditions are shown (scale bar = 40 µm). The sample size (n) is given as the number of cells/ number of seedlings that were analyzed in grey below the graph. Shown are means ± SD. *P*-

values were computed with a two-tailed student's t-test and are represented as follows: ***: P<0.001; *****: P<0.00001.
**B-D.** Bar plots for a direct comparison of the hypocotyl B) cell width, C) cell height and D) cell surface area of *trappii* mutants and the wild-type control (Col-0) shown in Fig. 5A-B and Fig. S11A. The cellular hypocotyl parameters in the light did not significantly differ between *trappii* mutants and the wild type. *club-2* and *trs120-4* showed a compromised, but significant adjustment of the cellular parameters under single stress conditions (light to dark). Only under multiple stress (dark to darkW), *trappii* mutants failed to correctly adjust their cellular hypocotyl parameters with the opposite adjustment of the cell height and surface area compared to the wild type (red letters). Shown are means ± SD. Letters indicate statistical significance in two-way ANOVA with Tukey's post hoc test. Related to Fig. 5.

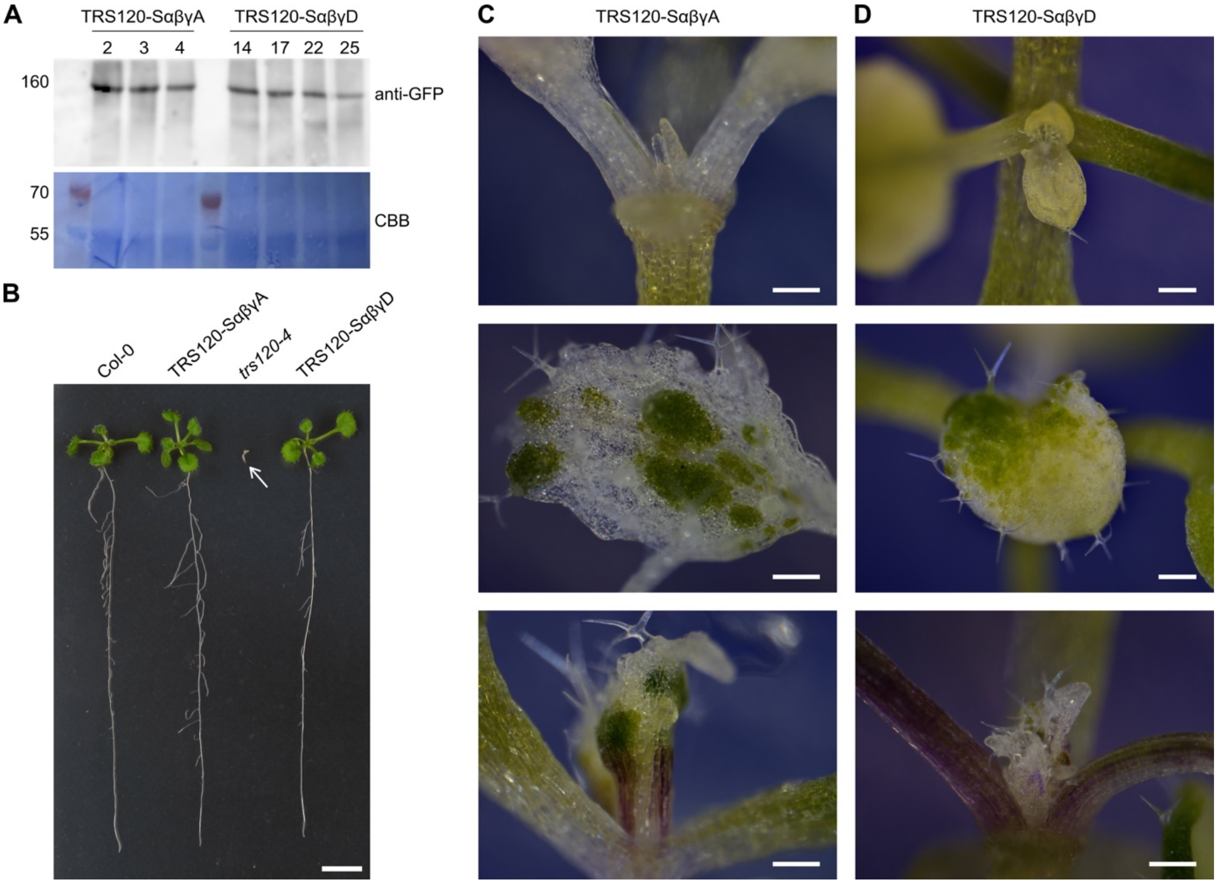


**Figure S12. Characterization of TRS120:GFP phosphovariants**.
**A.** Western blot depicting protein expression of the phosphovariants TRS120-SαβγA and TRS120-SαβγD in *trs120-4/trs120-4*. Presence of the GFP-fused phosphovariants was detected with an anti-GFP antibody (upper panel). Shown are the expression levels in different primary transformants for each variant. Loaded protein amounts are shown by the corresponding CBB staining (Coomassie stain, lower panel). Protein sizes are given in kDa on the left.
**B.** Complementation analysis. The TRS120-SαβγA and TRS120-SαβγD phosphovariants in *trs120-4/trs120-4* could rescue the null *trs120-4* seedling lethal phenotype (third seedling from the left; white arrow) and did not differ from the wild-type Col-0 control in the T1 and T2 generations. Kanamycin selection was used to select for the presence of the phosphovariant construct. Images taken 13 days after stratification. Scale bar is 1 cm.
**C-D.** Silencing of the phosphovariant constructs upon propagation beyond the T2 generation. Two-week-old seedlings were imaged using a camera-equipped binocular microscope. C) TRS120-SαβγA and D) TRS120-SαβγD seedlings grown on kanamycin had some white sectors due to loss of chlorophyll in the shoot apical meristem and first true leaves; this was presumably due to gene silencing. Scale bars represent 200 µm.

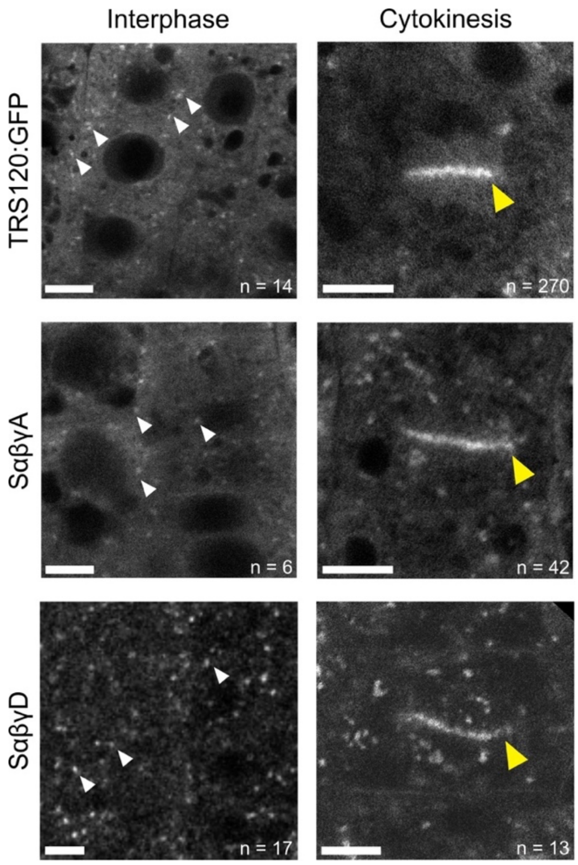


**Figure S13. Localization of TRS120:GFP phosphovariants**.
Localization patterns of TRS120:GFP phosphovariants in root tip cells of light-grown seedlings. Confocal Scanning Laser Microscopy. $P_{TRS120}$::TRS120:GFP (Rybak *et al*, 2014) and phosphovariants thereof. Scale bars represent 5 µm. Wild-type TRS120 resides in the cytosol and at the TGN, it labels endomembrane compartments (white arrowheads) and the cell plate (yellow arrowhead); the phosphovariants have a similar appearance. Sample numbers n = interphase or cytokinetic cells imaged. 44 TRS120-WT, 13 -SαβγA and 10 -SαβγD root tips were imaged.

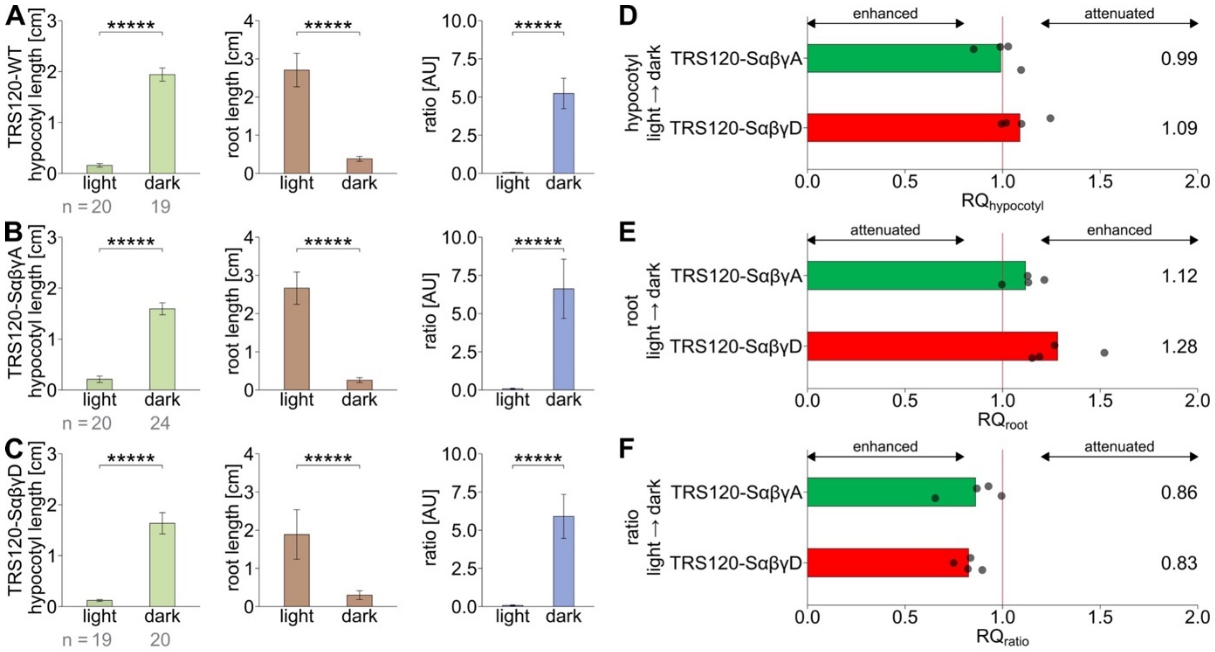


**Figure S14. The etiolation response in TRS120 phosphovariants.**

**A.** TRS120-WT ($P_{TRS120}$::TRS120:GFP in *trs120-4/trs120-4*) (Rybak *et al*, 2014).

**B.** The non-phosphorylatable TRS120 mutant TRS120-SαβγA in *trs120-4/trs120-4.*

**C.** The phosphomimetic TRS120 mutant TRS120-SαβγD in *trs120-4/trs120-4.*

Seedlings were grown on ½ MS media under light and dark conditions. All mutants showed a significant adaptation to light versus dark conditions, i.e. increased hypocotyl growth and reduced root growth leading to a higher hypocotyl/root ratio under dark conditions. For each mutant 4 biological replicates were analyzed, and the representative line based on the RQ and *P*-values is depicted here. Values are means ± SD of measured seedlings per condition, number (n) of seedlings are given in grey below the graphs. Stars indicate statistical significance with a two-tailed student's t-test (*****: P<0.00001).

**D-F.** Response quotients (RQ) of D) the hypocotyl, E) the root and F) the hypocotyl/root ratio of the non-phosphorylatable TRS120-SαβγA and the phosphomimetic TRS120-SαβγD mutants for light-to-dark adaptation. RQ response quotients are normalized to the TRS120-WT control, $P_{TRS120}$::TRS120:GFP in *trs120-4/trs120-4*. A value of 1 (vertical red line) corresponds to an identical adaptation to dark conditions as the control. TRS120-SαβγA and TRS120-SαβγD phosphovariants did not measurably differ from the control (TRS120-WT) line, with RQ values close to 1 (TRS120-SαβγA: mean $RQ_{ratio}$ = 0.86; TRS120-SαβγD: mean $RQ_{ratio}$ = 0.83). Note that due to the opposite adaptations of the hypocotyl and root from light-to-dark conditions (increase in hypocotyl, decrease in root length and therefore increased hypocotyl/root ratio), the respective thresholds for attenuated and enhanced responses are opposite. Dots represent biological replicates, mean RQ values are given on the right. Related to Fig. 6.

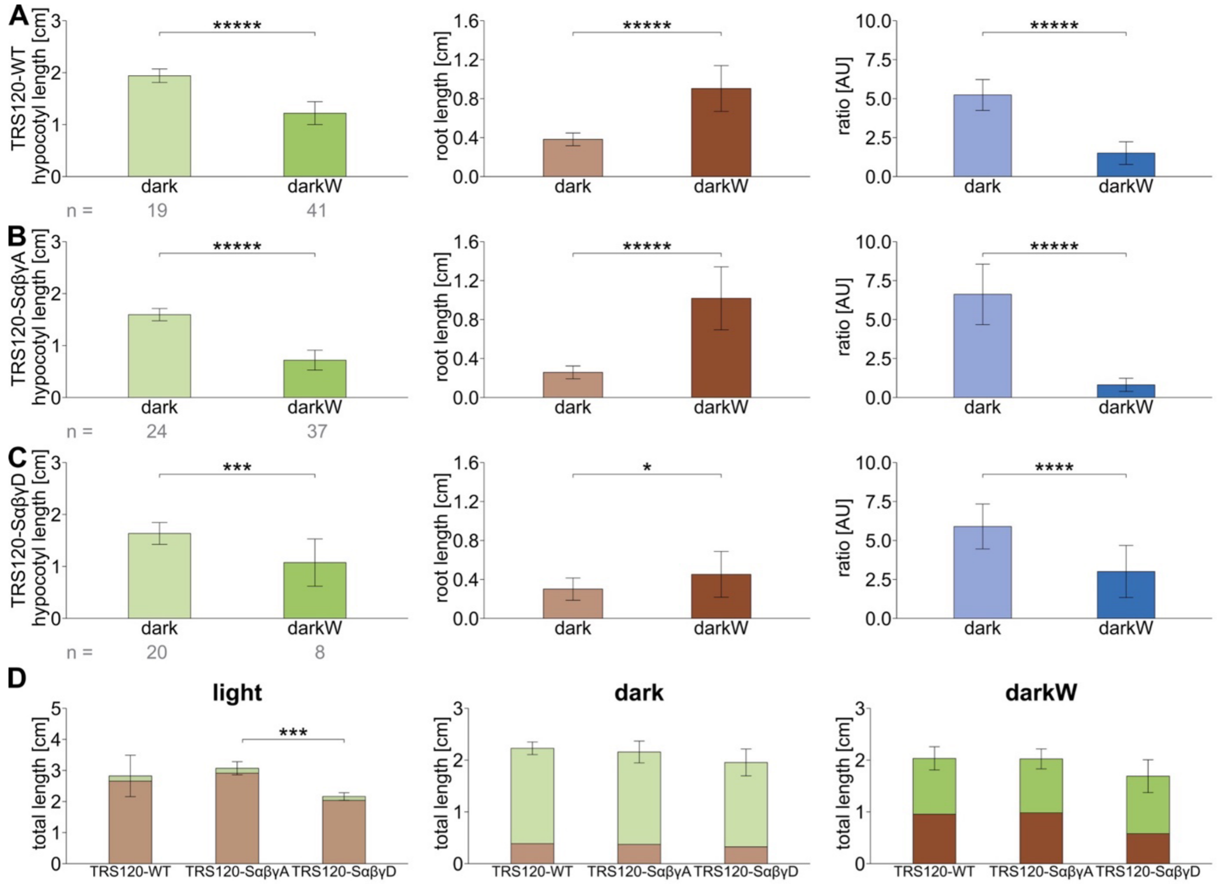


**Figure S15. Role of TRS120 phosphorylation status on adaptive growth responses under additive stress conditions.**

**A.** TRS120-WT ($P_{TRS120}$::TRS120:GFP in *trs120-4/trs120-4*) (Rybak *et al*, 2014).

**B.** The non-phosphorylatable TRS120-SαβγA in *trs120-4/trs120-4.*

**C.** The phosphomimetic TRS120-SαβγD in *trs120-4/trs120-4.*

TRS120-SαβγD mutants had attenuated root and ratio responses under darkW, whereas TRS120-SαβγA mutants showed a slightly enhanced root response.

**D.** Total seedling lengths of TRS120-WT, TRS120-SαβγA and TRS120-SαβγD. The hypocotyl length is depicted in green and the root in beige-brown shades with an increasing opacity gradient for the different conditions. The total seedling length did not vary from the TRS120-WT control, highlighting the adaptation defect of TRS120-SαβγD to darkW.

Seedlings were grown on ½ MS media under light, dark and dark with -0.4 MPa water stress (darkW) conditions. For each line, 4 biological replicates were analyzed. Values are means ± SD of measured seedlings per condition, number (n) of seedlings are given in grey below the graphs. A representative line, based on the RQ and *P*-values, is depicted for A-C. For total seedling lengths, means ± SD of 4 biological replicates are shown. Stars indicate statistical significance with a two-tailed student's t-test (*: P<0.05; **: P<0.01; ***: P<0.001; ****: P<0.0001; *****: P<0.00001). Related to Fig. 6.

**Movie S1. 3D projection of cross-kingdom structural alignment of TRAPPII.**
AlphaFold prediction (Jumper *et al*, 2021; Varadi *et al*, 2022) of *Arabidopsis thaliana* (At) AtTRS120/TRAPPC9 aligned with the open formation of the TRAPPII monomer structure without substrate in *Saccharomyces cerevisiae* (Sc) resolved with cryo-electron microscopy (*in vitro*) from Mi *et al* (2022). Alignment was performed with the align algorithm in PyMOL. Reported root-mean-square deviation (RMSD) was 16.966 Å (3775 atoms). The ScTRAPPII monomer is depicted in different shades of blue (ScCoreTRAPP with ScTCA17: blue-grey; ScTRS130: purple; ScTRS120: light blue; ScTRS65: dark blue). AtTRS120 is colored based on its sequence conservation (red: conserved sequences; orange: intermediate conservation; green: plant-specific sequences - as depicted in Fig. 2A, 2D, S4). Note that the β and γ phosphorylation sites of AtTRS120 (magenta sticks) face inwards, toward the active site chamber (including the RAB11/Rab-A GTPase binding pocket) proposed by Mi *et al* (2022) and Bagde & Fromme (2022). Related to Fig. 2, S4.

**Supplementary Table S1. Mutant lines used in this study.** a: Seedling (abbreviated “sdlg”)-lethal lines were propagated as hetero- or hemizygotes. b: Note that *club-2* is referred to as *attrs130* in Qi *et al* (2011). c: This is distinct from the hypomorphic allele later named *trs120-4* by Qi *et al* (2011).

| Allele | AGI gene identification | Polymorphism | Intron/Exon | Nature of the allele | Ecotype | Reference |
|---|---|---|---|---|---|---|
| *club-2*<br>*(attrs130*[b]*)* | At5g54440 | SALK_039353 | Intron | Null; sdlg[a] lethal | Col-0 | (Jaber *et al*, 2010) |
| *trs120-4*[c] | At5g11040 | SAIL_1285_D07 | Intron | Null; sdlg[a] lethal | Col-0 | (Thellmann *et al*, 2010) |
| *trs120i* | At5g11040 | TRS120 amiRNA | amiRNA target site in first exon | Knock-down | Col-0 | This study |
| *echidna* | At1g09330 | SAIL_163_E09 | Intron | Null; viable | Col-0 | (Gendre *et al*, 2011) |
| *bin2-3 bil1 bil2* | At4g18710<br>At2g30980<br>At1g06390 | FLAG_593C09<br>T-DNA insertion<br>T-DNA insertion | Exon<br>Exon<br>Exon | Triple null | Ws-2 | (Yan *et al*, 2009) |
| *bin2-3 bil1 bil2 trs120i* | At4g18710<br>At2g30980<br>At1g06390<br>At5g11040 | FLAG_593C09<br>T-DNA insertion<br>T-DNA insertion<br>TRS120 amiRNA | Exon<br>Exon<br>Exon<br>amiRNA target site in first exon | Triple null & knock-down | Ws-2 | This study |
| *phyA-201 phyB-5 cry1-1 cry2* | At1g09570<br>At2g18790<br>At4g08920<br>At1g04400 | EMS-induced amino acid substitution Q980STOP<br>EMS-induced amino acid substitution W552STOP<br>Fast neutrons-induced deletion<br>EMS-induced amino acid substitution W54STOP | Exon<br>Exon<br>Intron<br>Exon | Quadruple null | Ler | (Mazzella & Casal, 2001) |
| *pyr1-1 pyl1-1 pyl2-1 pyl4-1* | At4g17870<br>At5g46790<br>At2g26040<br>At2g38310 | EMS-induced amino acid substitution Q169STOP<br>SALK_054640<br>CSHL_GT2864<br>SAIL_517_C08 | Exon<br>Exon / 3’ UTR<br>Exon (3’ UTR)<br>Exon | Quadruple null | Col-0 | (Park *et al*, 2009) |
| *aba2-1* | At1g52340 | EMS-induced amino acid substitution S264N | Exon | Hypo-morph | Col-0 | (González-Guzmán *et al*, 2002) |
| *hab1-1 abi1-2 pp2ca-1* | At1g72770<br>At4g26080<br>At3g11410 | SALK_002104<br>SALK_072009<br>SALK_028132 | Intron<br>Exon (5’ UTR)<br>Exon | Triple null | Col-0 | (Rubio *et al*, 2009) |

**Supplementary Table S2. Selected brassinosteroid signaling proteins in the CLUB interactome.** Full protein name, abbreviation and AGI identifier of the depicted brassionsteroid-related proteins in the CLUB/AtTRS130 interactome in Fig. 1C and S1B.

| Abbreviation | AGI gene identification | Protein name |
|---|---|---|
| GRF6 | At5g10450 | G-BOX REGULATING FACTOR 6 |
| VIK | At1g14000 | VH1-INTERACTING KINASE |
| GF14 PHI | At1g35160 | GF14 PROTEIN PHI CHAIN |
| TOR | At1g50030 | TARGET OF RAPAMYCIN |
| 14-3-3 Omega | At1g78300 | GENERAL REGULATORY FACTOR 2 |
| GRF 10 | At1g22300 | GENERAL REGULATORY FACTOR 10 |
| ROC1 | At4g38740 | ROTAMASE CYP 1 |
| BSK8 | At5g41260 | BRASSINOSTEROID-SIGNALING KINASE 8 |
| SK11/12 | At5g26751/At3g05840 | SHAGGY-RELATED KINASE 11/12 |

**Supplementary Table S3. Thresholds for attenuated, normal and enhanced response quotients.** The thresholds for attenuated and enhanced response quotients (RQ) differ due to the opposite organ adaptations to the different conditions.

| | **dark-to-darkW** | | | **light-to-dark** | | |
|---|---|---|---|---|---|---|
| **RQ** | attenuated | normal | enhanced | attenuated | normal | enhanced |
| hypocotyl | < 0.8 | 0.8 - 1.2 | > 1.2 | > 1.2 | 0.8 - 1.2 | < 0.8 |
| root | > 1.2 | 0.8 - 1.2 | < 0.8 | < 0.8 | 0.8 - 1.2 | > 1.2 |
| ratio | < 0.8 | 0.8 - 1.2 | > 1.2 | > 1.2 | 0.8 - 1.2 | < 0.8 |

### SUPPLEMENTARY METHODS

#### Molecular techniques

Standard molecular techniques were used for subcloning (Sambrook *et al*, 1989). Phosphomutants of TRS120 were generated using a DpnI-mediated Site-Directed Mutagenesis protocol. Briefly, site-directed mutations were introduced into the template construct via polymerase chain reaction using mutagenic primers with the desired mutations (see Supplemental Table S4) and the KOD Hot Start DNA Polymerase (Novagen®) for strand extension. Subsequently, the methylated nonmutated DNA template was digested with the DpnI endonuclease (Thermo Scientific). Mutated vectors were transformed in *E. coli* DH5α for nick repair and amplification of the plasmids. After subsequent purification, the constructs were sequenced to ensure correct mutagenesis. For mutation of two or three phosphorylation sites, sequential mutagenesis was carried out using already mutated vectors as template.

For Y2H assays, mutations of phosphorylation sites were introduced into TRS120-T2 fused to GAL4-AD (pAD-GAL4). N-terminal GST fused TRS120-T2 phosphovariants (pDEST15) were used for *in vitro* kinase assays. For *in planta* experiments (confocal imaging, protein expression etc.) phosphovariants of genomic constructs with $P_{TRS120}$::TRS120:GFP in the pCAMBIA2300 plasmid were generated. The final constructs were transformed into the *Agrobacterium tumefaciens* strain GV3101 (pMP90). The phosphovariant constructs were inserted into the hemizygous null *trs120-4* background using the floral dip method (Clough & Bent, 1998). Transgenic lines were selected on ½ MS medium supplemented with 50 µg/ml kanamycin.

**Supplemental Table S4. List of sites mutated in TRS120 phosphovariants and corresponding primer sequences used for mutagenesis.**

| Phospho-variant | Amino acid substitutions | Primer sequences |
|---|---|---|
| SαA | S923A | fwd: 5'-GCC AAG GAA GAT GAT TCT GCA CCA GTA CAA GAT TCT CCA GAG-3'<br>rev: 5'-CTC TGG AGA ATC TTG TAC TGG TGC AGA ATC ATC TTC CTT GGC-3' |
| SβA | S971A, S973A, S974A, S975A | fwd: 5'-CCC TCC ACC TGG TGC CCC TGC AGC TGC TAG AAA TCC GAG CTT CTC-3'<br>rev: 5'-GAG AAG CTC GGA TTT CTA GCA GCT GCA GGG GCA CCA GGT GGA GGG-3' |
| SγA | T1163A, S1165A | fwd: 5'-GTA CTC AGA GCA CGA GCA GGA GCT GCT GCT CCA AAC GAA CCC ATC-3'<br>rev: 5'-GAT GGG TTC GTT TGG AGC AGC AGC TCC TGC TCG TGC TCT GAG TAC-3' |
| SαβA | S923A, S971A, S973A, S974A, S975A | fwd: 5'-GCC AAG GAA GAT GAT TCT GCA CCA GTA CAA GAT TCT CCA GAG-3' |

| | | |
|---|---|---|
| | | rev: 5’-CTC TGG AGA ATC TTG TAC TGG TGC AGA ATC ATC TTC CTT GGC-3’<br><br>fwd: 5’-CCC TCC ACC TGG TGC CCC TGC AGC TGC TAG AAA TCC GAG CTT CTC-3’<br><br>rev: 5’-GAG AAG CTC GGA TTT CTA GCA GCT GCA GGG GCA CCA GGT GGA GGG-3’ |
| SαγA | S923A, T1163A, S1165A | fwd: 5’-GCC AAG GAA GAT GAT TCT GCA CCA GTA CAA GAT TCT CCA GAG-3’<br><br>rev: 5’-CTC TGG AGA ATC TTG TAC TGG TGC AGA ATC ATC TTC CTT GGC-3’<br><br>fwd: 5’-GTA CTC AGA GCA CGA GCA GGA GCT GCT GCT CCA AAC GAA CCC ATC-3’<br><br>rev: 5’-GAT GGG TTC GTT TGG AGC AGC AGC TCC TGC TCG TGC TCT GAG TAC-3’ |
| SβγA | S971A, S973A, S974A, S975A, T1163A, S1165A | fwd: 5’-CCC TCC ACC TGG TGC CCC TGC AGC TGC TAG AAA TCC GAG CTT CTC-3’<br><br>rev: 5’-GAG AAG CTC GGA TTT CTA GCA GCT GCA GGG GCA CCA GGT GGA GGG-3’<br><br>fwd: 5’-GTA CTC AGA GCA CGA GCA GGA GCT GCT GCT CCA AAC GAA CCC ATC-3’<br><br>rev: 5’-GAT GGG TTC GTT TGG AGC AGC AGC TCC TGC TCG TGC TCT GAG TAC-3’ |
| SαβγA | S923A, S971A, S973A, S974A, S975A, T1163A, S1165A | fwd: 5’-GCC AAG GAA GAT GAT TCT GCA CCA GTA CAA GAT TCT CCA GAG-3’<br><br>rev: 5’-CTC TGG AGA ATC TTG TAC TGG TGC AGA ATC ATC TTC CTT GGC-3’<br><br>fwd: 5’-CCC TCC ACC TGG TGC CCC TGC AGC TGC TAG AAA TCC GAG CTT CTC-3’<br><br>rev: 5’-GAG AAG CTC GGA TTT CTA GCA GCT GCA GGG GCA CCA GGT GGA GGG-3’<br><br>fwd: 5’-GTA CTC AGA GCA CGA GCA GGA GCT GCT GCT CCA AAC GAA CCC ATC-3’<br><br>rev: 5’-GAT GGG TTC GTT TGG AGC AGC AGC TCC TGC TCG TGC TCT GAG TAC-3’ |
| SαD | S923D | fwd: 5’-GCC AAG GAA GAT GAT TCT GAC CCA GTA CAA GAT TCT CCA GAG-3’<br><br>rev: 5’-CTC TGG AGA ATC TTG TAC TGG GTC AGA ATC ATC TTC CTT GGC-3’ |

| | | |
|---|---|---|
| SβD | S971D, S973D, S974D, S975D | fwd: 5’-CCC TCC ACC TGG TGA CCC TGA CGA TGA TAG AAA TCC GAG CTT CTC-3’<br>rev: 5’-GAG AAG CTC GGA TTT CTA TCA TCG TCA GGG TCA CCA GGT GGA GGG-3’ |
| SγD | T1163D, S1165D | fwd: 5’-GTA CTC AGA GCA CGA GCA GGA GAT GCT GAT CCA AAC GAA CCC ATC-3’<br>rev: 5’-GAT GGG TTC GTT TGG ATC AGC ATC TCC TGC TCG TGC TCT GAG TAC-3’ |
| SαβD | S923D, S971D, S973D, S974D, S975D | fwd: 5’-GCC AAG GAA GAT GAT TCT GAC CCA GTA CAA GAT TCT CCA GAG-3’<br>rev: 5’-CTC TGG AGA ATC TTG TAC TGG GTC AGA ATC ATC TTC CTT GGC-3’<br>fwd: 5’-CCC TCC ACC TGG TGA CCC TGA CGA TGA TAG AAA TCC GAG CTT CTC-3’<br>rev: 5’-GAG AAG CTC GGA TTT CTA TCA TCG TCA GGG TCA CCA GGT GGA GGG-3’ |
| SαβγD | S923D, S971D, S973D, S974D, S975D, T1163D, S1165D | fwd: 5’-GCC AAG GAA GAT GAT TCT GAC CCA GTA CAA GAT TCT CCA GAG-3’<br>rev: 5’-CTC TGG AGA ATC TTG TAC TGG GTC AGA ATC ATC TTC CTT GGC-3’<br>fwd: 5’-CCC TCC ACC TGG TGA CCC TGA CGA TGA TAG AAA TCC GAG CTT CTC-3’<br>rev: 5’-GAG AAG CTC GGA TTT CTA TCA TCG TCA GGG TCA CCA GGT GGA GGG-3’<br>fwd: 5’-GTA CTC AGA GCA CGA GCA GGA GAT GCT GAT CCA AAC GAA CCC ATC-3’<br>rev: 5’-GAT GGG TTC GTT TGG ATC AGC ATC TCC TGC TCG TGC TCT GAG TAC-3’ |

**Generation of inducible *trs120* knock-down lines**

The miR319a precursor carrying the designed TRS120 amiRNA (5’-TATAACTCTTACAAGCGGCAT-3’) was synthesized as a gene strand with attached attB sites for GATEWAY® compatibility (Eurofins; see Supplemental Table S5). This TRS120 amiRNA precursor was first cloned into the pDONR207 entry vector and subsequently into the estradiol inducible pMDC7 vector (Curtis & Grossniklaus, 2003) using Gateway cloning. The final construct was introduced into the *Agrobacterium tumefaciens* strain GV3101 (pMP90) and inserted into the Col-0 wild type and the *bin2-3bil2bil1* mutant background using the floral dip method (Table S1; Clough & Bent, 1998). Transgenic lines were selected on ½ MS medium supplemented with 20 µg/ml hygromycin B.

**Supplemental Table S5. Gene strand carrying the TRS120 amiRNA precursor.** The TRS120 amiRNA sequence is highlighted in blue and the respective amiRNA* strand in orange capital letters. attB sites are designated as underlined capital letters whereas the miR319a precursor sequence is in lower case.

| |
|---|
| 5'-<br>GGGGACAAGTTTGTACAAAAAAGCAGGCTccccaaacacacgctcggacgcatattacacatgttcatac<br>acttaatactcgctgttttgaattgatgttttaggaatatatatgtagaATACCGCTTGTAACAGTTATTtcacaggtcg<br>tgatatgattcaattagcttccgactcattcatccaaataccgagtcgccaaaattcaaactagactcgttaaatgaatgaatgat<br>gcggtagacaaattggatcattgattctctttgaTATAACTCTTACAAGCGGCATtctctcttttgtattccaattttcttg<br>attaatctttcctgcacaaaaacatgcttgatccactaagtgacatatatgctgccttcgtatatatagttctggtaaaattaacatttt<br>gggtttatctttatttaaggcatcgccatggggACCCAGCTTTCTTGTACAAAGTGGTCCC<br>-3' |

#### *In vitro* kinase assays with mass-spectrometry readout - sample preparation

Every sample was reduced with 10 mM dithiothreitol (DTT) (30 min at 30°C) and alkylated with 55 mM chloracetamide (CAA) in the dark (30 min, 25°C). Afterwards, the samples were diluted four-fold in 50 mM $NH_4HCO_3$ and digested with trypsin (1 h at 30°C, Trypsin gold Mass Spectrometry Grade, Promega) in a ratio of 1:100, i.e. 10 µg protein sample were digested with 0.1 µg trypsin. After the 1 h trypsin incubation time, the same amount of trypsin was added a second time. After 16 h of incubation the enzymatic reaction was stopped with 1% formic acid (FA).

Afterwards, stage tip purification was performed. To this end, the pH of the samples was measured (pH < 3) with pH strips (MColorpHast, Merck). The in-house built C18 tips (three disks, ø 1.5 mm, C18 material, 3M Empore) were equilibrated consecutively with 250 µl 100% ACN, 250 µl elution solution (40% ACN, 0.1% FA) and 250 µl washing solution (2% ACN, 0.1% FA) at 1500 g. The sample was loaded onto the column (5 min at 500 g) and desalted with three washing steps (washing buffer: 2% ACN, 0.1% FA; 2 min at 1500 g, 250 µl). Finally, the peptides were eluted with two times 40 µl elution solution (40% ACN, 0.1% FA) for 2 min at 500 g. The solvent of all samples was completely subtracted in a centrifugal evaporator (Centrivap Cold Trap -50, Labconco, US), freshly suspended before MS measurement in washing solution (2% ACN, 0.1% FA) and ~0.25 µg of digest was injected into the mass spectrometer per measurement.

#### LC-MS/MS data acquisition for *in vitro* kinase assays and co-immunoprecipitations

Generated peptides were analyzed on a Dionex Ultimate 3000 RSLCnano system coupled to a Q-Exactive HF-X mass spectrometer (Thermo Fisher Scientific). Peptides were delivered to a trap column (ReproSil-pur C18-AQ, 5 µm, Dr. Maisch, 20 mm × 75 µm, self-packed) at a flow rate of 5 µl/min in HPLC grade water with 0.1% formic acid. After 10 min of loading, peptides were transferred to an analytical column (ReproSil Gold C18-AQ, 3 µm, Dr. Maisch, 450 mm × 75 µm, self-packed) and separated using a linear gradient (50 min for co-immunoprecipitation samples and 30 min for kinase assay samples) from 4% to 32% of solvent B (0.1% formic acid in acetonitrile and 5% (v/v) DMSO) at 300 nl/min flow rate. Both nanoLC solvents (solvent A: 0.1% formic acid in HPLC grade water and 5% (v/v) DMSO) contained 5% DMSO to boost MS intensity. The Q-Exactive HF-X mass spectrometer was operated in data dependent acquisition (DDA) and positive ionization mode. MS1 spectra (360-1300 m/z) were recorded at a resolution of 60,000 using an automatic gain control (AGC) target value of 3e6 and maximum injection time (maxIT) of 45 msec. Up to 18 peptide precursors were selected for fragmentation in case of the full proteome analyses. Only precursors with charge state 2 to 6 were selected and dynamic exclusion of 25 sec was enabled. Peptide fragmentation was performed using higher energy collision induced dissociation (HCD) and a normalized collision energy (NCE) of 26%. The precursor isolation window width was set to 1.3 m/z. MS2 resolution was 15.000 with an automatic gain control (AGC) target value of 1e5 and maximum injection time (maxIT) of 25 msec.

**Co-immunoprecipitation: LC-MS/MS data analysis**

For AtTRS120 co-immunoprecipitations, peptide identification and quantification were performed using the software MaxQuant (version 1.6.1.0) (Tyanova *et al*, 2016a; see below) with its built-in search engine Andromeda (Cox *et al*, 2011). MS raw data was searched against an Arabidopsis reference database (Araport, updated 2016-06) supplemented with common contaminants (built-in option in MaxQuant). Carbamidomethylated cysteine was used as fixed modification; variable modifications included oxidation of methionine and N-terminal protein acetylation. Trypsin/P was specified as proteolytic enzyme with up to two missed-cleavage sites. Label-free quantification (Cox *et al*, 2014), match-between-runs intensity based absolute quantification (iBAQ) and label-free quantification (LFQ) options were enabled. Precursor tolerance was set to 4.5 ppm, and fragment ion tolerance to 20 ppm. Results were adjusted to 1% false discovery rate (FDR) on peptide spectrum match (PSM) level and protein level employing a target-decoy approach using reversed protein sequences.

Perseus (Tyanova *et al*, 2016b) and Python (matplotlib (Hunter, 2007), pandas (McKinney, 2010), and seaborn (Waskom, 2021)) were used for statistical data analysis. Normalization was performed by median centering. LFQ intensities were log2-transformed. Protein groups identified only in a single sample were removed from the analysis. Missing values were replaced from a normal distribution (width = 0.3, down shift = 1.8). Protein fold-changes and their significance were computed via a two sample t-test. A fold-change of > 3 and a *P*-value < 0.05 was regarded significant.

For CLUB/AtTRS130, IP-MS data were analyzed following the method used in Kalde *et al* (2019). In short, raw MS files were loaded into the MaxQuant software (version 1.5.7.4; Cox & Mann, 2008) and searched against an Arabidopsis thaliana RefSeq database (Araport11_genes.201606.pep.fasta). Trypsin was specified as proteolytic enzyme and up to 2 missed cleavages were allowed. Protein identifications were filtered at 1% protein false-discovery rate on PSM and protein level. In downstream analysis, we used the intensity-based absolute quantification (iBAQ) (Schwanhäusser *et al*, 2011), whose values were calculated using the implemented iBAQ algorithm in the MaxQuant software. Before statistical analysis, iBAQ intensities were normalized according the iBAQ of the bait protein used in the co-immunoprecipitation (Co-IP) and missing values were replaced by a constant (1000). Then the iBAQ intensities were log2 transformed. The proteins abundance between control and Co-IP samples were compared using t-test to evaluate the statistical significance. Ratios of protein abundance between control and Co-IP were considered high if larger than 8 in log2 scale, and intermediate if in the 5-8 range (log2 scale).

**LC-MS/MS phosphopeptide analysis**

For a targeted analysis of the AtTRS120 phosphostatus in kinase assays (Fig. 3B, S5B) and in the bikinin and PPZ data sets (Fig. S6), MS1 chromatograms from selected phosphopeptides were extracted and analyzed using the Skyline daily software (MacLean *et al*, 2010). Peak integration, interferences and integration boundaries were reviewed manually for all precursors. For the bikinin and PPZ treatments, a two-tailed student's t-test was performed to determine statistical significance. For the kinase assay, raw intensities as exported from Skyline daily were plotted.

**Gene ontology (GO) enrichment analysis of CLUB:GFP interactome**

The CLUB:GFP interactome (Kalde *et al*, 2019) was used for gene ontology (GO) term enrichment analysis. To reduce the TRAPPII interactome complexity, the GO enrichment analysis of biological processes of level 0 was done separately for high confidence interactors (intensity ratio > 8) and for intermediate intensity interactors (intensity ratio > 5 and < 8) (see cutoffs Fig. 1B). The values were computed based on a comparison between observed versus random protein occurrences for each GO term. Each term is part of the hierarchical structure of GO and has defined relationships to one or more other terms in the GO enrichment. Metabolic, transcriptional and translational processes were excluded for simplification. Terms were grouped by manually annotated super-categories. Cutoffs for enriched GO terms were set at ≥ 4 for fold enrichment and ≤ 0.003 for the FDR-adjusted *P*-value.

**Structural and multiple sequence alignments**

AlphaFold predicted structures for *Arabidopsis thaliana* AtTRS120 (UniProt: Q9FY61) and CLUB/AtTRS130 (UniProt: F4K0C4) were aligned with the *Saccharomyces cerevisiae* open formation of the TRAPPII monomer structure without substrate resolved with cryo-electron microscopy (*in vitro*) (PDB: 7E2C; Mi *et al*, 2022). The alignments were done with the align algorithm in PyMOL which first performs a sequence alignment, followed by a structural superposition and then iteratively refines the alignment.

The tree of the AtSK clades was created by a multiple sequence alignment with the Clustal Omega program in UniProt using full-length protein sequences.
